# Lipid signatures and inter-cellular heterogeneity of naïve and lipopolysaccharide-stimulated human microglia-like cells

**DOI:** 10.1101/2023.03.08.531670

**Authors:** Max A. Müller, Norman Zweig, Bernhard Spengler, Maria Weinert, Sven Heiles

**Affiliations:** Institute of Inorganic and Analytical Chemistry, Analytical Chemistry, Justus Liebig University Giessen, 35392 Giessen, Germany; Department of Brain Sciences, Imperial College London, Hammersmith Hospital, London, W12 0NN, United Kingdom; Leibniz-Institut für Analytische Wissenschaften - ISAS - e.V., 44139 Dortmund, Germany; Faculty of Chemistry, University of Duisburg-Essen, 45141 Essen, Germany

## Abstract

Microglia are non-neuronal cells, which are residing in the central nervous system and are known to play an important role in health and disease. We investigated lipidomic phenotypes of human naïve and stimulated microglia-like cells by atmospheric-pressure scanning microprobe matrix-assisted laser desorption/ionization mass spectrometry imaging (AP-SMALDI MSI). With lateral resolutions between 5 μm and 1.5 μm, we were able to chart lipid compositions of individual cells, enabling to differentiate cell lines and stimulation conditions. This allowed us to reveal local lipid heterogeneities in naïve and lipopolysaccharide (LPS)-stimulated cells. We were able to identify individual cells with elevated triglyceride (TG) levels and could show that the number of these TG-enriched cells increased with LPS stimulation as a hallmark for a proinflammatory phenotype. Additionally, observed local abundance alterations of specific phosphatidylinositols (PIs) indicate a cell specific regulation of the PI metabolism.

## Introduction

Microglia are non-neuronal cells of the central nervous system,^1^ that play important roles in brain development, maintenance,^2^ and homeostasis.^3,4^ They are the immune cells of the brain^5^ and have been implicated in several neurodegenerative diseases, including Alzheimer’s disease, Parkinson’s disease, and multiple sclerosis.^6–10^ Microglia respond to a changing micro-environment morphologically and through their protein expression pattern.^11–14^ Both characteristics have been used historically to crudely define microglia phenotypes. Recent advances in single-cell and single-nucleus transcriptomics have expanded our knowledge of microglia heterogeneity across brain region, age, and disease states.^15,13^ Thereby, it is clear now that multiple subtypes coexist within a single region of the adult brain.^16^ It is still a matter of debate, however, how these different subsets of microglia contribute to homeostasis and disease.

Another access to cell function is to directly probe downstream products such as metabolites and their make-up by mass spectrometry, for which lipids are particularly attractive bioanalytes. Besides making up most of the cell membrane, lipids fulfill bioactive functions such as signaling^17^ and energy storage,^18^ and are both directly affected by and are themselves affecting cell states.^19–21^ Mass spectrometry imaging (MSI) further enables the spatial discrimination of phenotypes both in tissue and in vitro, based on lipidomic signatures.^19,22,23^ Although lipidomic analyses of microglia cell lines have already been performed,^24,25^ the approach or spatial resolution in previous studies did not allow for interrogation of single-cell heterogeneity.

Here, we used human induced-pluripotent stem cell (iPSC)-derived microglia-like cells (MGL) to investigate the heterogeneity between and within cell lines by applying atmospheric-pressure scanning microprobe matrix-assisted laser desorption/ionization (AP-SMALDI) MSI.^26,27^ Technical and preparative methods were optimized to achieve a pixel size of 1.5 μm, allowing for single-cell analysis. This enabled us to distinguish between otherwise isogenic cell lines, indicating a high sensitivity for detecting different subpopulations. We used AP-SMALDI MSI to identify individual molecular species, involved in the well-established upregulation of triglycerides in MGL, classically activated with the proinflammatory stimulus lipopolysaccharide (LPS). Finally, our single-cell analysis revealed for the first time MGL heterogeneity based on lipid classes and individual lipid species, despite a homogenous microglia marker expression profile.

## Experimental

### Cell cultures

Microglia-like cells (MGL) were differentiated from human induced pluripotent stem cells (iPSCs) via primitive macrophage precursors (pMacPre) according to Haenseler *et al*.^28^ with small modifications. Of note, in this protocol microglia-like identity is induced through the use of brain-specific microglia growth factor IL-34 as opposed to peripherally expressed macrophage growth factor M-CSF during final differentiation. In brief, iPSC were grown in OXE8 medium^29^ on Geltrex (Gibco, Thermo Fisher Scientific, Altrincham, UK) with daily medium change and EDTA lifting at approximately 80% confluence for standard culture. For embryoid body production, 3·10^6^ iPSC were lifted with TrypLE Express (Gibco) and plated in one well of a 24-well Aggrewell 800 plate (STEMCELL Technologies, Cambridge, UK) in 2 ml EB medium (OXE8 supplemented with 20 ng/mL SCF (Miltenyi Biotec, Woking, UK), 50 ng/mL BMP4 (Gibco), and 50 ng/mL VEGF (PeproTech, London, UK)) for induction of mesoderm. Embryoid bodies were incubated for one week in EB medium with daily exchange of 75% medium, and then transferred into 2x T175 flasks containing FM7a factory medium^29^ for factory maturation. For 4 weeks, factories were fed weekly with addition of 10-15 mL FM7a. From week 4 to 6, the presence of pMacPre cells in the supernatant was assessed by flow cytometry using antibodies against CD45, CD14 (both Immunotools, Friesoythe, Germany), and CD11b (Biolegend, London, UK). Flasks that produced >95% triple-positive cells were subsequently used for weekly cell harvest. pMacPre were harvested from the factory supernatant, plated directly at 25000 cells/cm2 on glass slides, and finally differentiated into MGL for 7 days in MIC10 medium (SILAC Advanced DMEM/F12, 2mM GlutaMax (both Gibco), 10 mM glucose, 0.5 mM L-lysine, 0.7 mM L-arginine, 0.00075% phenol red (all Sigma Aldrich, Gillingham, UK), 100 ng/mL IL-34 (PeproTech), 10 ng/mL GM-CSF (Gibco)). An equal volume of MIC10 was added after 3 days in culture. The five cell lines used for this study (OX1, B8, D9, C12, E2) are clonal expansions of the parental line SFC841-03-01 (OX1; StemBANCC). For proinflammatory activation, MGL were exposed to 100 ng/mL lipopolysaccharide (LPS; Sigma Aldrich)^30^ in MIC10 for 24h. For MSI sample preparation, medium was removed, cells were washed once in phosphate-buffered saline (PBS; Sigma Aldrich) and either snap-frozen in liquid nitrogen for storage at −80°C or fixed in 4% paraformaldehyde (PFA; Thermo Fisher Scientific) for 5 min at room temperature before washing and storage in PBS at 4°C.

### Flow cytometry

Freshly harvested 5×10^5^ pMacPre were incubated for 30 min at 4°C with primary antibodies CD45-FITC (Immunotools, 21270453, 1:20), CD14-PE (Immunotools, 21620144, 1:20), and CD11b-APC (Biolegend, 301309, 1:20) or CD11c-FITC (Immunotools, 21487113, 1:20) in 50 μl FACS buffer (1% FBS, 10 μg/ml human IgG in PBS), spun at 500xg for 5 min and washed 3x with PBS. Fluorescence was acquired on a FACSCalibur (BD Biosciences).

### Microscopy

Microscopic images were produced with a digital microscope (VHX 5000, Keyence GmbH, Neu Isenburg, Germany) after washing and prior to matrix application.

### Fluorescence microscopy

MGL were grown in 96-well glass bottom imaging plates, fixed for 10 min with 4% PFA (Alfa Aesar, Heysham, UK), and washed 3x with PBS (Sigma, Gillingham, UK). For visualisation of lipid droplets, cells were stained with 0.5 μg/mL Bodipy™ 493/503 and 1 μg/mL DAPI (both Thermo Fisher Scientific, Paisley, UK) diluted in PBS for 30 min at room temperature and washed 3x with PBS. For immune fluorescence, cells were incubated over night with primary antibody Iba-1 (Abcam, ab5076, 1:250) or PU.1 (Cell Signaling, 2258, 1:250) in staining buffer (5% bovine serum albumin, 0.2% TritonX-100 in PBS), washed 3x with wash buffer (PBS/0.1% TritonX-100), incubated for 2 h with secondary antibody anti-goat-AF488 (Thermo Fisher, A-11055, 1:1000) or anti-rabbit-AF488 (Thermo Fisher, A-21206, 1:1000) and 1 μg/mL DAPI in wash buffer. MGL were washed 3x with PBS and images were acquired on a Leica SP8 confocal microscope (Leica Microsystems, Wetzlar, Germany). In our experiments, staining was only possible on samples not measured by AP-SMALDI MSI, due to the destructive nature of the method at high lateral resolution.

### AP-SMALDI MSI

Matrix was applied using an ultra-fine pneumatic spraying system (SMALDIPrep, TransMIT GmbH). For measurements in positive-ion mode, 80 μL of a solution of 60 mg dihydroxybenzoic acid (DHB; Merck, Darmstadt, Germany) in a mixture of 999 μL deionized water, 999 μL acetone and 2 μL pure trifluoroacetic acid (TFA; Merck) were applied to the samples. For measurements in negative-ion mode, 9-aminoacridine (9-AA; TCI Deutschland GmbH, Eschborn, Germany) was used. Samples for statistical analysis were covered with 300 μL of a solution of 10 mg/mL 9-AA in 70:30 ethanol-water. Samples intended for high-resolution AP-SMALDI MSI experiments were spray-coated with an optimized protocol using 160 μL of a solution of 7 mg/mL 9-AA in 60:40 acetone-water (Figure S1).

AP-SMALDI MSI was performed with a custom-built ion source based on an AP-SMALDI5 AF system (TransMIT GmbH, Giessen, Germany). The ion source is equipped with a specialized focusing objective lens capable to produce ablation spots with an average diameter of 1.1 μm (Figure S2) in reflective geometry while retaining sufficient ion signal. The distance between MS inlet and laser focal point was carefully optimized to yield a Gaussian ablation profile and maximize ion transmission. A distance of 4 mm was found to be optimal to have a preferably short distance for ions to cover while minimizing reflection and cut-off of the laser beam by the inlet capillary, which would result in greatly reduced beam quality. The ion source was coupled to a Q Exactive (Thermo Fisher Scientific, Bremen, Germany) orbital trapping mass spectrometer. All measurements were performed with a mass resolution of 140,000 at *m/z* 200 and a high voltage of 2.5 kV was applied between sample stage and inlet capillary. Laser energy was adjusted in order to produce ablation spots smaller than the pixel size chosen for each experiment (Figure S3). Laser energy settings were fixed for a given set of parameters such as matrix used and pixel size.

### Data analysis

Data evaluation was performed using the Mirion^31^ software to create MS images normalized to total ion charge^32^ and to export data for statistical analysis. The latter was performed using Perseus software package.^33^ Results of mass spectrometric data acquisition, based on signals averaged over the corresponding measurement area, were filtered for differences between cell lines in naïve or LPS-stimulated state, respectively, by an Analysis of Variance (ANOVA)-based multiple-sample test followed by a principal component analysis (PCA). Additionally, cell-line specific differences between naïve and LPS-stimulated cells were investigated following the same workflow. Lists of signals identified as statistically different by the statistical approach, as well as corresponding lipid annotations based on exact mass measurements and individual parameter settings for every statistical comparison can be found in the Tables S1 – S12. Cluster analysis was performed using t-SNE, PCA and k-means algorithms in Matlab (The Mathworks Inc., Natick, Massachusetts, USA). Violin plots were created using a Matlab script by Bechthold.^34^ Signal annotations were performed with the help of Metaspace^35^ using LIPID MAPS^36^ and SwissLipids^37^ databases.

## Results and Discussion

### High-lateral-resolution AP-SMALDI MSI experiments of microglia cells

We commenced by studying the impact of sample preparation on lipidome coverage, lipid distributions, and cell morphology. For this purpose, different washing and fixation workflows were tested as summarized in Figure S4 and S5 and summarized in the methods section of the supporting information. While fixation of cells with PFA best preserved the morphology (Figure 1a,b,f,g), signal intensities and thereby the number of individual lipid signals were higher for snap-frozen cells (Figure 1c,d,h,i). A large fraction of the most intense lipid-associated signals was not affected by sample fixation, indicating that PFA does not induce major changes of the lipidome (Figure 1e, j). The choice of solvents for matrix deposition also impacted lipid distribution patterns. Use of ethanol in solvent mixtures for 9-aminoacridine (9-AA), for example, resulted in significant lipid delocalization (Figure S6), whereas dihydroxybenzoic acid (DHB) and 9-AA matrices sprayed from acetone-water solutions yielded lipid distributions coinciding with cell locations as determined by optical microscopy (Figure S7, S8 and S9).

**Figure 1:**
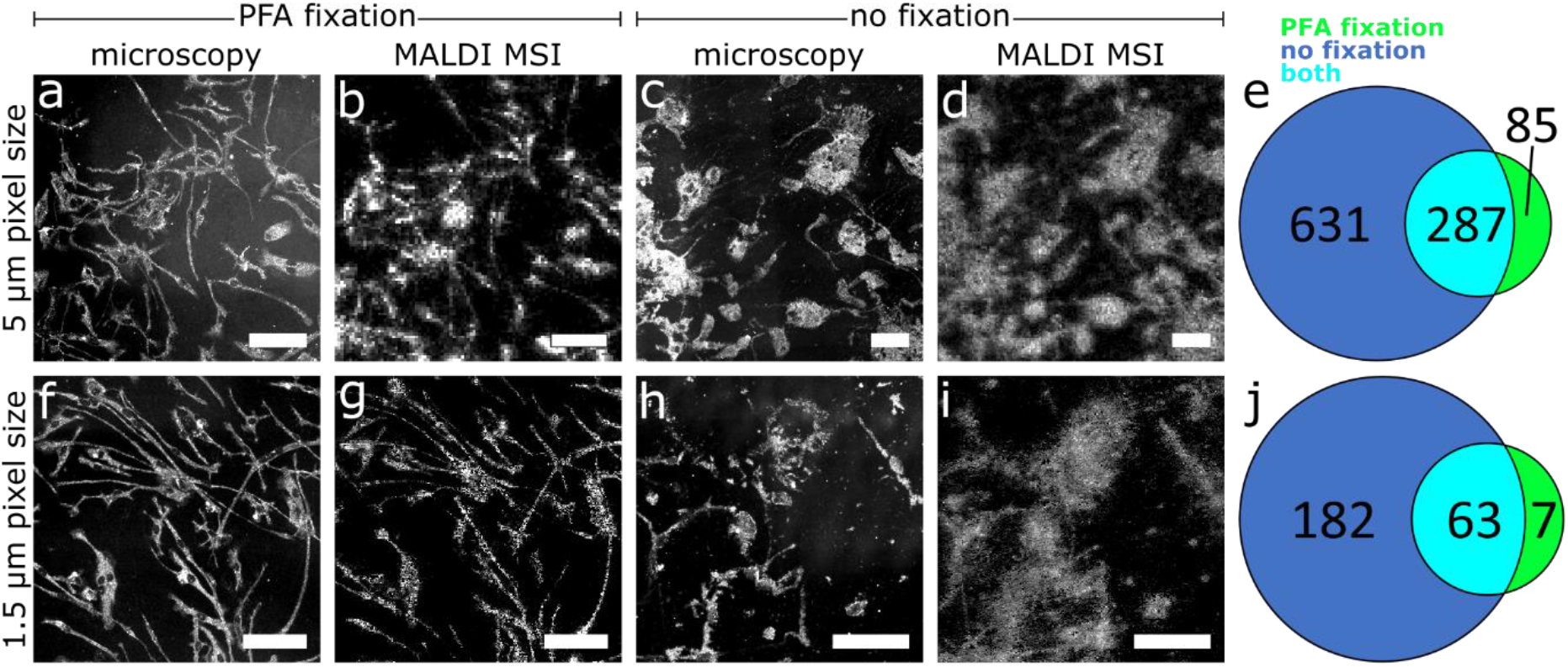
The effect of sample preparation and laser spot size on the detection of lipids from iPSC-derived microglia-like cells. a,c,f,h: Microscopic images. b,d,g,i: AP-SMALDI MS images of PC 34:1 (m/z 896.6012) at 5 or 1.5 μm laser spot size. b: LPS-stimulated MGL from the B8 cell line, 100×100 pixels, 5 μm pixel size (PFA fixation). d: LPS-stimulated MGL from the D9 cell line, 150×150 pixels, 5 μm pixel size (no fixation). g: Naïve MGL from the B8 cell line, 300×300 pixels, 1.5 μm pixel size (PFA fixation). i: Naïve MGL from the E2 cell line, 250×250 pixels, 1.5 μm pixel size (no fixation). e,j: Venn diagrams of individual lipid signals annotated to cells shown in b,d,g, and i, respectively, with fixation (green), without fixation (blue) or with both methods (cyan) for 5 μm (e) and 1.5 μm (j) measurements. Scale bars: 100 μm.

For fixed and snap frozen MGL, the number of annotations based on accurate mass measurements are reported in Figure 1e and j, and selected MS images are shown in Figure 1b, d, g and i. Mainly phosphatidylcholine (PC), phosphatidylethanolamine (PE), phosphatidic acid (PA) and triacylglyceride (TG) lipids were detected in positive-ion mode as [M+H]^+^, [M+Na]^+^, [M+K]^+^ or matrix adducts, e.g. [M + DHB – H2O + H]^+^, ions. In negative-ion mode, [M-H]^-^ ions of phosphatidylinositols (PIs) dominated the spectra.

We further tested the effect of the lateral resolution (ablation spot size) on lipid detection from MGL. Among the most intense signals, no major change in the list of detected species with pixel size was observed in the mass spectra (Figure S10). However, less intense signals gradually fell under the detection threshold with decreasing pixel size due to the reduced laser spot area. For example, at 5 μm laser focus diameter, 918 individual lipid-associated signals were observed in at least 1% of all pixels, whereas at 1.5 μm laser focus diameter, only 245 individual lipid-associated signals were detected under otherwise same conditions for snap-frozen cells.

Most of our lipid annotations at 5 μm and 1.5 μm pixel size are in line with reports by Fitzner and co-workers for lipid extracts of mouse microglia using nano-electrospray ionization following liquid chromatographic separation, one of the most commonly used methods for high coverage lipid identification, yielding the most comprehensive database on the microglia lipidome in literature.^38^ In our 5 μm and 1.5 μm measurements, we were able to detect signals that correspond to 94% and 92% of the annotations presented in the work of Fitzner and co-workers, respectively. The 6% and 8% of signals that could not be reproduced, primarily corresponded to lysophospholipids. We also detected additional lipid-associated signals that were not covered by Fitzner and co-workers. This indicates that our AP-SMALDI MSI workflow is able to sample a large fraction of all glycero-, sphingo-as well as glycerophospholipids. Annotation numbers are reduced at the higher lateral resolution of 1.5 μm but individual and overlapping cells are better resolved. Especially acquisition of the fine branched cell protrusions benefits from high lateral resolution and PFA-fixation (Figure 1 and Figure S11). At the same time, ion intensity remained high enough in our measurements to facilitate a high pixel coverage of up to 96%, 91% or 75% (Figure S12) on MGL cells at 5 μm, 2 μm or 1.5 μm pixel size, respectively, minimizing the occurrence of blank pixels. Therefore, we employed 5 μm lateral resolution to compare lipid signatures between samples and switched to higher lateral resolution to study the spatial distribution of selected lipid classes or individual lipid ion signals within samples.

### Differentiation between cell lines and activation profile

Next, we investigated the ability to discern inter-cell-line heterogeneity of naïve and LPS-stimulated MGL based on lipid signatures recorded in positive- and negative-ion mode at 5 μm resolution. Three biological replicates cultured in separate wells on the same glass slide, each containing a multitude of individual cells, where investigated per cell line and stimulation status, respectively. When comparing AP-SMALDI MSI results of different cell lines or of naïve and LPS-stimulated cells of the same cell line, some signals were only detected in one group, while others had a similar abundance in all observed groups (Figure 2e,f and Figure S13). Data is averaged over all cells measured per sample and biological triplicates were investigated. Principal component analysis (PCA) groups data of biological triplicates of identical cell lines (Figure 2a,b and Figure S14) and separates them from other cell lines based on the 1^st^ and 2^nd^ principal component. Naïve and LPS-stimulated states of the same cell line (Figure 2c and Figure S14) are also separated from each other, while biological triplicates group together. This allows to distinguish cell lines or activation status based on AP-SMALDI MSI results.

**Figure 2:**
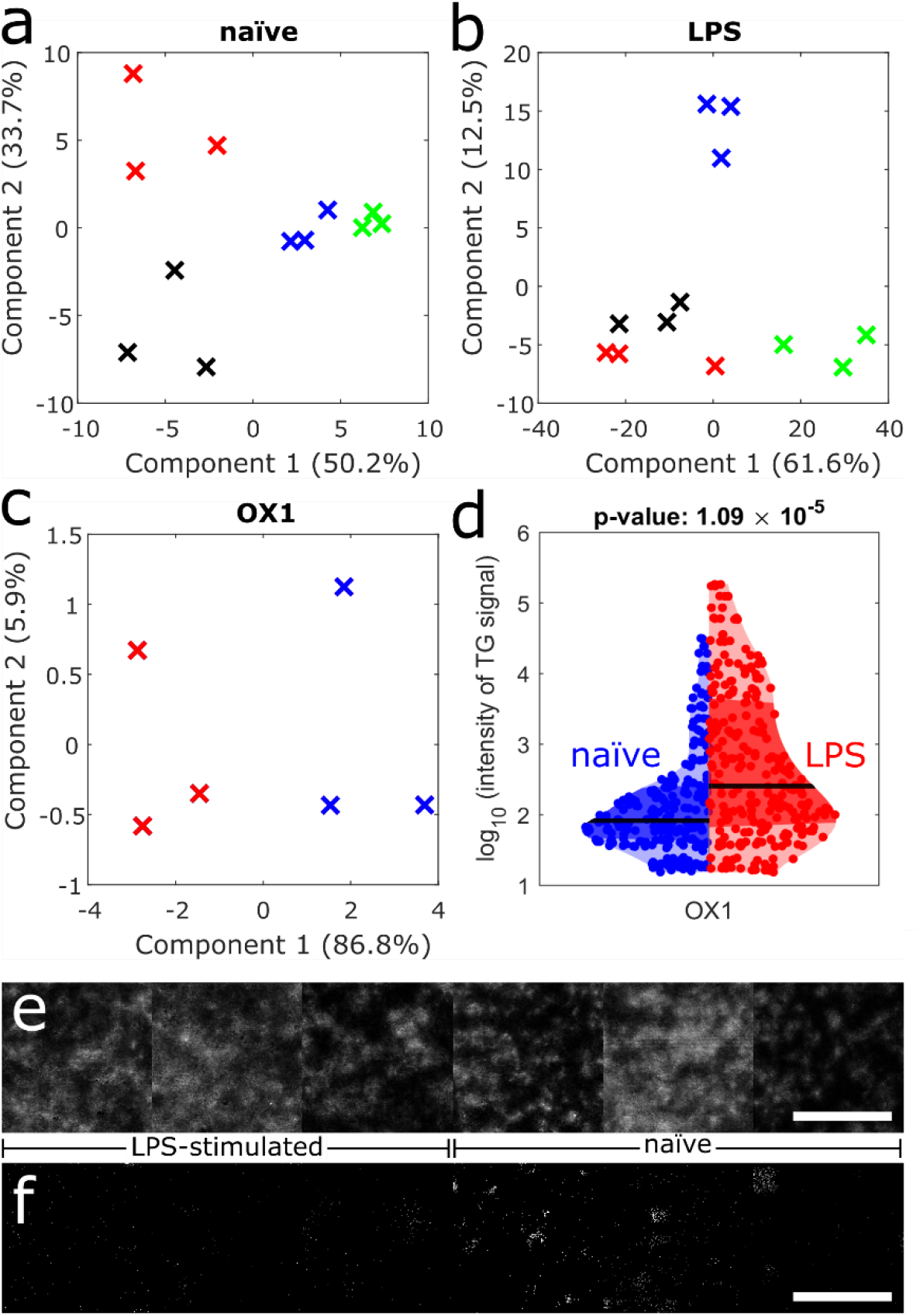
Cell-line heterogeneity assessed by AP-SMALDI MSI. a-c: PCA of AP-SMALDI MSI results in positive-ion mode using an ANOVA-based multiple sample test. Three biological replicates of cell cultures grown in separate wells on the same glass slide, each containing a multitude of individual cells, were used for analysis and mass spectrometric signal intensities were averaged over 22,500 mass spectra. Significance of signals was evaluated using p-values of <0.01. a: Cell lines OX1 (red), D9 (green), C12 (blue) and E2 (black) in naïve state. b: Cell lines OX1 (red), D9 (green), C12 (blue) and E2 (black) in LPS-stimulated state. c: Naïve (blue) and LPS-stimulated (red) cells of the OX1 cell line. d: Violin plot of 252 TG-associated signal intensities at 5 μm pixel size for naïve (blue) and LPS-stimulated (red) OX1 cells averaged over all cells of the 3 biological replicates of the cell line. Black line indicates median. p-Value calculated by double-sided t-test. e: AP-SMALDI MS image of PC 34:1 (m/z 896.6140) of biological triplicate cell cultures grown in separated wells on the same slide of the OX1 cell line in naïve and LPS-stimulated state, respectively, at 5 μm pixel size, showing similar distributions for both conditions. f: AP-SMALDI MS image of DG O-38:8 (m/z 661.4593) of biological triplicate cell cultures grown in separated wells on the same slide of the OX1 cell line in naïve and LPS-stimulated state, respectively, at 5 μm pixel size, showing enrichment in LPS-stimulated cells. Scale bars: 500 μm.

Especially LPS stimulation influences lipidomic profiles of cells (Figure 2c and Figure S14). In the PCA plot, the separation along the 1st principal component alone (making up between 78.2% to 91.4% of total variance in positive-ion mode and 91.0% to 95.4% of total variance in negative-ion mode) allows for differentiation of the stimulation status. A major contribution to this separation relates to TG species. When comparing the averaged intensity per cell line and stimulation status of all signals annotated as TGs by Metaspace^35^ using the LIPIDMAPS database,^36^ TG intensities in LPS-stimulated MGL are significantly increased (p-values of 1.09 · 10^-5^, 6.95 · 10^-6^ and 6.73 · 10^-4^ for OX1, C12 and E2 cell lines, respectively) over naïve cells (Figure 2d and Figure S15), consistent with earlier reports regarding macrophage cell lines.^39,40^ Further, LPS-stimulated cells showed a higher number of individual TG-associated signals (Figure S16 and S17), with an average increase of 8.4, 2.4, and 4.4-fold relative to naïve cells in OX1, C12, and E2 lines, respectively.

Only the D9 cell line showed an opposing behavior with the number of TGs and the mean intensity of TG signals virtually unchanged in naïve compared to LPS-stimulated cells (Figure S15, S16 and S17), which is driven by an increased number of TG signals in the naïve state, rather than a missing reaction to LPS stimulation (Figure S15). Although the MGL of D9 expressed the conventional myeloid markers CD14 and CD11b, indicating successful differentiation, cell yield and time of productivity of the D9 cell factories was consistently lower compared to all other cell lines (Figure S18). The lack of MGL precursor proliferation and increased TG signal intensities is consistent with a premature senescence of D9 mesodermal factories.^41^

TGs are known to be the main component of lipid droplets (LD) in cells.^42^ In microglia, reports indicate that lipid droplets play a major role during metabolic changes such as glucose deprivation^43^ and that their accumulation is associated with a pro-inflammatory phenotype^20^ and inflammatory response mechanisms, such as those elicited by LPS.^44,30^ AP-SMALDI MSI allows to elucidate the composition of these TG accumulations, and a list of TG annotations is given in Figure S16.

### Lipid droplet distribution differs locally between individual cells

We investigated the capability of AP-SMALDI MSI to identify heterogeneous compound distributions within cell cultures of the same line. Representative results with 5 μm, 2 μm, and 1.5 μm pixel size of PFA-fixed B8 cells are shown in Figure 3 a, c and e and compared to corresponding microscopic results in Figure 3b, d and f. With increasing pixel resolution, cellular features such as cell bodies and especially protrusions were readily resolved by visualizing major cell membrane components such as PC 34:1 (in Figure 3, green). The diameter of the protrusions imaged by AP-SMALDI MSI with 1.5 μm pixel size was ≈5 μm, consistent with the diameter of the same features in the optical images (Figure S11). This indicates that lipids do not disperse in the matrix upon our sample preparation.

**Figure 3:**
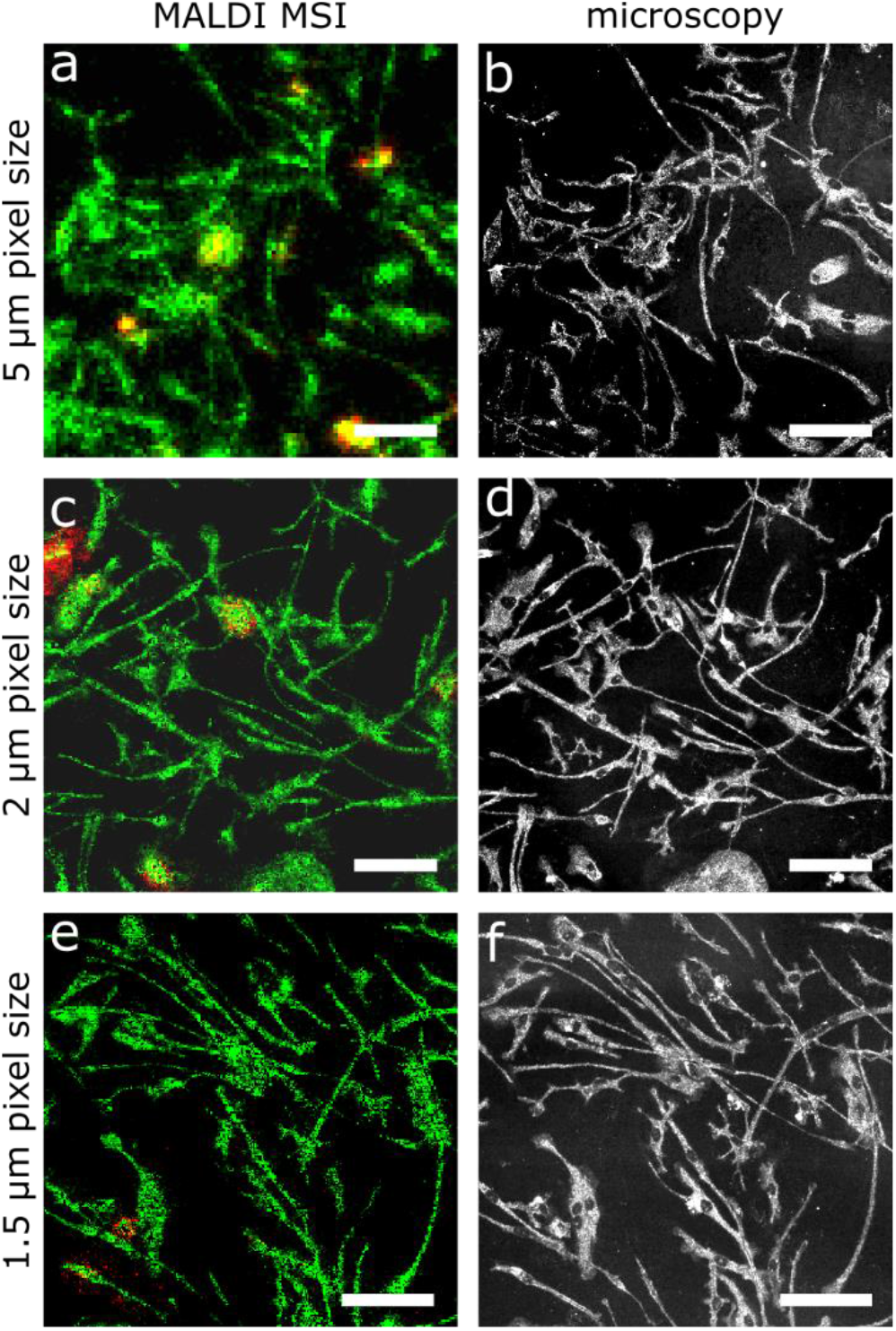
Increasing lateral resolution enables the localization of discrete subcellular features by AP-SMALDI MSI. a: LPS-stimulated MGL cells of the B8 cell line, 100×100 pixels, 5 μm pixel size. c: Naïve MGL cells of the B8 cell line, 250×250 pixels, 2 μm pixel size. e: Naïve MGL cells from the B8 cell line, 300×300 pixels, 1.5 μm pixel size. b,d,f: Corresponding microscopic images of the measured sample regions. a,c,e: Color-coding: red: TG 52:2 (m/z 881.7568), green: PC 34:1 (m/z 896.6140). Scale bars: 100 μm.

In addition to signals that colocalize with all cells, many signals were confined to individual cells and were either absent or decreased in intensity in other cells of the same culture (red color representing TG 52:2 associated ions in Figure 3, Figure S19 & S20). This was observed in all cell lines for all lateral resolutions down to 1.5 μm, including LPS-stimulated B8 cells shown in Figure 4. When employing t-distributed stochastic neighbor embedding (t-SNE) on mass spectrometric data from all on-cell pixels of a measurement, two clusters were identified, colored in green and red in Figure 4a. A third cluster was identified, but corresponding mass spectra were virtually identical with those of the main cluster (green), showing >95% congruence for all signals with an intensity >2%, and corresponding pixels seem to be randomly distributed (Figure S21, blue pixels). These data points were thus included in the main cluster (green, Figure 4a). Differences between mass spectra of pixels of the two identified groups (red and green cluster, Figure 4a) were found especially in the mass range above *m/z* 800, contributed by high-intensity signals, while spectra were rather similar in the lower mass range where only low-intensity signals were dissimilar. (Figure 4b). The majority of signals in the range *m/z* 800 – 1000 were assigned to TGs, with some being elevated in signal intensities for the red cluster while being reduced or absent for the green cluster. When visualizing the spatial distributions of the two clusters, pixels of the less abundant red cluster were highly localized to individual cells (Figure 4d). Comparison of the spatial distribution of the red cluster (Figure 4d) with that of the highly variable signal corresponding to TG 52:2 (Figure 4c) revealed that both images virtually overlap. Analysis of other MS images yielded similar results (Figure S22). This demonstrates that our methodology is able to locally resolve regions with elevated TG-content, most likely areas of lipid droplet accumulation. They further serve as a proof of concept that our approach is suitable to study subtype heterogeneity among cells.

**Figure 4:**
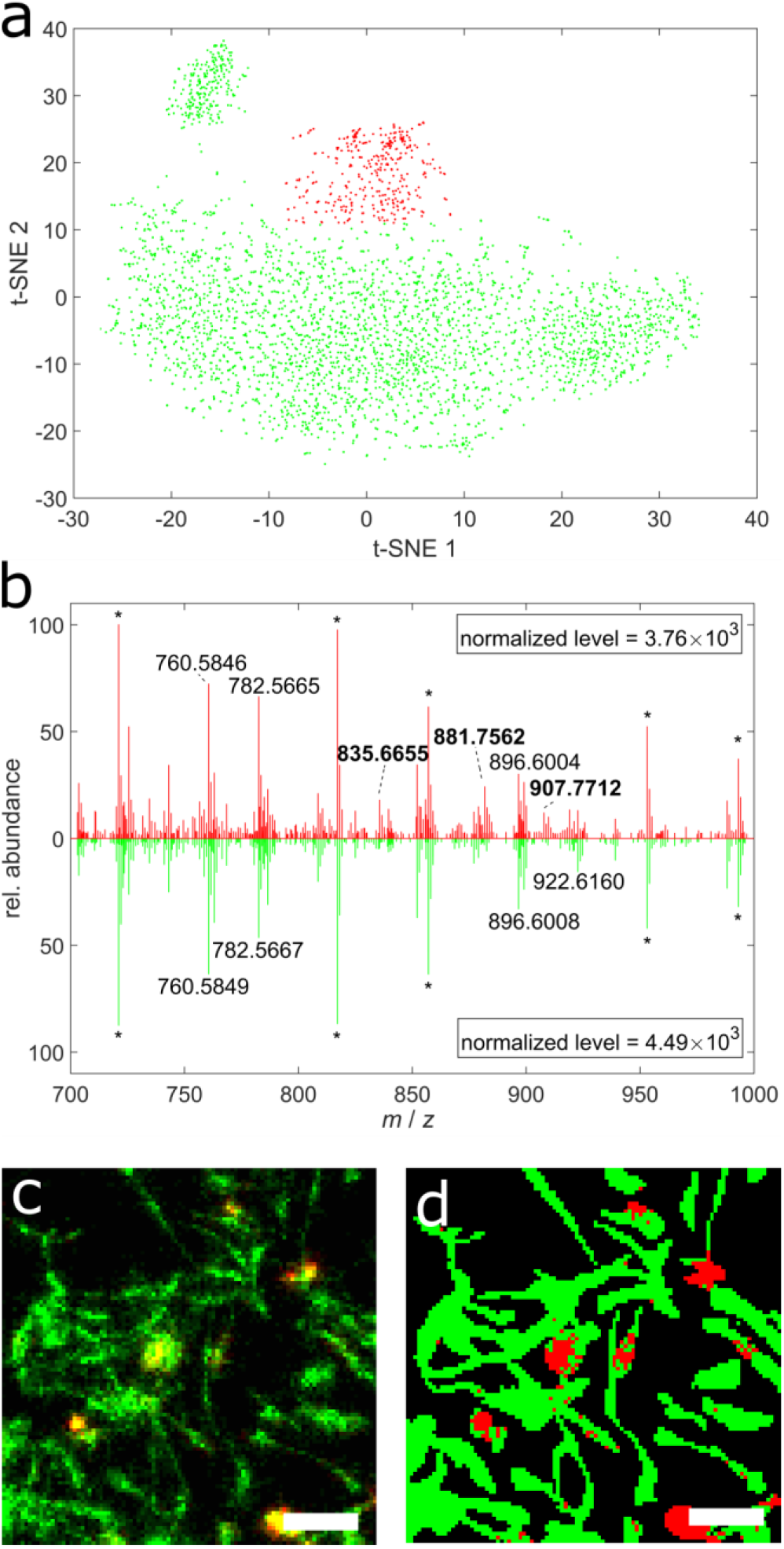
AP-SMALDI MSI and statistical analysis for identification of LPS-stimulated B8 cells with increased TG production. a: Pixel-wise t-SNE analysis of mass spectrometric data (m/z 800-1000), where each of the 6511 datapoint relates to an individual mass spectrum. Off-cell signals were excluded from analysis. b: Example of single-pixel mass spectra corresponding to the red and green cluster as identified by pixel-based t-SNE analysis in a. Stars indicate matrix-related signals. c: AP-SMALDI MSI image with 100×100 pixels at 5 μm pixel size of LPS-stimulated PFA-fixed B8 cells. Color-coding: red: TG 52:2 (m/z 881.7568), green: PC 36:2 (m/z 808.5826). d: Pixel-wise classification based on t-SNE analysis of the mass spectrometric data with automated color-coding referring to clusters identified in a. Additional information in Figure S21. Scale bars: 100 μm.

When comparing MS images of naïve and LPS-stimulated MGL, we observed that the number of cells increased with elevated TG levels increased with activation (Figure S19 & S20), consistent with the finding that LPS stimulation augmented triglyceride levels (Figure S15). However, data also revealed that not all cells responded with TG accumulation, suggesting a heterogeneous phenotype even when activated with the strong stimulant LPS. We confirmed the heterogeneous nature of LD formation with fluorescence microscopy as shown in Figure S23. Lipid droplets were found to be heterogeneously distributed in the cell culture, consistent with our AP-SMALDI MSI findings. Our results are also in line with reports for microglia in human and mouse brain that exhibited highly heterogeneous LD distributions with more cells containing increased LD levels upon LPS stimulation.^44,20^ Unlike in previous studies, our untargeted methodology identified heterogeneous lipid distributions between individual cells, most likely associated with LDs. In addition, AP-SMALDI MSI provided molecular information about the TG composition, a feature that cannot be achieved with fluorescence microscopy.

### Fatty acid composition of PI lipids differentiates MGL phenotypes

Other signals besides TGs were found to be heterogeneously distributed between cells. In negative-ion mode several phosphatidylinositols (PI) showed locally varying intensities. A reproducible pattern of locally different degrees of fatty acid (FA) saturation was found for PI 38:X species when measured with AP-SMALDI MSI (Figure 5 and Figure S8). While PI 38:4 (*m*/*z* 885.5498) was detected in all cells (Figure 5c, f), PI 38:3 (*m*/*z* 887.5655), and PI 38:5 (*m*/*z* 883.5342) were found enriched in different subpopulations (Figure 5b, e and S24). This heterogeneity might be linked to differential desaturase enzyme activity, which is known to occur for different microglia phenotypes.^45–47^ Interestingly, this variance was found in both, naïve and LPS stimulated cells (Figure S8) and can be used to classify cells based on hierarchical clustering of mass spectrometric data (Figure S25). This heterogeneity was found to be exclusive for PI 38:X species in our measurements. Other phospholipid classes such as PC 38:X or PE 38:X were found to be evenly distributed across all cells, making a heterogeneity that is caused by selective FA production or FA uptake unlikely. However, not all phospholipid classes are synthesized from FAs via the same pathways.^48^ While in mammalian cells PC and PE are synthesized via the Kennedy pathway,^49^ PI is synthesized via a cytidinediphosphate diacylglycerol (CDP-DG) intermediate.^50^ A selectivity for the synthesis of PI 38:4 over other fatty acid derivates was described earlier^51^ and is supported by the homogeneous expression in all cells in our experiments. Kim et al. have recently demonstrated that considerably FA remodeling of PIs compared to PA precursors takes place.^52^ This remodeling is most likely associated with lysophosphatidylinositol acetyltransferase (LPIAT) enzymes that could also be the cause for cell-specific PI expression.

**Figure 5:**
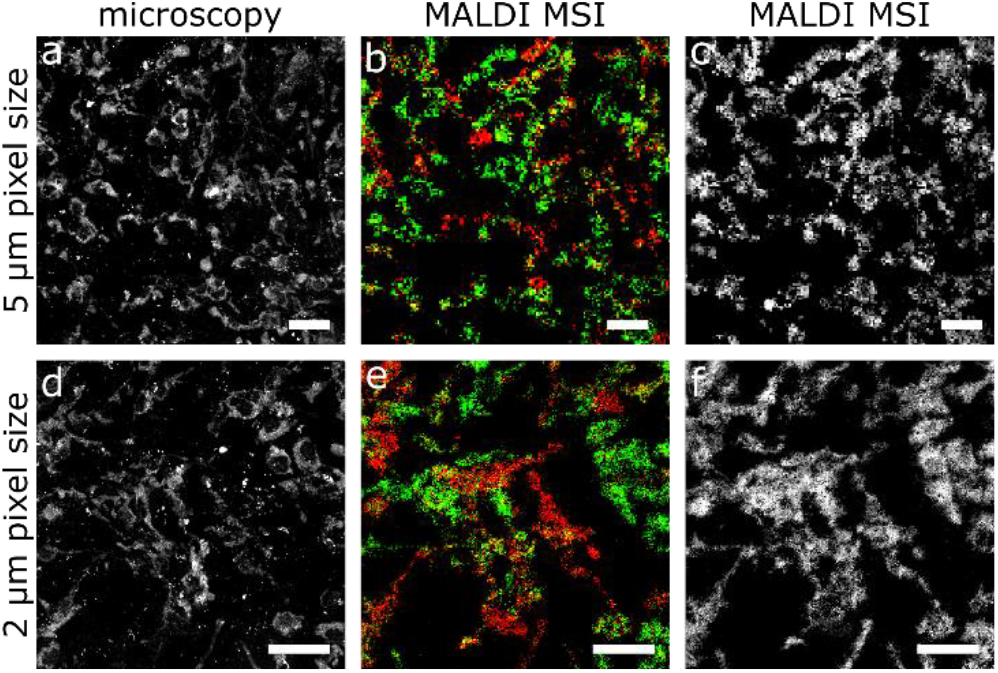
Degree of FA saturation differs between MGL cells. a,d: Microscopic image of the region investigated by AP-SMALDI MSI of naïve B8 MGL. b,c,d,e: AP-SMALDI MSI images acquired with 150×150 pixels at 5 μm pixel size (b,c) or 250×250 pixels at 2 μm pixel size (e,f) in negative-ion mode, showing in b,e the heterogeneity of PI 38:3 (m/z 887.5655, red) and PI 38:5 (m/z 883.5342, green) and in c,f the homogeneous distribution of PI 38:4 (m/z 885.5499, greyscale). Scale bars: 100 μm.

Also, PIs were recently found to be directly involved in cellular stress regulation.^53^ Combined with the heterogeneous stress-related lipid-droplet formation observed in our study, we hypothesize that cell-specific regulatory switches linked to LPIAT for PI metabolism exist, similar to recent reports for glycosphingolipids,^19^ leading to spontaneous formation of two distinct phenotypes. However, further research is needed to resolve the cause for the observation that MGL in culture exhibit heterogeneous PI distributions.

## Conclusion

We were able to visualize population-based and single-cell-based heterogeneities in fixed or snap-frozen human MGL, cultured on glass slides, using AP-SMALDI MSI with lateral resolutions of down to 1.5 μm. Multiple cell lines were cultured separately and developed unique lipidomic profiles despite their genetic similarity. We were able to differentiate multiple cell lines from each other in positive- and negative-ion mode, using high-mass-resolution data and statistical analysis on biological triplicates. Further, our method was able to identify and visualize inter- (e.g. LD) or intra-cellular (e.g. PI species) heterogeneity between individual cells of the same cell-culture, indicating the differentiation of MGL into different phenotypes, which is consistent with previous findings.^54,11–14^ On the one hand, we observed sub-cellular inflammation-induced heterogeneity expressed through lipid-droplet formation^44^. To the best of our knowledge, this is the first reported visualization of intact biomolecules within sub-cellular features, i.e. LDs, by MSI. LDs were localized at specific positions in some cells, whereas the rest of the cells remained unaffected, consistent with fluorescence microscopy and literature reports. This kind of heterogeneity was more pronounced in LPS-stimulated samples, due to LPS triggering the inflammatory response mechanism of microglia. On the other hand, single-cell heterogeneity affecting a complete cell body was found independent of cell line and stimulation in positive- and negative-ion mode, which was particularly prominent in negative-ion mode detecting PI lipid species. These findings together with recent reports of fibroblasts^19^ and multiple cell lines^55^ reveal that state-of-the-art MSI technologies allow to differentiate lipidomic phenotypes of single cells in culture and call for large scale studies to chart the heterogeneity of the cellular lipidome or even sub-cellular compositions within the context of results of other single-cell methods. This will facilitate further insights in biochemical switches of cell-specific lipid metabolism and plasticity.

## Acknowledgements

We gratefully acknowledge financial support by the German Science Foundation (DFG) under the grants Sp314/23-1 and INST 162/500-1 FUGG. M.A.M. thanks the Fonds der Chemischen Industrie for granting a Kekulé fellowship. S.H. thanks the Fonds der Chemischen Industrie for granting a Liebig fellowship, and financial support by the Deutsche Forschungsgemeinschaft (HE 8521/1-1) is gratefully acknowledged. S.H. acknowledges the support by the “Ministerium für Kultur und Wissenschaft des Landes Nordrhein-Westfalen” and the German Ministry of Research and Education (BMBF) and is grateful for financial support by the Justus Liebig University via the JLU price 2022.

## Author contributions

Original manuscript draft and figures: M.A.M.; manuscript review: M.A.M., S.H., M.W., B.S., N.Z.; cell growth: M.W.; sample preparation: M.A.M., N.Z.; AP-SMALDI MSI instrumentation and experiments: M.A.M., S.H., N.Z.; light microscopy: M.A.M., S.H.; fluorescence microscopy: M.W.; flow cytometry: M.W.; statistical evaluation: M.A.M.; software code: M.A.M.; review of data and planning of experiments: M.A.M., S.H., M.W.; concept and design of project: S.H. and M.W.; supervision: S.H. and B.S..

## Notes

### Competing Interest Statement

M.A.M. is employee and B.S. is consultant of TransMIT GmbH.

## References

(1) Lawson, L. J.; Perry, V. H.; Dri, P.; Gordon, S. Neuroscience 1990, DOI: 10.1016/0306-4522(90)90229-W.

(2) Lenz, K. M.; McCarthy, M. M. The Neuroscientist: a review journal bringing neurobiology, neurology and psychiatry 2015, DOI: 10.1177/1073858414536468.

(3) Butovsky, O.; Weiner, H. L. Nature reviews. Neuroscience 2018, DOI: 10.1038/s41583-018-0057-5.

(4) Li, Q.; Barres, B. A. Nature reviews. Immunology 2018, DOI: 10.1038/nri.2017.125.

(5) Salter, M. W.; Beggs, S. Cell 2014, DOI: 10.1016/j.cell.2014.06.008.

(6) Block, M. L.; Hong, J.-S. Progress in neurobiology 2005, DOI: 10.1016/j.pneurobio.2005.06.004.

(7) Cartier, N.; Lewis, C.-A.; Zhang, R.; Rossi, F. M. V. Acta Neuropathol 2014, DOI: 10.1007/s00401-014-1330-y.

(8) Colonna, M.; Butovsky, O. Annual review of immunology 2017, DOI: 10.1146/annurev-immunol-051116-052358.

(9) Efthymiou, A. G.; Goate, A. M. Mol Neurodegeneration 2017, DOI: 10.1186/s13024-017-0184-x.

(10) Loving, B. A.; Bruce, K. D. Frontiers in physiology 2020, DOI: 10.3389/fphys.2020.00393.

(11) Chhor, V.; Le Charpentier, T.; Lebon, S.; Oré, M.-V.; Celador, I. L.; Josserand, J.; Degos, V.; Jacotot, E.; Hagberg, H.; Sävman, K.; Mallard, C.; Gressens, P.; Fleiss, B. Brain, behavior, and immunity 2013, DOI: 10.1016/j.bbi.2013.02.005.

(12) Lauro, C.; Limatola, C. Front. Immunol. 2020, DOI: 10.3389/fimmu.2020.00493.

(13) Masuda, T.; Sankowski, R.; Staszewski, O.; Prinz, M. Cell reports 2020, DOI: 10.1016/j.celrep.2020.01.010.

(14) Orihuela, R.; McPherson, C. A.; Harry, G. J. British journal of pharmacology 2016, DOI: 10.1111/bph.13139.

(15) Li, Z.; Cheng, S.; Lin, Q.; Cao, W.; Yang, J.; Zhang, M.; Shen, A.; Zhang, W.; Xia, Y.; Ma, X.; Ouyang, Z. Nat Commun 2021, DOI: 10.1038/s41467-021-23161-5.

(16) Smith, A. M.; Davey, K.; Tsartsalis, S.; Khozoie, C.; Fancy, N.; Tang, S. S.; Liaptsi, E.; Weinert, M.; McGarry, A.; Muirhead, R. C. J.; Gentleman, S.; Owen, D. R.; Matthews, P. M. Acta Neuropathol 2022, DOI: 10.1007/s00401-021-02372-6.

(17) Hannun, Y. A.; Obeid, L. M. Nature reviews. Molecular cell biology 2008, DOI: 10.1038/nrm2329.

(18) Hu, T.; Zhang, J.-L. Journal of separation science 2018, DOI: 10.1002/jssc.201700709.

(19) Capolupo, L.; Khven, I.; Lederer, A. R.; Mazzeo, L.; Glousker, G.; Ho, S.; Russo, F.; Montoya, J. P.; Bhandari, D. R.; Bowman, A. P.; Ellis, S. R.; Guiet, R.; Burri, O.; Detzner, J.; Muthing, J.; Homicsko, K.; Kuonen, F.; Gilliet, M.; Spengler, B.; Heeren, R. M. A.; Dotto, G. P.; La Manno, G.; D’Angelo, G. Science (New York, N.Y.) 2022, DOI: 10.1126/science.abh1623.

(20) Marschallinger, J.; Iram, T.; Zardeneta, M.; Lee, S. E.; Lehallier, B.; Haney, M. S.; Pluvinage, J. V.; Mathur, V.; Hahn, O.; Morgens, D. W.; Kim, J.; Tevini, J.; Felder, T. K.; Wolinski, H.; Bertozzi, C. R.; Bassik, M. C.; Aigner, L.; Wyss-Coray, T. Nat Neurosci 2020, DOI: 10.1038/s41593-019-0566-1.

(21) Riera-Borrull, M.; Cuevas, V. D.; Alonso, B.; Vega, M. A.; Joven, J.; Izquierdo, E.; Corbí, Á. L. Journal of immunology (Baltimore, Md.: 1950) 2017, DOI: 10.4049/jimmunol.1700845.

(22) Bandu, R.; Mok, H. J.; Kim, K. P. Mass spectrometry reviews 2018, DOI: 10.1002/mas.21510.

(23) Tian, H.; Sparvero, L. J.; Anthonymuthu, T. S.; Sun, W.-Y.; Amoscato, A. A.; He, R.-R.; Bayir, H.; Kagan, V. E.; Winograd, N. Analytical chemistry 2021, DOI: 10.1021/acs.analchem.0c05311.

(24) Blank, M.; Enzlein, T.; Hopf, C. Sci Rep 2022, DOI: 10.1038/s41598-022-06894-1.

(25) Chausse, B.; Kakimoto, P. A.; Caldeira-da-Silva, C. C.; Chaves-Filho, A. B.; Yoshinaga, M. Y.; da Silva, R. P.; Miyamoto, S.; Kowaltowski, A. J. Bioscience reports 2019, DOI: 10.1042/BSR20190072.

(26) Römpp, A.; Spengler, B. Histochem Cell Biol 2013, DOI: 10.1007/s00418-013-1097-6.

(27) Spengler, B. Analytical chemistry 2015, DOI: 10.1021/ac504543v.

(28) Haenseler, W.; Sansom, S. N.; Buchrieser, J.; Newey, S. E.; Moore, C. S.; Nicholls, F. J.; Chintawar, S.; Schnell, C.; Antel, J. P.; Allen, N. D.; Cader, M. Z.; Wade-Martins, R.; James, W. S.; Cowley, S. A. Stem cell reports 2017, DOI: 10.1016/j.stemcr.2017.05.017.

(29) Vaughan-Jackson, A.; Stodolak, S.; Ebrahimi, K. H.; Browne, C.; Reardon, P. K.; Pires, E.; Gilbert-Jaramillo, J.; Cowley, S. A.; James, W. S. Stem cell reports 2021, DOI: 10.1016/j.stemcr.2021.05.018.

(30) Lund, S.; Christensen, K. V.; Hedtjärn, M.; Mortensen, A. L.; Hagberg, H.; Falsig, J.; Hasseldam, H.; Schrattenholz, A.; Pörzgen, P.; Leist, M. Journal of neuroimmunology 2006, DOI: 10.1016/j.jneuroim.2006.07.007.

(31) Paschke, C.; Leisner, A.; Hester, A.; Maass, K.; Guenther, S.; Bouschen, W.; Spengler, B. Journal of the American Society for Mass Spectrometry 2013, DOI: 10.1007/s13361-013-0667-0.

(32) Müller, M. A.; Kompauer, M.; Strupat, K.; Heiles, S.; Spengler, B. Journal of the American Society for Mass Spectrometry 2021, DOI: 10.1021/jasms.0c00368.

(33) Tyanova, S.; Temu, T.; Sinitcyn, P.; Carlson, A.; Hein, M. Y.; Geiger, T.; Mann, M.; Cox, J. Nat Methods 2016, DOI: 10.1038/nmeth.3901.

(34) Bechtold, B. Violin Plots for Matlab, https://github.com/bastibe/Violinplot-Matlb.

(35) Palmer, A.; Phapale, P.; Chernyavsky, I.; Lavigne, R.; Fay, D.; Tarasov, A.; Kovalev, V.; Fuchser, J.; Nikolenko, S.; Pineau, C.; Becker, M.; Alexandrov, T. Nature Methods 2017, DOI: 10.1038/nmeth.4072.

(36) Sud, M.; Fahy, E.; Cotter, D.; Brown, A.; Dennis, E. A.; Glass, C. K.; Merrill, A. H.; Murphy, R. C.; Raetz, C. R. H.; Russell, D. W.; Subramaniam, S. Nucleic acids research 2007, DOI: 10.1093/nar/gkl838.

(37) Aimo, L.; Liechti, R.; Hyka-Nouspikel, N.; Niknejad, A.; Gleizes, A.; Götz, L.; Kuznetsov, D.; David, F. P. A.; van der Goot, F. Gisou; Riezman, H.; Bougueleret, L.; Xenarios, I.; Bridge, A. Bioinformatics 2015, DOI: 10.1093/bioinformatics/btv285.

(38) Fitzner, D.; Bader, J. M.; Penkert, H.; Bergner, C. G.; Su, M.; Weil, M.-T.; Surma, M. A.; Mann, M.; Klose, C.; Simons, M. Cell reports 2020, DOI: 10.1016/j.celrep.2020.108132.

(39) Feingold, K. R.; Shigenaga, J. K.; Kazemi, M. R.; McDonald, C. M.; Patzek, S. M.; Cross, A. S.; Moser, A.; Grunfeld, C. Journal of leukocyte biology 2012, DOI: 10.1189/jlb.1111537.

(40) Funk, J. L.; Feingold, K. R.; Moser, A. H.; Grunfeld, C. Atherosclerosis 1993, DOI: 10.1016/0021-9150(93)90224-I.

(41) Hamsanathan, S.; Gurkar, A. U. Frontiers in physiology 2022, DOI: 10.3389/fphys.2022.796850.

(42) Olzmann, J. A.; Carvalho, P. Nat Rev Mol Cell Biol 2019, DOI: 10.1038/s41580-018-0085-z.

(43) Churchward, M. A.; Tchir, D. R.; Todd, K. G. Mol Neurobiol 2018, DOI: 10.1007/s12035-017-0422-9.

(44) Khatchadourian, A.; Bourque, S. D.; Richard, V. R.; Titorenko, V. I.; Maysinger, D. Biochimica et biophysica acta 2012, DOI: 10.1016/j.bbalip.2012.01.007.

(45) Bogie, J. F. J.; Grajchen, E.; Wouters, E.; Corrales, A. G.; Dierckx, T.; Vanherle, S.; Mailleux, J.; Gervois, P.; Wolfs, E.; Dehairs, J.; van Broeckhoven, J.; Bowman, A. P.; Lambrichts, I.; Gustafsson, J.-Å.; Remaley, A. T.; Mulder, M.; Swinnen, J. V.; Haidar, M.; Ellis, S. R.; Ntambi, J. M.; Zelcer, N.; Hendriks, J. J. A. The Journal of experimental medicine 2020, DOI: 10.1084/jem.20191660.

(46) Garcia Corrales, A. V.; Haidar, M.; Bogie, J. F. J.; Hendriks, J. J. A. International journal of molecular sciences 2021, DOI: 10.3390/ijms22158159.

(47) Guillou, H.; Zadravec, D.; Martin, P. G. P.; Jacobsson, A. Progress in lipid research 2010, DOI: 10.1016/j.plipres.2009.12.002.

(48) Harayama, T.; Shimizu, T. Journal of Lipid Research 2020, DOI: 10.1194/jlr.R120000800.

(49) Kennedy, E. P.; Weiss, S. B. Journal of Biological Chemistry 1956, DOI: 10.1016/S0021-9258(19)50785-2.

(50) Blunsom, N. J.; Cockcroft, S. Front. Cell Dev. Biol. 2020, DOI: 10.3389/fcell.2020.00063.

(51) Barneda, D.; Janardan, V.; Niewczas, I.; Collins, D. M.; Cosulich, S.; Clark, J.; Stephens, L. R.; Hawkins, P. T. The EMBO journal 2022, DOI: 10.15252/embj.2021110038.

(52) Kim, Y. J.; Sengupta, N.; Sohn, M.; Mandal, A.; Pemberton, J. G.; Choi, U.; Balla, T. EMBO reports 2022, DOI: 10.15252/embr.202154532.

(53) Thürmer, M.; Gollowitzer, A.; Pein, H.; Neukirch, K.; Gelmez, E.; Waltl, L.; Wielsch, N.; Winkler, R.; Löser, K.; Grander, J.; Hotze, M.; Harder, S.; Döding, A.; Meßner, M.; Troisi, F.; Ardelt, M.; Schlüter, H.; Pachmayr, J.; Gutiérrez-Gutiérrez, Ó.; Rudolph, K. L.; Thedieck, K.; Schulze-Späte, U.; González-Estévez, C.; Kosan, C.; Svatoš, A.; Kwiatkowski, M.; Koeberle, A. Nat Commun 2022, DOI: 10.1038/s41467-022-30374-9.

(54) Boche, D.; Perry, V. H.; Nicoll, J. A. R. Neuropathology and Applied Neurobiology 2013, DOI: 10.1111/nan.12011.

(55) Bien, T.; Koerfer, K.; Schwenzfeier, J.; Dreisewerd, K.; Soltwisch, J. Proceedings of the National Academy of Sciences of the United States of America 2022, DOI: 10.1073/pnas.2114365119.

