## Supporting Information for "Lipid signatures and inter-cellular heterogeneity of naïve and lipopolysaccharide-stimulated human microglia-like cells"

*Address correspondence to:
Sven Heiles
Leibniz-Institut für Analytische Wissenschaften - ISAS - e.V.
Otto-Hahn-Straße 6b
44227 Dortmund, Germany


### Table of content


| **Methods** | | **page** |
|---|---|---|
| | Cell pretreatment for AP-SMALDI MSI | 4 |
| | Matrix optimization and application protocols | 4 |
| **Figures** | | |
| S1 | Microscopic image of the optimized 9-AA matrix surface | 5 |
| S2 | Ablation spot sizes for 100 induvial spots on red dye | 5 |
| S3 | Ablation spot sizes on DHB matrix for different laser energy settings | 6 |
| S4 | Sample preparation workflow | 6 |
| S5 | Microscopic images of microglia cells before and after washing | 7 |
| S6 | Washout effect in negative-ion mode | 8 |
| S7 | MALDI MSI showcasing triglyceride distribution in flash-frozen microglia at different pixel sizes | 8 |
| S8 | MALDI MSI showcasing PI 38:X heterogeneity in naïve and LPS-stimulated microglia | 9 |
| S9 | MALDI MSI showcasing phospholipid heterogeneity | 9 |
| S10 | Mass spectra at different pixel sizes | 10 |
| S11 | Intensity contrast on microglia projections at 1.5 µm pixel size | 11 |
| S12 | Pixel coverage at different pixel sizes | 12 |
| S13 | Population-based heterogeneity between naïve and LPS-stimulated microglia | 13 |
| S14 | PCA-plots for all statistical comparisons | 14 |
| S15 | Violin plot of triglyceride intensity comparing naïve and LPS stimulated microglia for different cell lines | 14 |
| S16 | List of detected triglyceride species in different cell lines | 15 |
| S17 | Number of signals annotated as triglycerides in Metaspace | 16 |
| S18 | iPSC-derived microglia precursor identity confirmation based on the expression of myeloid marker proteins | 16 |
| S19 | Stitched MALDI MSI results for triglyceride heterogeneity at 5 µm pixel size with phospholipid background for cell morphology | 17 |
| S20 | Stitched MALDI MSI results for triglyceride heterogeneity at 5 µm pixel size | 17 |
| S21 | Statistical classification of lipid droplets for fixed microglia using t-SNE allowing for 3 clusters | 18 |
| S22 | Statistical classification of lipid droplets for flash-frozen microglia using t-SNE | 19 |
| S23 | Fluorescence microscopy of microglia cells using BODIPY and DAPI | 20 |
| S24 | Single-pixel mass spectra in negative-ion mode | 20 |
| S25 | Statistical classification of phenotypes for flash-frozen microglia using PCA allowing for identification of 3 clusters by a k-means algorithm | 21 |
| **Tables** | | |
| S1 | List of annotations from statistical analysis between naïve OX1, D9, C12 and E2 cells for positive-ion mode | 22 |
| S2 | List of annotations from statistical analysis between LPS-stimulated OX1, D9, C12 and E2 cells for positive-ion mode | 23 |
| S3 | List of annotations from statistical analysis between naïve and LPS-stimulated OX1 cells for positive-ion mode | 34 |

| | | |
|---|---|---|
| S4 | List of annotations from statistical analysis between naïve and LPS-stimulated D9 cells for positive-ion mode | 35 |
| S5 | List of annotations from statistical analysis between naïve and LPS-stimulated C12 cells for positive-ion mode | 38 |
| S6 | List of annotations from statistical analysis between naïve and LPS-stimulated E2 cells for positive-ion mode | 42 |
| S7 | List of annotations from statistical analysis between naïve OX1, B8, C12 and E2 cells for negative-ion mode | 45 |
| S8 | List of annotations from statistical analysis between LPS-stimulated OX1, B8, C12 and E2 cells for negative-ion mode | 55 |
| S9 | List of annotations from statistical analysis between naïve and LPS-stimulated OX1 cells for negative-ion mode | 56 |
| S10 | List of annotations from statistical analysis between naïve and LPS-stimulated B8 cells for negative-ion mode | 58 |
| S11 | List of annotations from statistical analysis between naïve and LPS-stimulated C12 cells for negative-ion mode | 62 |
| S12 | List of annotations from statistical analysis between naïve and LPS-stimulated E2 cells for negative-ion mode | 63 |

**Information**

| | | |
|---|---|---|
| | References | 64 |

#### Cell pretreatment for AP-SMALDI MSI

Microscopic images of the samples upon warming to room temperature and drying revealed an irregular coverage with salt residues potentially affecting lipid ion yields in MALDI MSI experiments (Figure S1). For example, lipid annotations of unfixed LPS-stimulated microglia cells from the OX1 cell line using Metaspace[1] (false discovery rate 10%) based on the SwissLipids[2] database reveal a high number of sodium adducts (116) being detected, whereas protonated signals (33) were suppressed. An additional washing step with 250 µL of deionized water flowing over individual cell-compartments with the sample tilted to facilitate rinsing, followed by drying at room temperature in an evacuated desiccator (50 mbar) for 10 minutes removed salt residues efficiently. Washed samples showed a more even ratio of protonated (156) and sodiated (78) adducts under otherwise same conditions. Sample morphology was unaffected by the washing process (Figure S2). However, cell morphology was preserved best for cells with PFA fixation.

#### Matrix optimization and application protocols

The choice of the matrix substance and its preparation protocol play a major role for ionization in MALDI MSI. In positive-ion mode, the above-described sample pretreatment for flash-frozen cells led to 150 annotations with Metaspace using 2,5-dihydroxybenzoic acid (DHB), an approximately 3-fold increase compared to annotations using the α-Cyano-4-hydroxycinnamic acid (CHCA) matrix (55). In negative-ion mode, matrix application of 9-AA was optimized by varying solvent composition, matrix concentration and application volume, since recently reported approaches[3,4] did result in intense washout effects. The spatial distribution of analytes was not preserved but dispersed all over the measurement area, resulting in a highly blurred representation of cellular features. (Figure S2). With the newly developed preparation method, the sample was covered homogeneously with small crystals (< 1 µm, Figure S22) and minimal thickness variations, which are favorable characteristics for high-resolution MALDI MSI.[5]

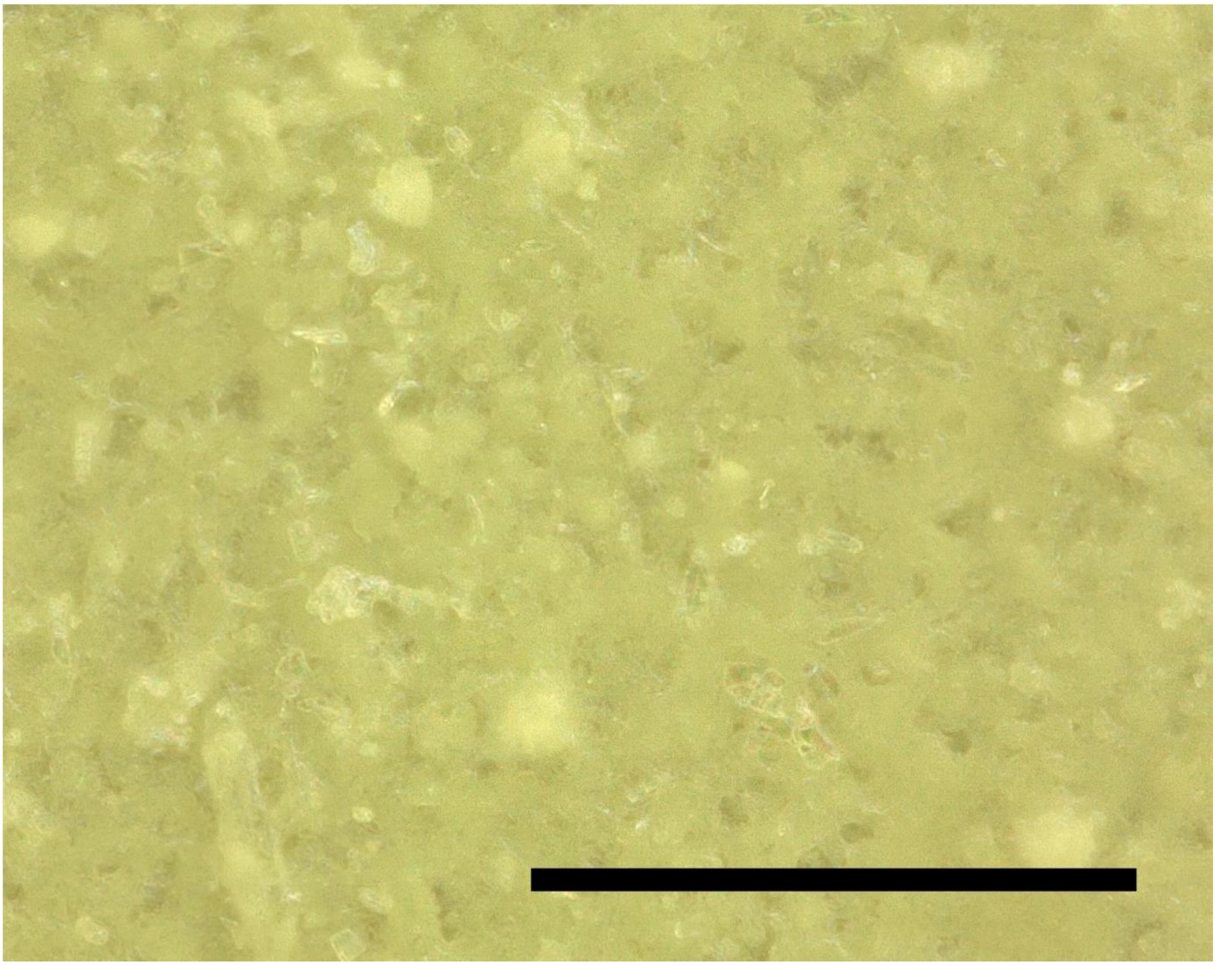

*Supporting Figure 1: Microscopic image of 9-AA matrix applied to microglia cells. Scale bar: 100 µm.*

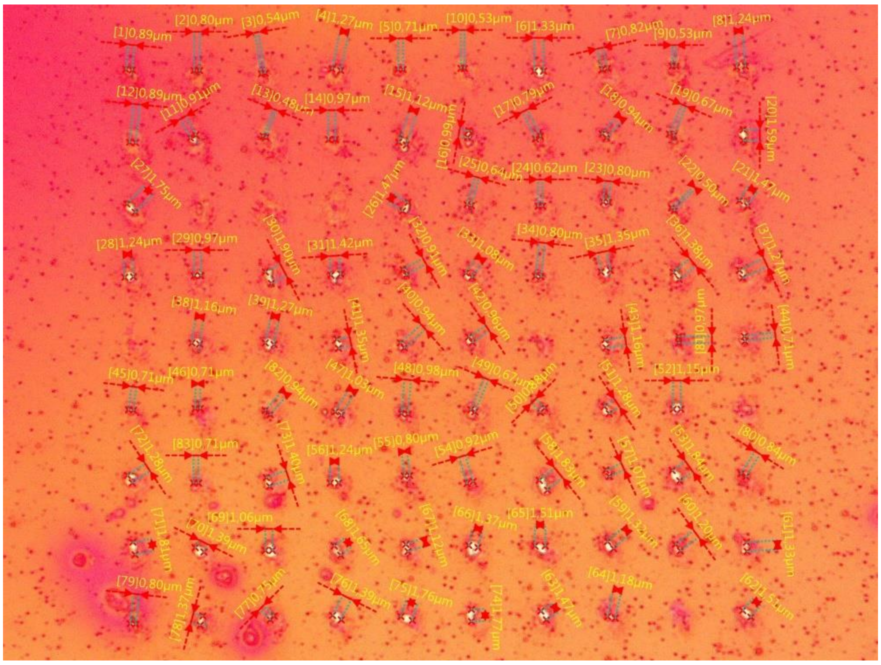

*Supporting Figure 2: Microscopic image of 100 laser ablation spots in a 10x10 grid acquired with a step size of 10 µm on red dye. Measurements of spot diameter were performed with a digital microscope and showed a mean spot diameter of 1.1 µm.*

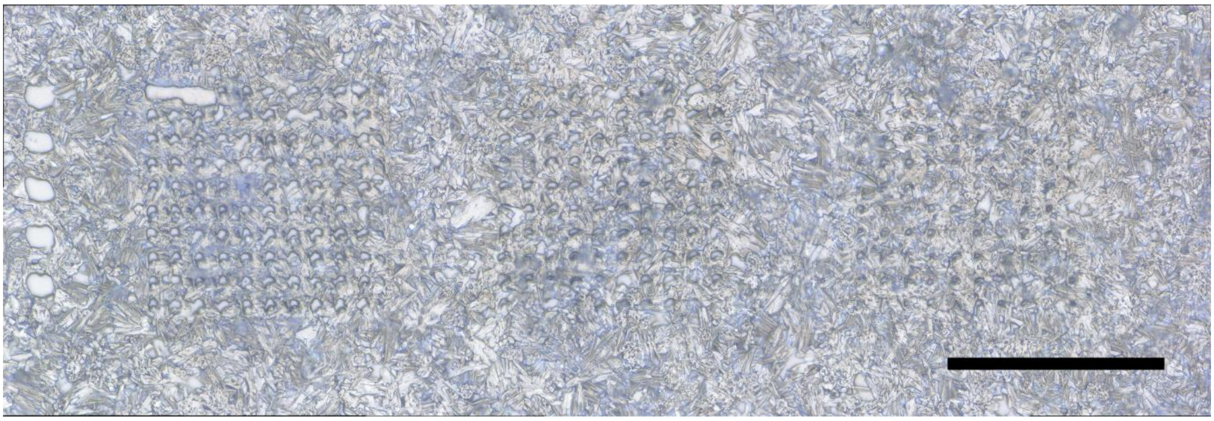

*Supporting Figure 3: Microscopic images of laser ablation spots on microglia cells covered with DHB matrix at different laser energy settings. Laser energy was decreased from left to right. Scale bar: 100 µm.*

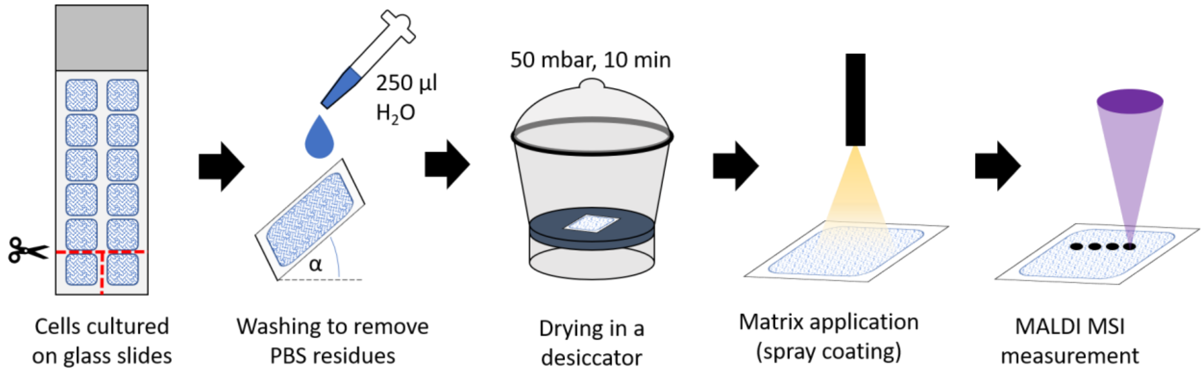


*Supporting Figure 4: Scheme of the sample preparation workflow of cells grown on glass slides for MALDI MSI at high lateral resolution.*

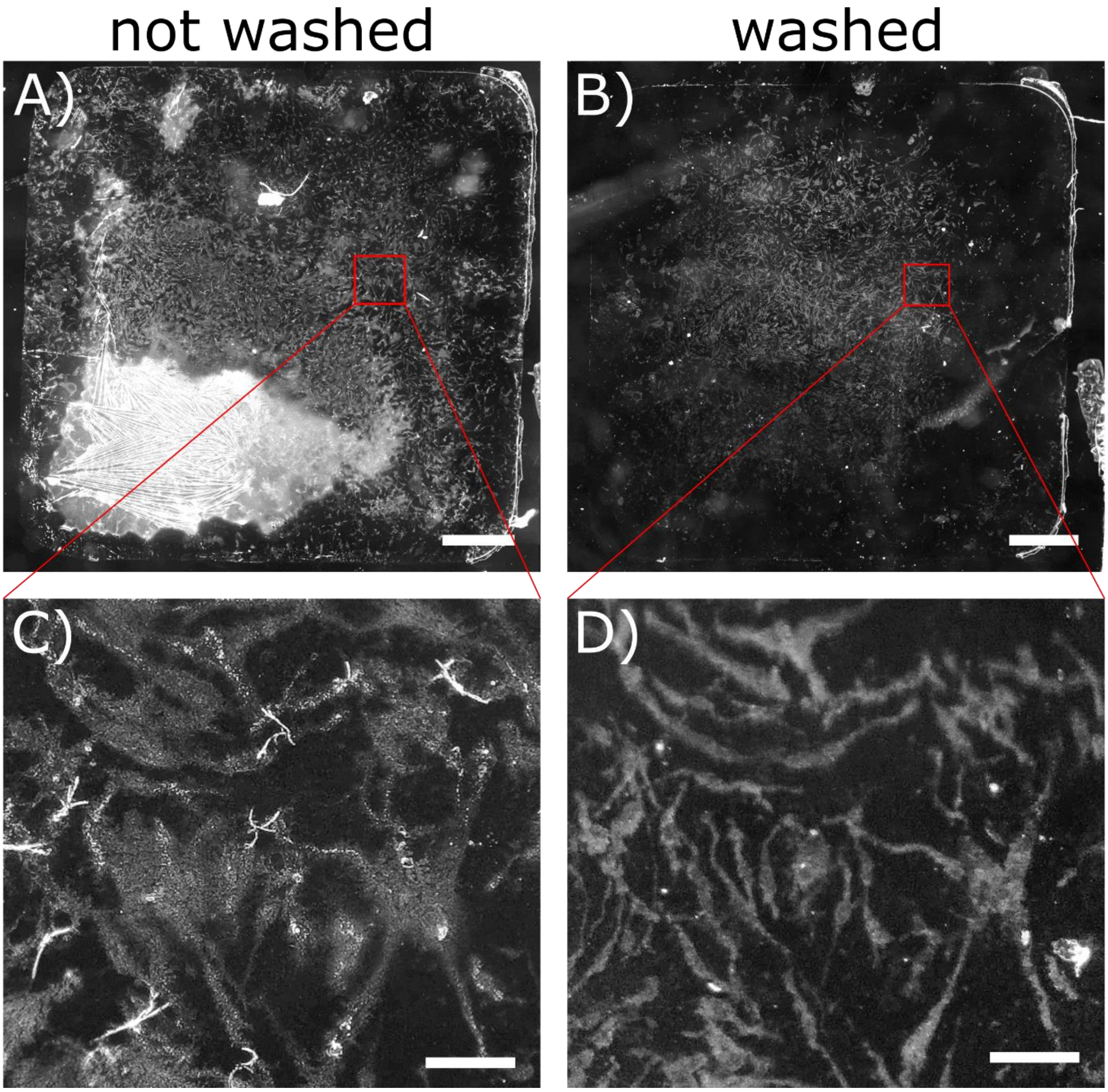


*Supporting Figure 5: Microscopic images of LPS-stimulated MGL from the OX1 cell line before (A, C) and after washing with deionized water (B,D). Zoom-in area indicated by red rectangles. Unwashed samples were covered with a salt layer of varying thickness originating from culturing in PBS. Sample morphology was preserved after washing (compare C and D). Scale bar: 1 mm (A, B) or 100 µm (C, D).*

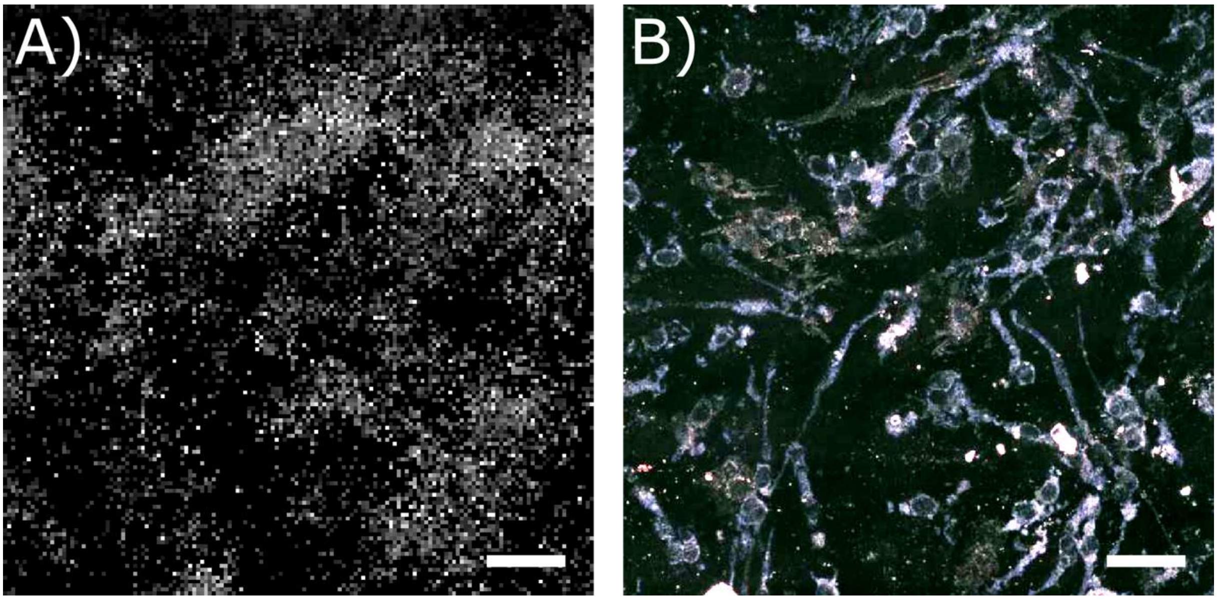


*Supporting Figure 6: Wash-out effect of analytes from naïve MGL from the B8 cell line with an inappropriate matrix preparation protocol in negative-ion mode. A) MALDI MSI image of [PE 36:2 – H]$^{-}$ (m/z 742.5392) with 150x150 pixels at 5 µm pixel size showing little to no morphological features. B) Corresponding microscopic image of the area investigated in A). Scale bars: 100 µm.*

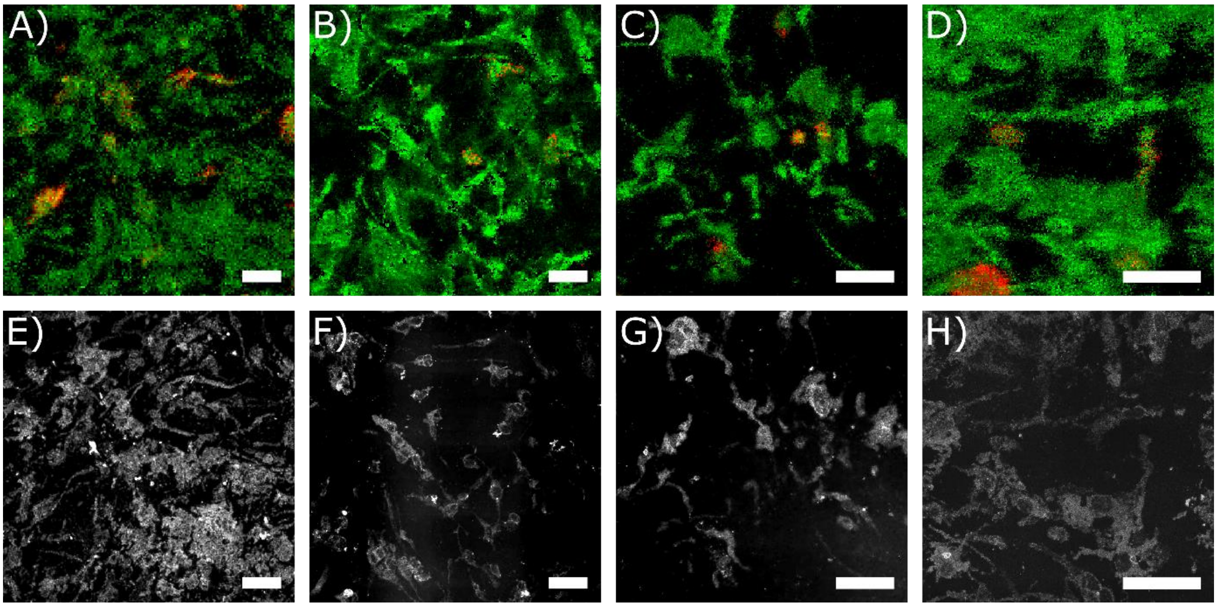


*Supporting Figure 7: MALDI MSI images of microglia cells visualizing the distribution of triglycerides represented by [TG 52:2 + Na]$^{+}$ (red, m/z 881.7569) compared to evenly distributed [PC 36:2 + Na]$^{+}$ (green, m/z 808.5827). A): LPS-stimulated D9 MGL cells, 150x150 pixels, 5 µm pixel size. B): LPS-stimulated E2 MGL cells, 250x250 pixels, 3 µm pixel size. C): Naïve D9 MGL cells, 250x250 pixels, 2 µm pixel size. D): LPS-stimulated D9 MGL cells, 250x250 pixels, 1.5 µm pixel size. E)-H): Corresponding microscopic images of the areas investigated. Scale bars: 100 µm.*

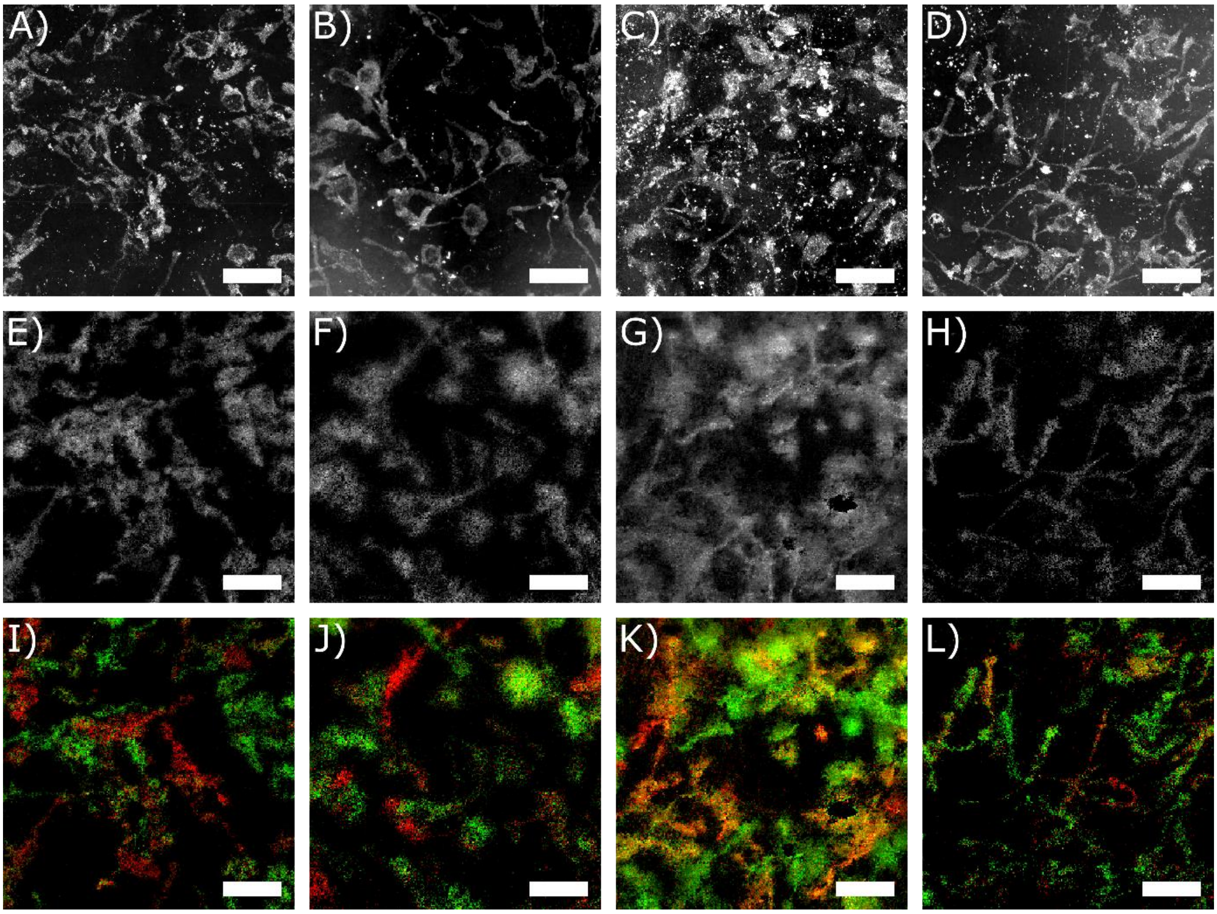


*Supporting Figure 8: Examples for cell-to-cell heterogeneity found in MALDI MSI experiments (250x250 pixels, 2 µm pixel size) using negative-ion mode for naïve (A, B, E, F, I, J) and LPS-stimulated (C, D, G, H, K, L) B8 MGL cells. A)-D): Microscopic images of the regions investigated. E)-H): MALDI MSI images of [PI 38:4 - H]$^-$ (m/z 885.5498) as an example for a homogeneously distributed lipid. I)-L): MALDI MSI images of [PI 38:3 - H]$^-$ (m/z 887.5655, red) and [PI 38:5 - H]$^-$ (m/z 883.5342, green) showing a heterogeneous distribution of these lipids. Scale bars: 100 µm.*

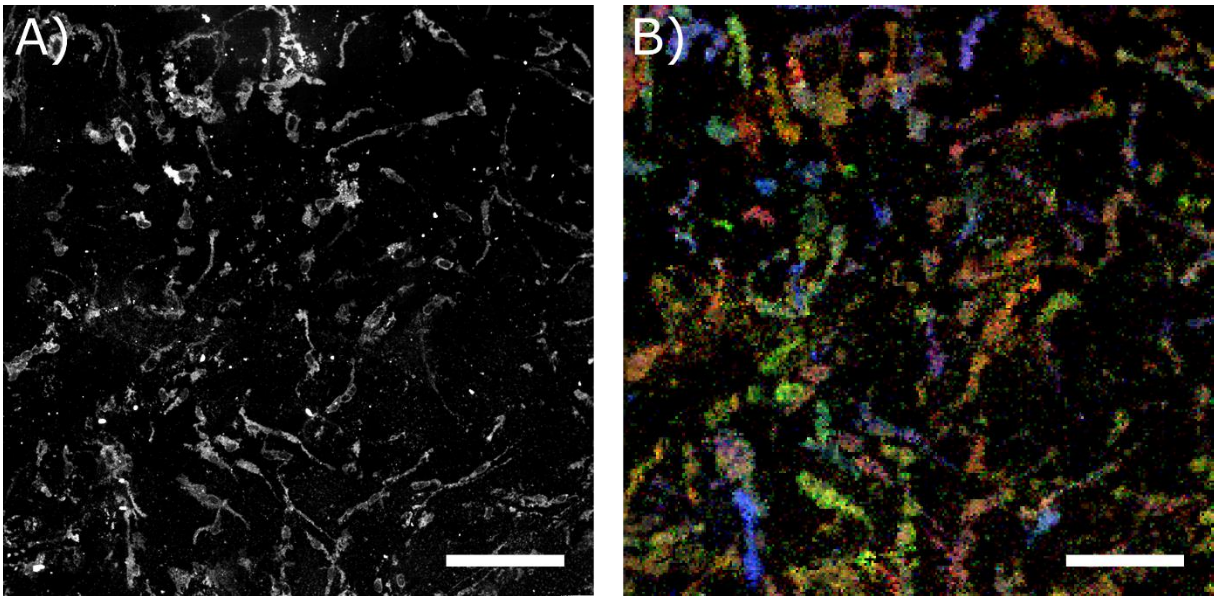


*Supporting Figure 9: Example for cell-to-cell heterogeneity found in MALDI MSI experiments (250x250 pixels, 5 µm) using negative ion mode for LPS-stimulated B8 MGL cells. A): Microscopic image of the region investigated. B) MALDI MSI image of [PI 38:4 - H]$^-$ (m/z 885.5501, red), [PI 38:3 - H]$^-$ (m/z 887.5653, green) and [PI 36:1 - H]$^-$ (m/z 863.5657, blue). Scale bars: 250 µm.*

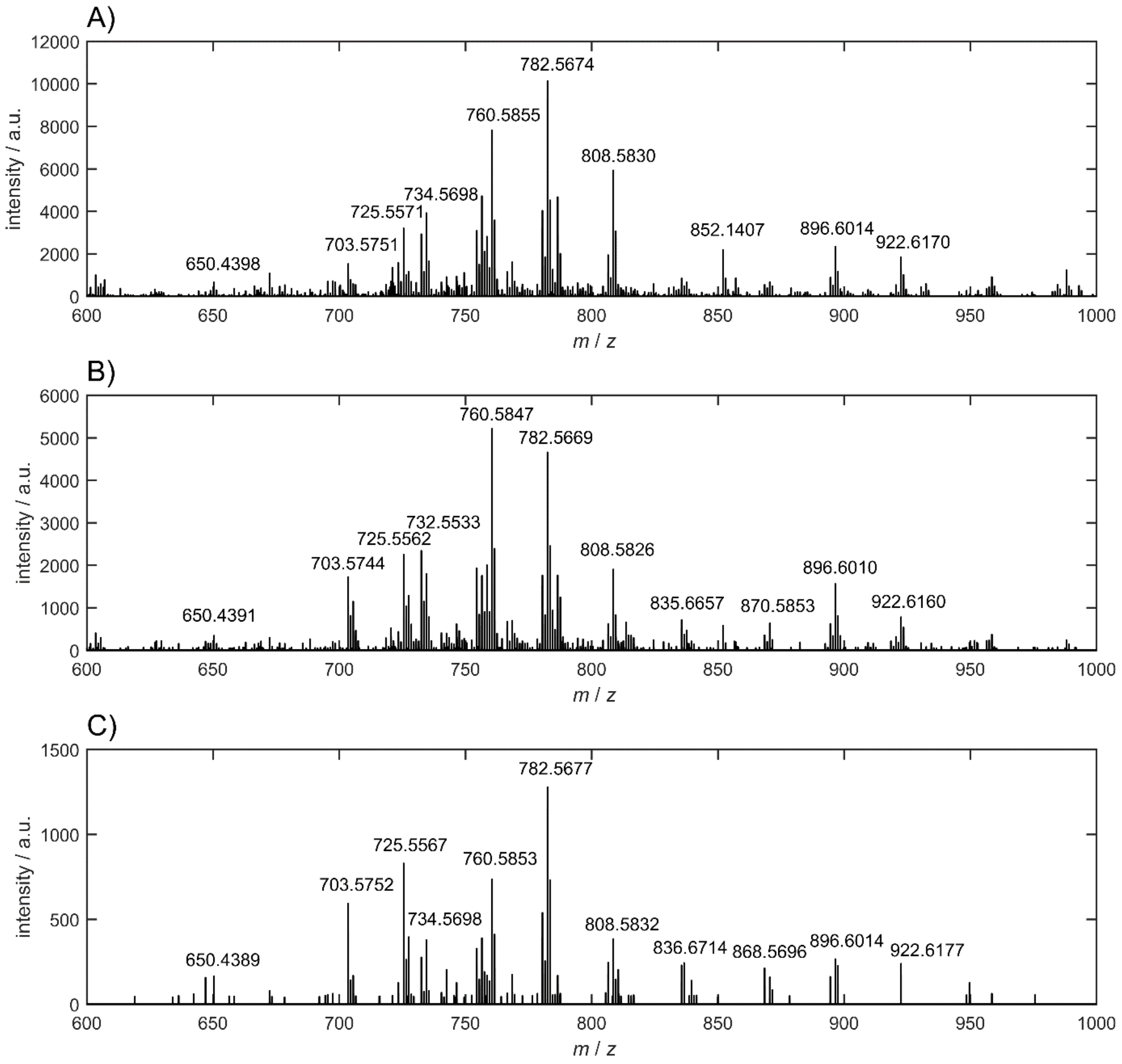


*Supporting Figure 10: Exemplary single-pixel mass spectra from microglia in positive-ion mode at different lateral resolution settings. A) 5 µm pixel size (LPS-stimulated D9 MGL cells), B) 3 µm pixel size (naïve D9 MGL cells), C) 1.5 µm pixel size (naïve E2 MGL cells).*

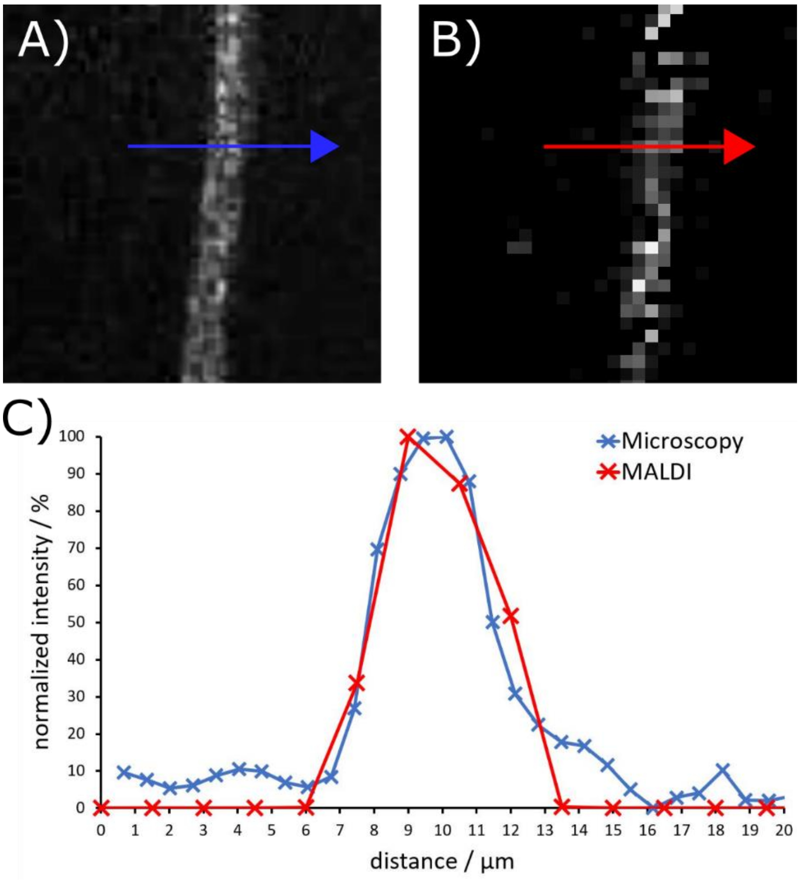


*Supporting Figure 11: Intensity contrast microglia projections from naïve B8 MGL cells in microscopy and MALDI MSI on fixed samples. A) Section of a microscopic image of a microglia projection. B) Section of a MALDI MSI image at 1.5 µm pixel size. C) Intensity plot over a microglia projection as indicated by arrows in A) and B) extracted from a microscopy image (blue) and a MALDI MSI image at 1.5 µm pixel size (red).*

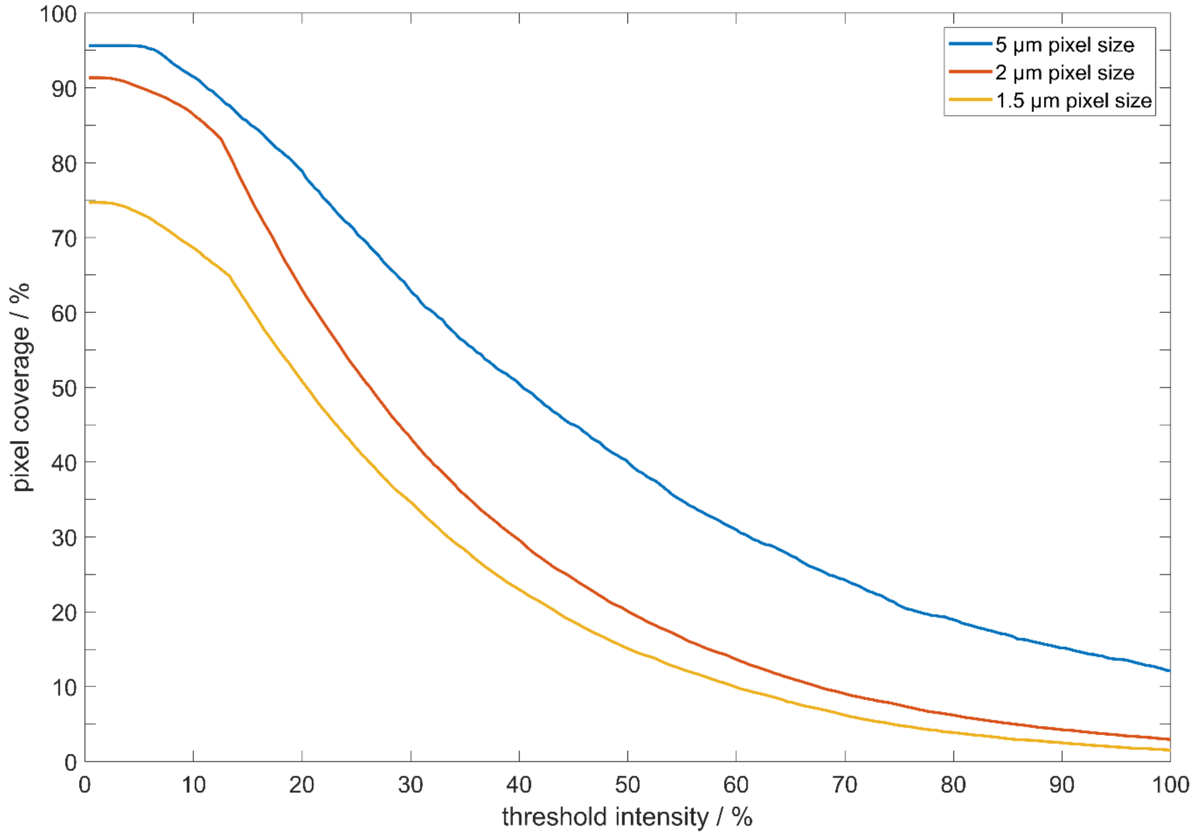


*Supporting Figure 12: Plot of percentage of pixels, in which the analyte ion signal (exemplary for PC 34:1) is above a given intensity threshold (relative to highest intensity pixel) at different lateral resolution settings. Pixel coverage is normalized to total number of on-cell pixels.*

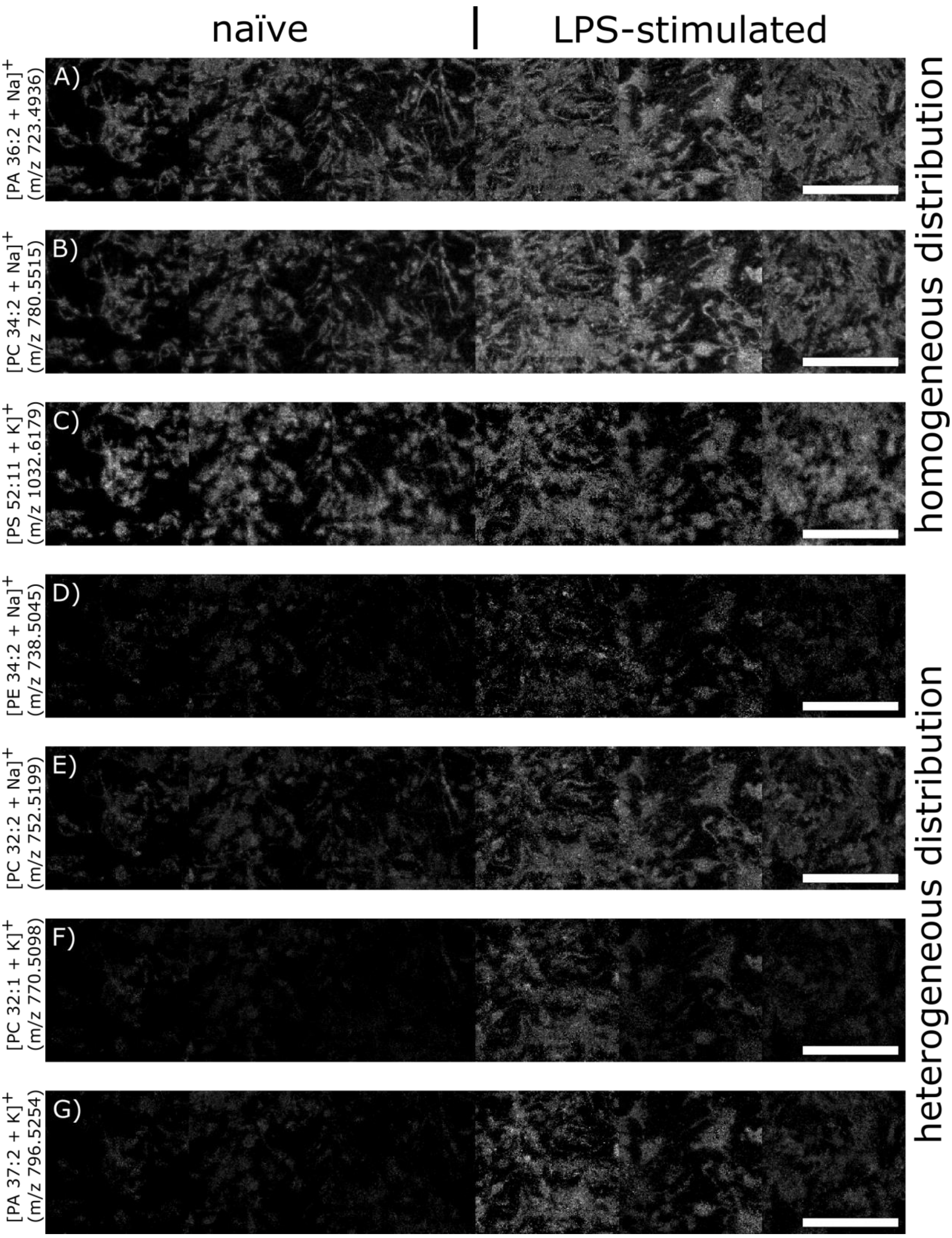


*Supporting Figure 13: Exemplary MALDI MSI images of phospholipid signals in the D9 cell line, distributed comparably (A-C) or dissimilarly (D-G) between naïve and LPS-stimulated cells. Each image consists of individual measurements of 3 naïve and 3 LPS-stimulated samples with 150x150 pixels at 5 µm pixel size, respectively. Scale bars: 500 µm.*

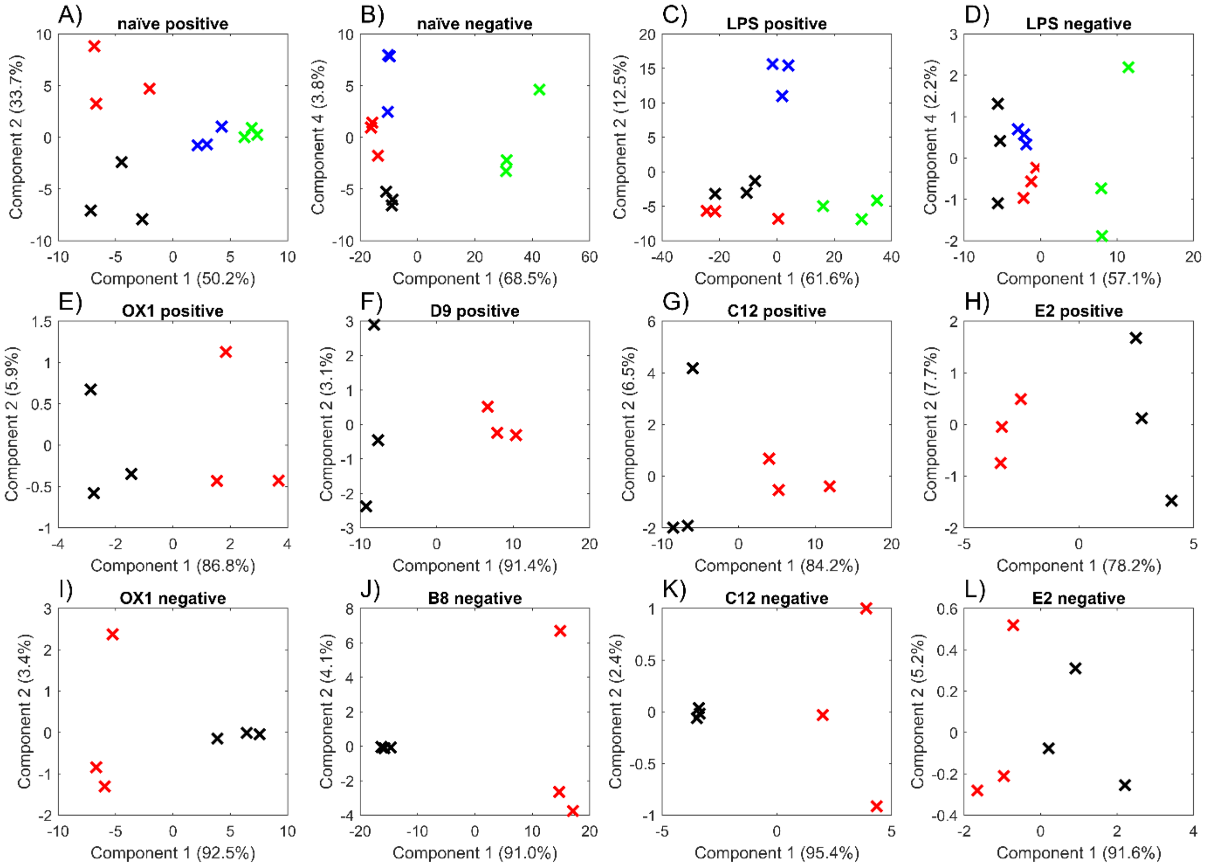


*Supporting Figure 14: PCA-plots obtained from statistical analysis of MALDI MSI results at 5 µm pixel size by ANOVA-based multiple-sample test. Three biological replicates of cell cultures grown in separate wells on the same glass slide, each containing a multitude of individual cells, were used for analysis and mass spectrometric signal intensities were averaged over 22,500 mass spectra. P-values were set to 0.01 (A, B, C, D, F, I, J, K) or 0.05 (E, G, H, L), respectively. Signals that passed the multiple-sample test can be found in Tables S1-S12. Color coding: A)-D): OX1 (red), D9 (green, positive-ion mode), B8 (green, negative-ion mode), C12 (blue), E2 (black). E)-L): naïve (red), LPS-stimulated (black).*

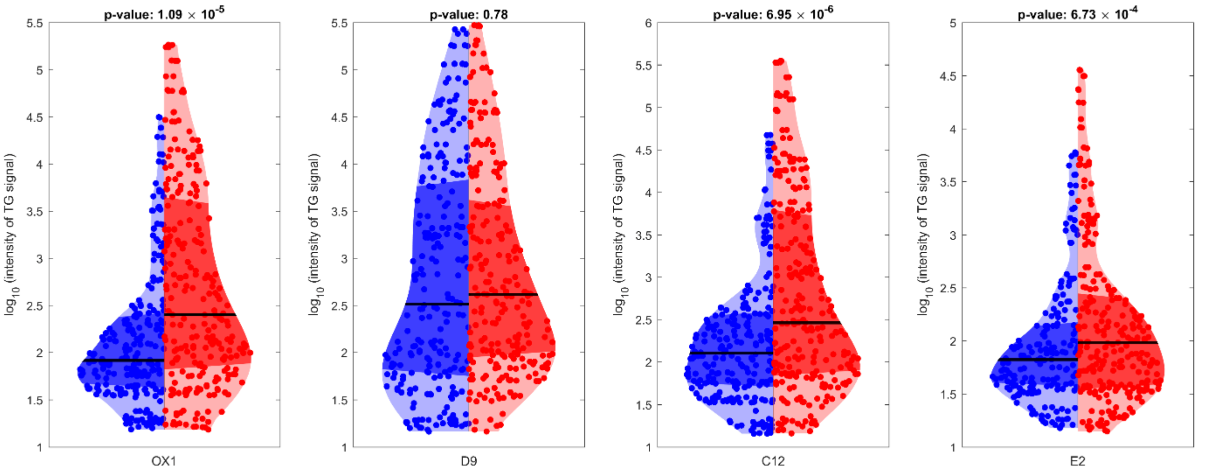


*Supporting Figure 15: Violin plots of TG-associated signal intensities from a 5 µm pixel size measurement in positive-ion mode of naïve (blue) and LPS-stimulated (red) MGL cells from the OX1, D9, C12 and E2 line, respectively. Each data point corresponds to a TG-associated mass spectrometric signal, averaged from 3 biological replicates. The black line indicates the median. p-Values calculated by double-sided t-test.*

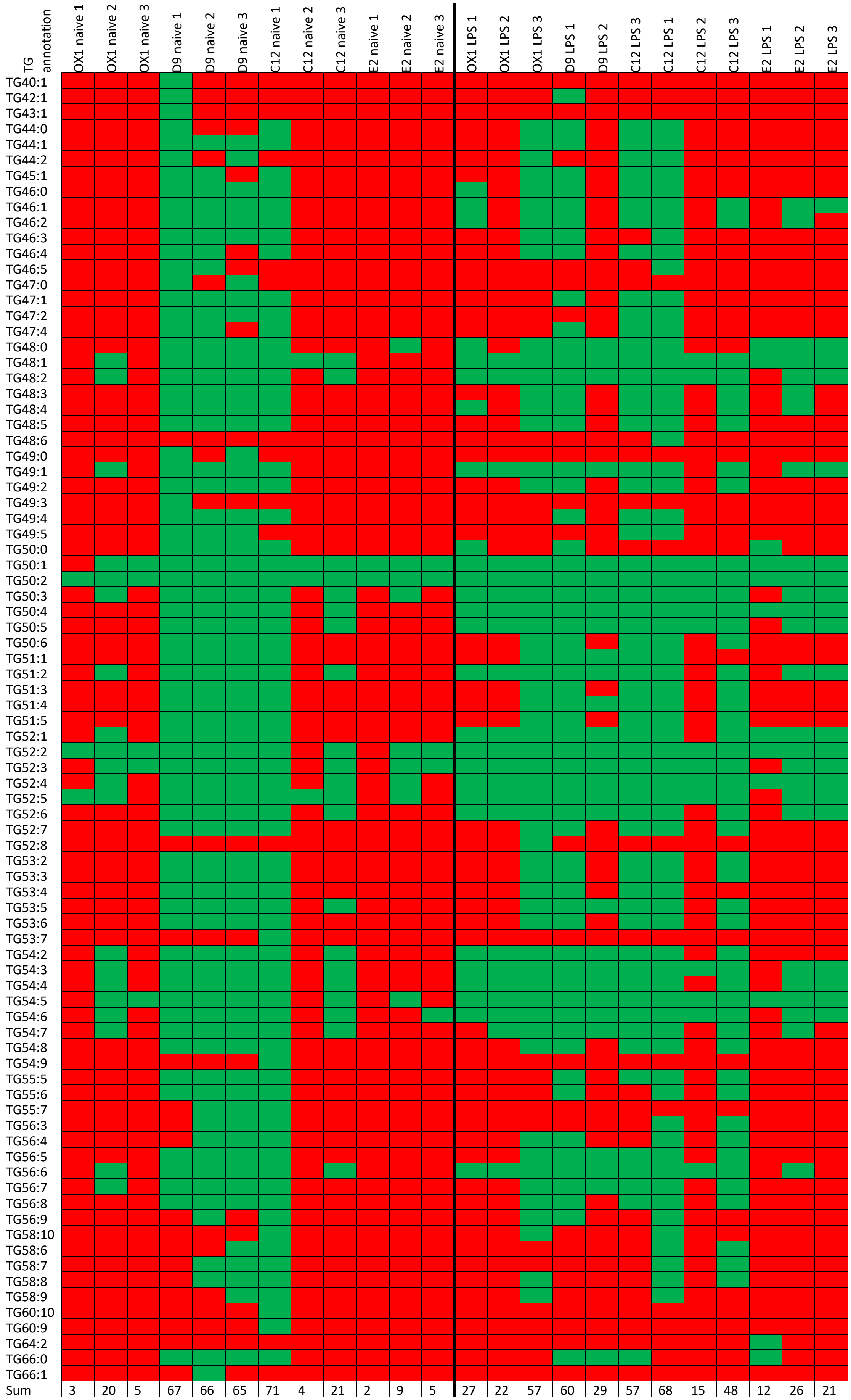


*Supporting Figure 16: Detected triglyceride species in different cell lines. Green indicates detection, red indicates no detection in Metaspace using LIPID MAPS database.*

| Cell line | Naïve | LPS |
|---|---|---|
| OX1 | 3 | 30 |
| | 22 | 24 |
| | 5 | 70 |
| D9 | 77 | 71 |
| | 77 | 33 |
| | 78 | 64 |
| C12 | 90 | 84 |
| | 4 | 15 |
| | 22 | 54 |
| E2 | 2 | 12 |
| | 9 | 28 |
| | 5 | 21 |

*Supporting Figure 17: Number of signals annotated as triglycerides by Metaspace using the LIPID MAPS database and a false discovery rate of 20% in different cell lines in naïve and LPS-stimulated state for biological triplicates, respectively. Highest values per replicate indicated in red. All measurements were carried out in positive-ion mode with 150x150 pixels using 5 µm pixel size.*

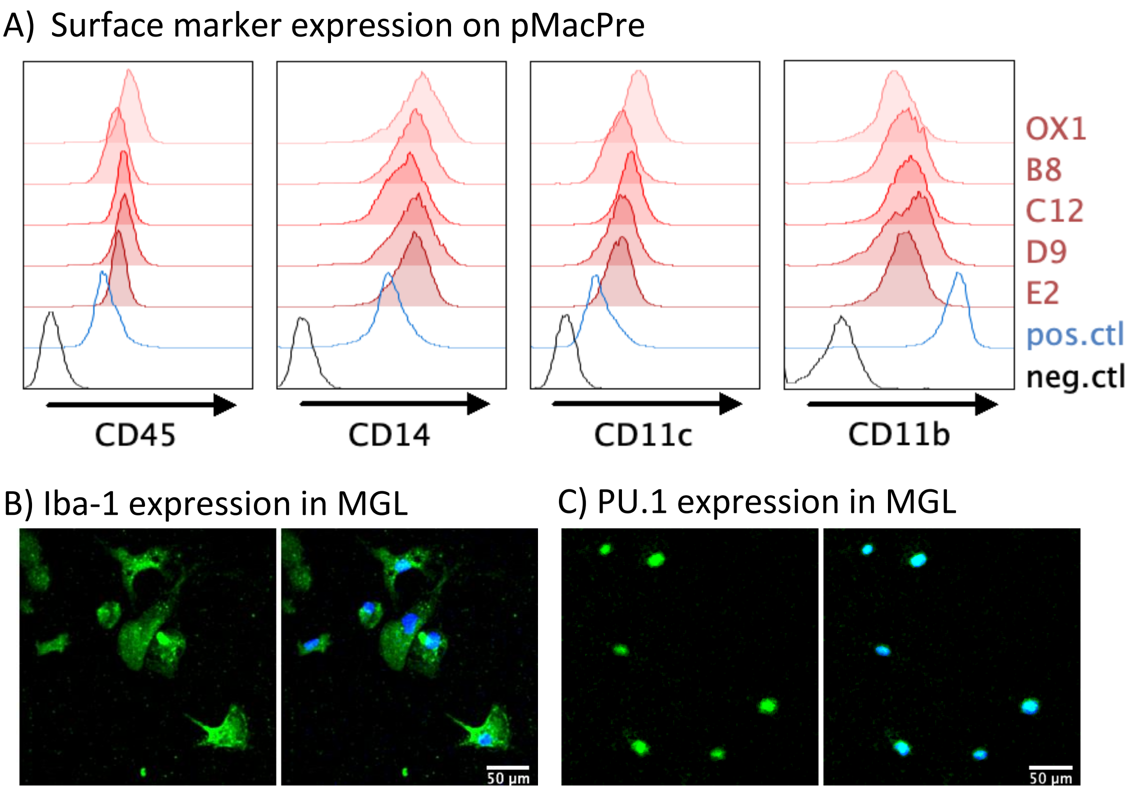


*Supporting Figure 18: A) Cell surface expression of immune cell marker CD45 and myeloid markers CD14, CD11c and, CD11b in iPSC-derived primitive macrophage precursor (pMacPre) is confirmed by flow cytometry. Negative control: iPSC; positive control: primary blood-derived monocytes. B,C) Representative images of intracellular myeloid markers Iba-1 (green, B) and PU.1 (green, C) in fully differentiated OX1 microglia-like cells. Nuclei were counterstained with DAPI (blue).*

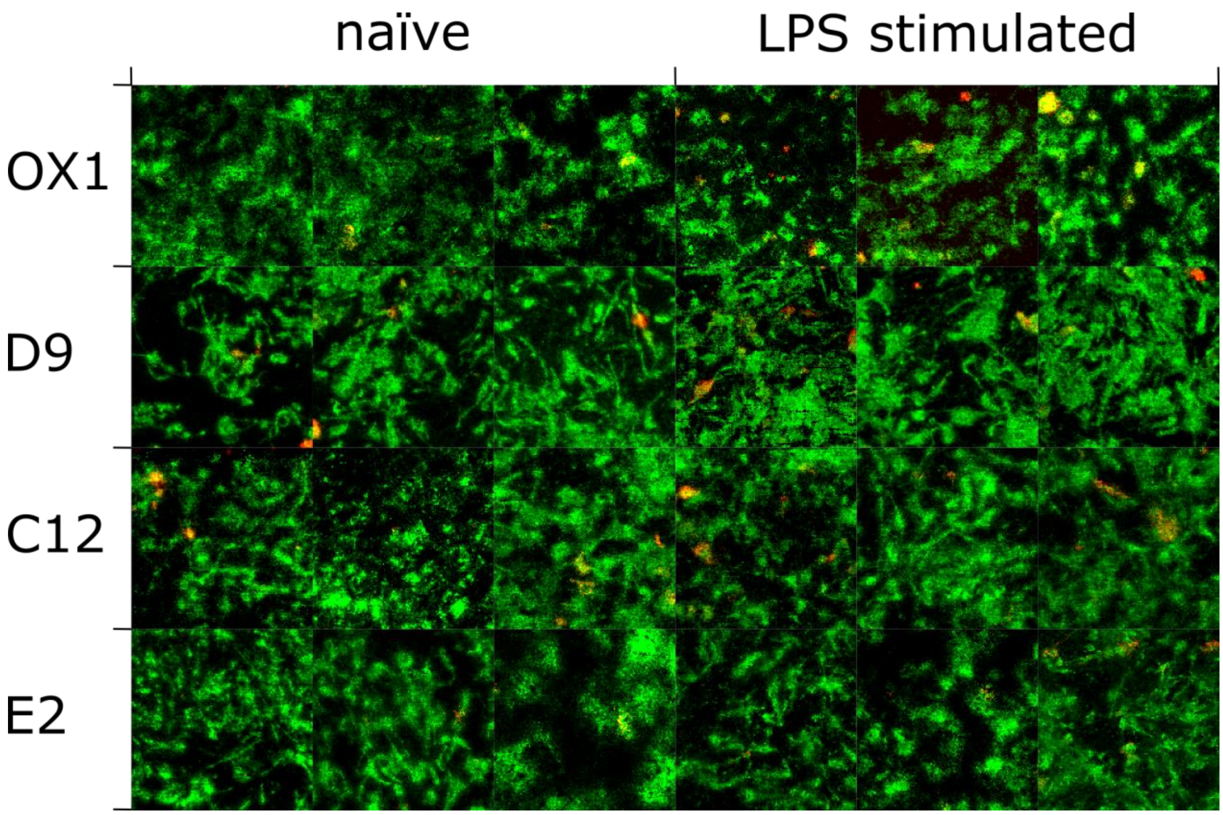


*Supporting Figure 19: 24 stitched MALDI MSI images, each 150x150 pixels at 5 µm pixel size, in positive-ion mode of biological triplicates of different cell lines in naïve or LPS-stimulated state showing the distribution of lipid droplets, represented by [TG 52:2 + Na]$^+$ (m/z 881.7569, red), compared to the cellular background, represented by [PC 36:2 + Na]$^+$ (m/z 808.5827, green).*

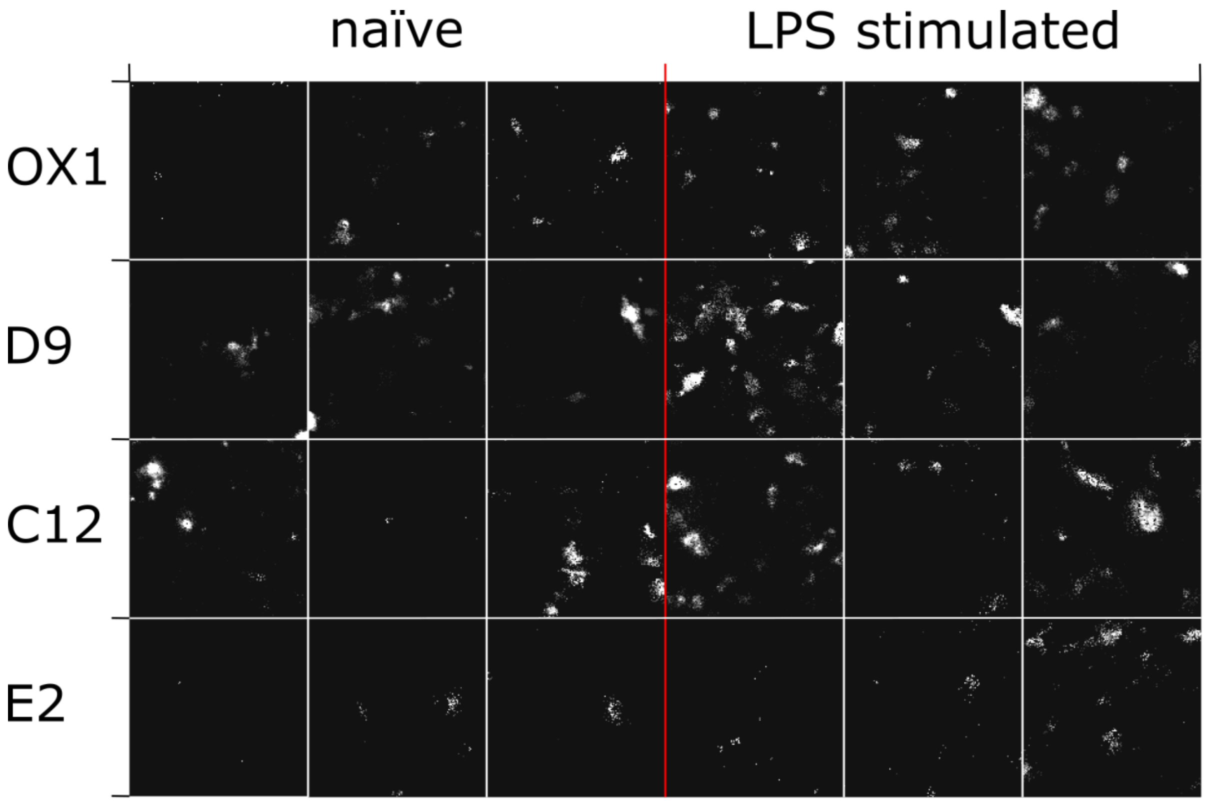


*Supporting Figure 20: 24 stitched MALDI MSI images, each 150x150 pixels at 5 µm pixel size, of [TG 52:2 + Na]$^+$ in biological triplicates of different cell lines in naïve or LPS-stimulated state. Red line indicates border between naïve and LPS-stimulated cells. White lines indicate borders between different measurements.*

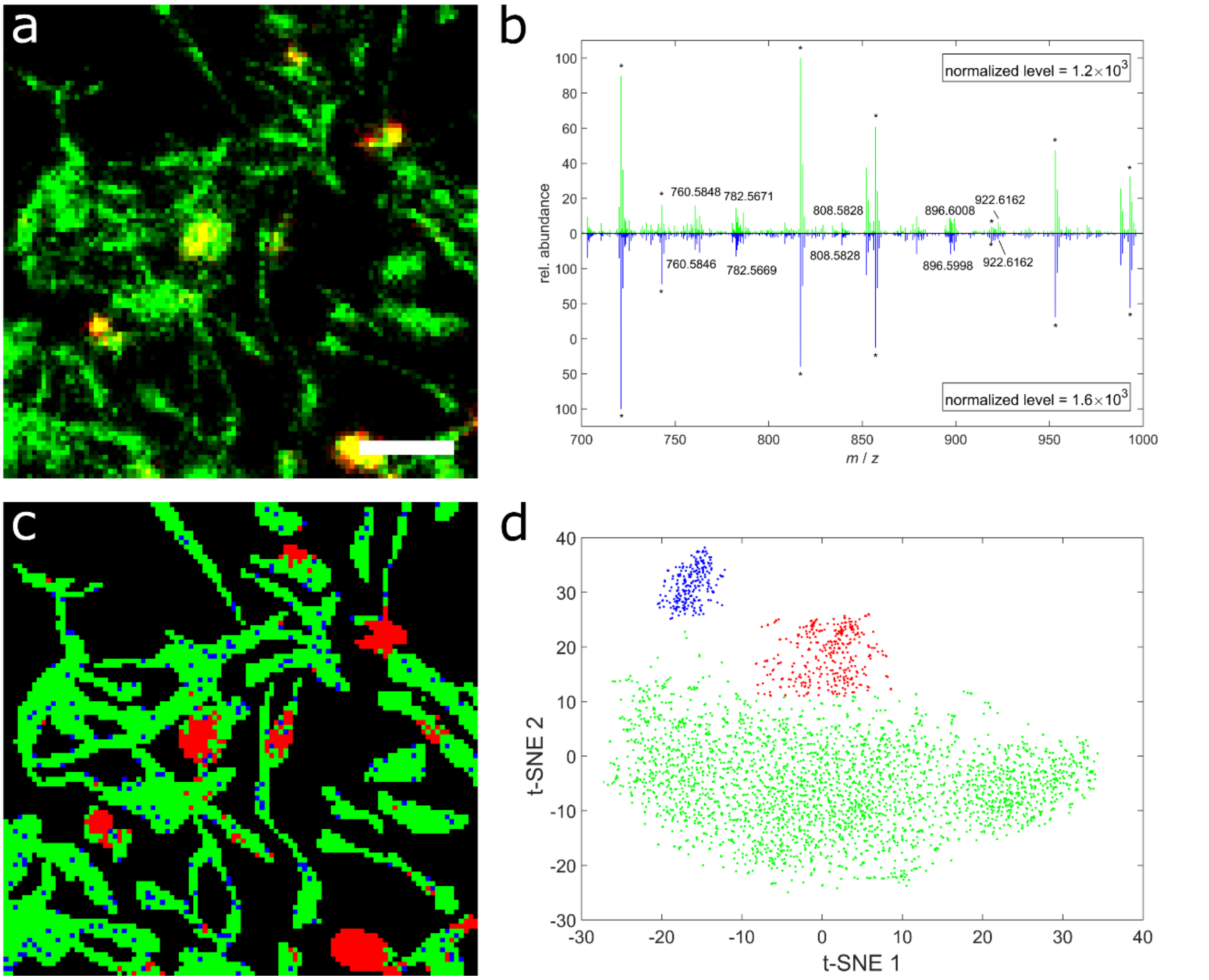


*Supporting Figure 21: MALDI MSI and statistical analysis for identification of cells with increased TG production.* ***a*** *MALDI MSI image with 100x100 pixels at 5 µm pixel size of fixed LPS-stimulated B8 MGL cells. Color-coding: red: TG 52:2 (m/z 881.7568), green: PC 34:1 (m/z 896.6140)* ***b*** *Example of single-pixel mass spectra corresponding to the green or blue clusters as identified by pixel-based t-SNE analysis in d. Stars indicate matrix-related signals.* ***c*** *Pixel-wise classification based on t-SNE analysis of the mass spectrometric data with automated color-coding referring to clusters identified in d.* ***d*** *Plot of pixel-wise t-SNE analysis of mass spectrometric data (m/z 800-1000) allowing for 3 clusters, while each of the 6511 data points represents an individual mass spectrum. The green cluster nicely outlines the cells in c, the red cluster identifies pixels with increased triglyceride production and the blue cluster is most likely an artifact of statistical analysis, since mass spectra as presented in b are virtually identical. Off-sample signals were excluded from analysis.*

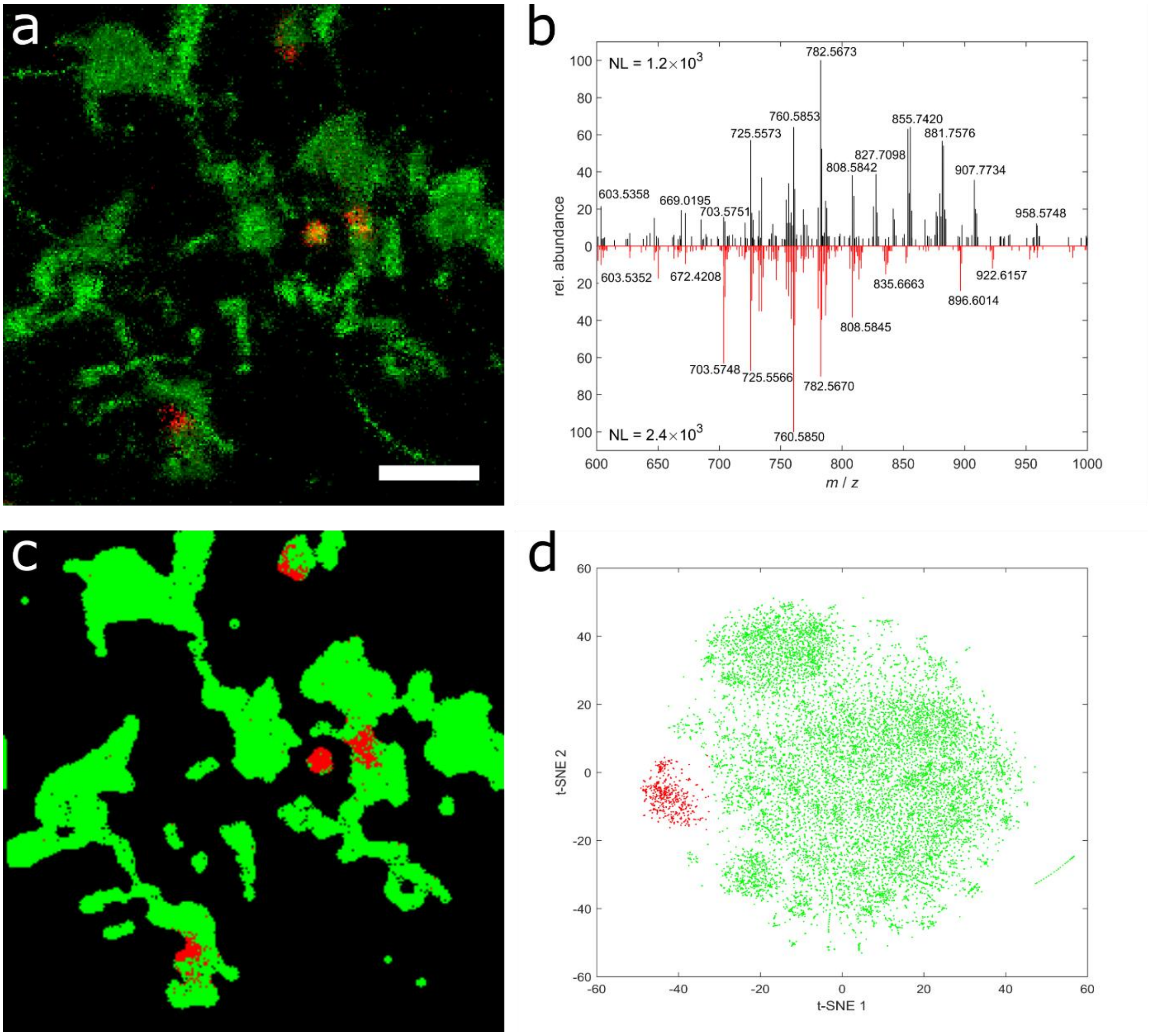


*Supporting Figure 22 MALDI MSI and statistical analysis for identification of cells with increased TG production.* ***a*** *MALDI MSI image with 250x250 pixels at 2 µm pixel size of naïve D9 cells. Color-coding: red: TG 52:2 (m/z 881.7568), green: PC 36:2 (m/z 808.5827)* ***b*** *Example of single-pixel mass spectra from pixels identified as TG-upregulated (black) or TG-downregulated (red).* ***c*** *Pixel-wise classification based on t-SNE analysis of the mass spectrometric data with automated color-coding referring to clusters identified in d.* ***d*** *Plot of pixel-wise t-SNE analysis of mass spectrometric data (m/z 800-1000) color-coded for clusters identified by hierarchical clustering, while each of the 48041 data points represents an individual mass spectrum. Off-sample signals were excluded from analysis.*

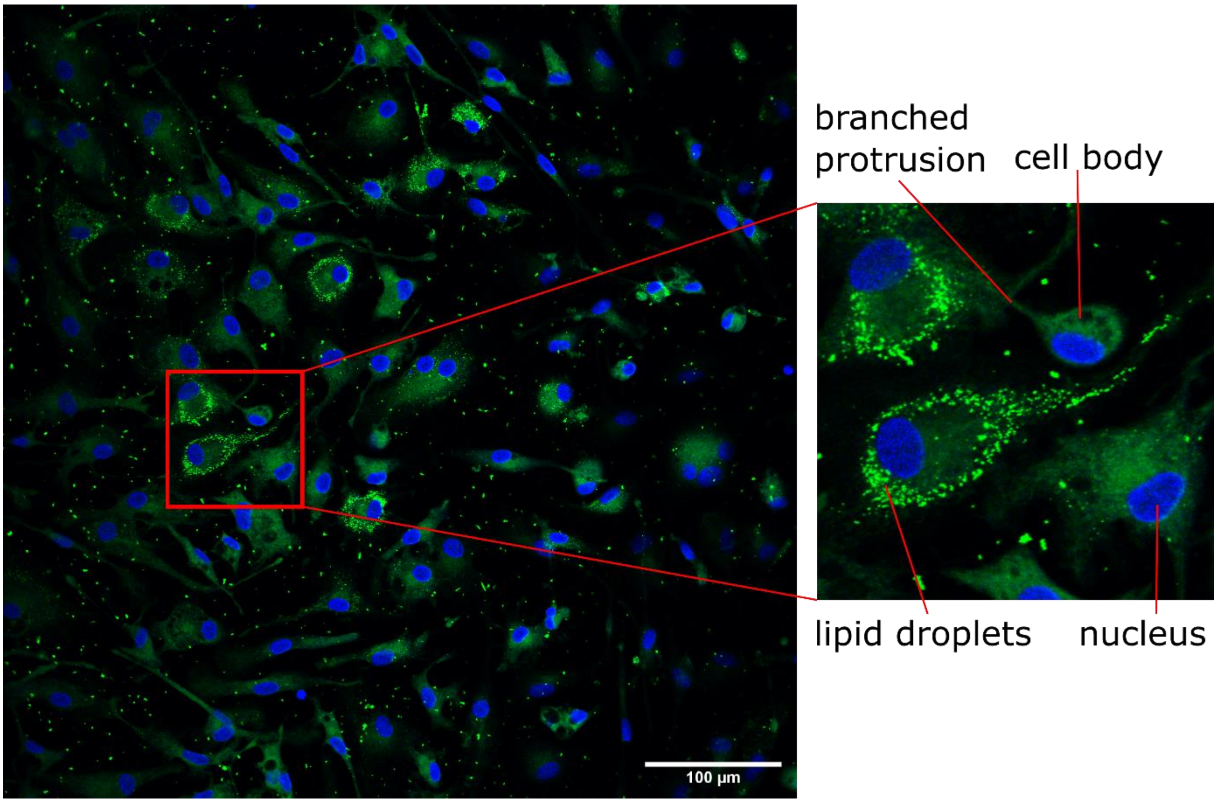


*Supporting Figure 23: Fluorescence microscopy image of naïve MGL cells of the B8 cell line, stained with BODIPY (green, selective for apolar domains, e.g. lipid droplets) and DAPI (blue, selective for DNA, e.g. cell nuclei), indicating a heterogeneous distribution of lipid droplets. Red lines indicate zoom-in on interesting area.*

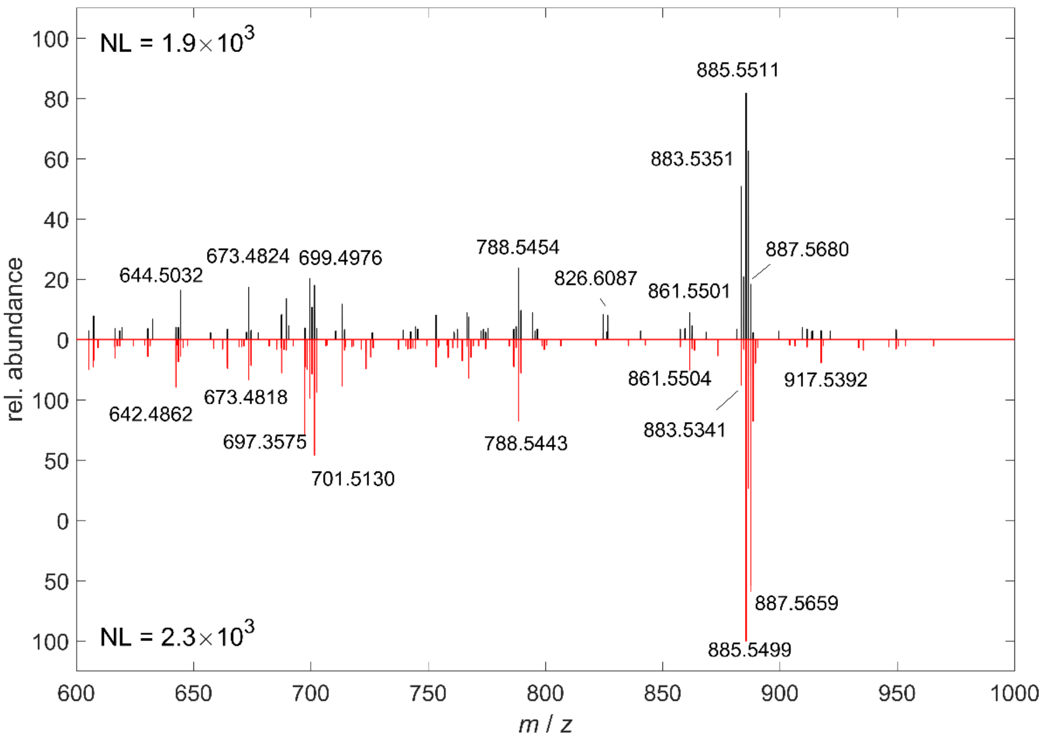


*Supporting Figure 24: Single-pixel mass spectra of different subpopulations in naïve B8 MGL cells measured in negative-ion mode.*

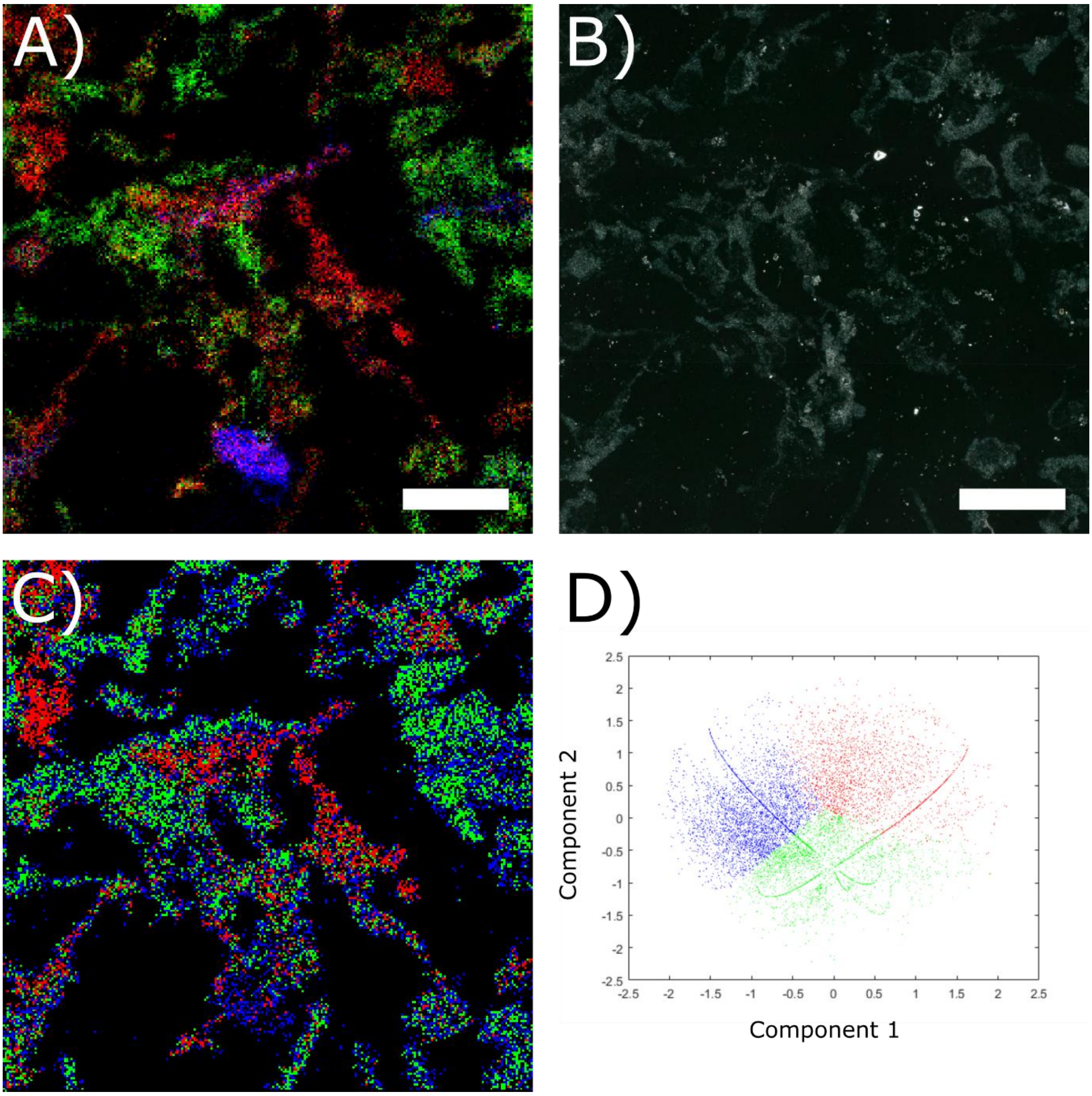


*Supporting Figure 25: Statistical analysis of a naïve sample from the B8 microglia cell line. A): MALDI MSI image showing [PI 38:3 - H]$^-$ (red, m/z 887.5655), [PI 38:5- H]$^-$ (green, m/z 883.5342) and [PI 36:1 - H]$^-$ (blue, m/z 863.5491). B): Corresponding microscopic image of the area investigated. C): Pixelwise classification using a k-means algorithm for automated clustering based on mass spectrometric data from m/z 860 – 900. Colors red, green and blue symbolize cluster affiliation. D): Principal component analysis plot of mass spectrometric data, colorized corresponding to clusters identified by k-means algorithm (as shown in C)). Each of the 46051 data points represents an individual mass spectrum. Clusters found by k-means also cluster in principal component analysis.*

*Supporting Table 1: List of annotations for signals identified as statistically different (ANOVA multiple-sample test, p-value ≤ 0.01) between the OX1, D9, C12 and E2 cell line (biological triplicates, each consisting of 22,500 individual mass spectra of multiple individual cells) in naïve state in positive-ion mode from LIPID MAPS database research with the lowest mass difference for each input mass.*

| Input mass | Matched mass | Δ *m/z* | Name | Formula | Ion |
|---|---|---|---|---|---|
| 870.58558 | 870.5854 | 0.0001 | DGCC 42:10 | C52H81NO8 | [M+Na]+ |
| 839.590695 | 839.5925 | 0.0018 | PA O-46:7 | C49H85O7P | [M+Na]+ |
| 871.588784 | 871.5847 | 0.004 | PG O-45:10 | C51H83O9P | [M+H]+ |
| 840.59397 | 840.5902 | 0.0038 | PE O-45:10 | C50H82NO7P | [M+H]+ |
| 841.607066 | 841.6082 | 0.0011 | PA O-46:6 | C49H87O7P | [M+Na]+ |
| 856.569217 | 856.5698 | 0.0006 | DGCC 41:10 | C51H79NO8 | [M+Na]+ |
| 842.609823 | 842.6058 | 0.004 | PE O-45:9 | C50H84NO7P | [M+H]+ |
| 977.621906 | 977.6242 | 0.0023 | PG 50:10 | C56H91O10P | [M+Na]+ |
| 842.55402 | 842.5541 | 0.0001 | DGCC 40:10 | C50H77NO8 | [M+Na]+ |
| 923.684503 | 923.6864 | 0.0019 | PA O-52:7 | C55H97O7P | [M+Na]+ |
| 622.44437 | 622.4442 | 0.0001 | PE 27:0 | C32H64NO8P | [M+H]+ |
| 743.540718 | 743.5375 | 0.0032 | DG O-44:9 | C47H76O4 | [M+K]+ |
| 872.591666 | 872.593 | 0.0013 | PE O-44:6 | C49H88NO7P | [M+K]+ |
| 566.309161 | 566.3089 | 0.0003 | PS 20:1 | C26H48NO10P | [M+H]+ |
| 857.572377 | 857.5691 | 0.0033 | TG 52:14 | C55H78O6 | [M+Na]+ |
| 869.637305 | 869.6395 | 0.0022 | PA O-48:6 | C51H91O7P | [M+Na]+ |
| 636.459937 | 636.4599 | 0.0001 | PE 28:0 | C33H66NO8P | [M+H]+ |
| 922.470795 | 922.4995 | 0.0287 | PS 44:10 | C50H78NO10P | [M+K]+ |
| 843.612753 | 843.611 | 0.0018 | PG 41:3 | C47H87O10P | [M+H]+ |
| 843.557386 | 843.5534 | 0.0039 | PG O-43:10 | C49H79O9P | [M+H]+ |
| 924.687991 | 924.6841 | 0.0039 | PE O-51:10 | C56H94NO7P | [M+H]+ |
| 608.428754 | 608.4286 | 0.0002 | PE 26:0 | C31H62NO8P | [M+H]+ |
| 594.340368 | 594.3403 | 0.0001 | DGCC 20:2 | C30H53NO8 | [M+K]+ |
| 839.696444 | 839.6912 | 0.0053 | DG O-54:12 | C57H90O4 | [M+H]+ |
| 925.699112 | 925.7021 | 0.0029 | PA O-52:6 | C55H99O7P | [M+Na]+ |
| 870.640934 | 870.6371 | 0.0038 | PE O-47:9 | C52H88NO7P | [M+H]+ |
| 754.519755 | 754.5228 | 0.0031 | DGCC 33:5 | C43H73NO8 | [M+Na]+ |
| 923.474614 | 923.4681 | 0.0065 | PI 40:11 | C49H73O13P | [M+Na]+ |
| 727.558935 | 727.5612 | 0.0023 | PA O-37:0 | C40H81O7P | [M+Na]+ |
| 760.566716 | 760.5698 | 0.0031 | DGCC 33:2 | C43H79NO8 | [M+Na]+ |
| 623.447803 | 623.4436 | 0.0042 | DG O-35:6 | C38H64O4 | [M+K]+ |
| 637.463337 | 637.4593 | 0.0041 | DG O-36:6 | C39H66O4 | [M+K]+ |
| 926.703562 | 926.6997 | 0.0038 | PE O-51:9 | C56H96NO7P | [M+H]+ |
| 840.700758 | 840.6841 | 0.0167 | PE O-44:3 | C49H94NO7P | [M+H]+ |
| 678.373253 | 678.3741 | 0.0009 | PS O-28:5 | C34H58NO9P | [M+Na]+ |
| 533.38227 | 533.3755 | 0.0067 | FA 34:7 | C34H54O2 | [M+K]+ |
| 813.575563 | 813.5769 | 0.0013 | PA O-44:6 | C47H83O7P | [M+Na]+ |
| 722.555096 | 722.5541 | 0.001 | DGCC 30:0 | C40H77NO8 | [M+Na]+ |
| 678.354409 | 678.3532 | 0.0012 | PE 29:5 | C34H58NO8P | [M+K]+ |
| 908.455188 | 908.4838 | 0.0287 | PS 43:10 | C49H76NO10P | [M+K]+ |
| 855.621178 | 855.6238 | 0.0026 | PA O-47:6 | C50H89O7P | [M+Na]+ |

| | | | | | |
|---|---|---|---|---|---|
| 609.432099 | 609.428 | 0.0041 | MG 34:6 | C37H62O4 | [M+K]+ |
| 897.668305 | 897.6708 | 0.0025 | PA O-50:6 | C53H95O7P | [M+Na]+ |
| 728.411316 | 728.4052 | 0.0061 | PE O-34:8 | C39H64NO7P | [M+K]+ |
| 844.561847 | 844.5617 | 0.0002 | PE O-42:6 | C47H84NO7P | [M+K]+ |
| 827.591684 | 827.5925 | 0.0008 | PA O-45:6 | C48H85O7P | [M+Na]+ |
| 931.506968 | 931.5097 | 0.0028 | PI O-40:8 | C49H81O12P | [M+K]+ |
| 757.592101 | 757.5824 | 0.0097 | MGDG 34:1 | C43H80O10 | [M+H]+ |
| 607.390803 | 607.3969 | 0.0061 | TG 33:6 | C36H56O6 | [M+Na]+ |
| 609.382905 | 609.3891 | 0.0062 | PA O-29:3 | C32H59O7P | [M+Na]+ |
| 894.475639 | 894.4682 | 0.0074 | PS 42:10 | C48H74NO10P | [M+K]+ |
| 723.557559 | 723.5558 | 0.0017 | TG 43:7 | C46H74O6 | [M+H]+ |
| 834.67897 | 834.6793 | 0.0004 | DGCC 38:0 | C48H93NO8 | [M+Na]+ |
| 798.490902 | 798.4834 | 0.0075 | PE O-39:8 | C44H74NO7P | [M+K]+ |
| 830.730248 | 830.6997 | 0.0305 | PE O-43:1 | C48H96NO7P | [M+H]+ |
| 772.617936 | 772.6215 | 0.0035 | PE O-39:2 | C44H86NO7P | [M+H]+ |
| 909.459382 | 909.4524 | 0.0069 | PI 39:11 | C48H71O13P | [M+Na]+ |
| 871.644404 | 871.6423 | 0.0021 | PG 43:3 | C49H91O10P | [M+H]+ |
| 884.604911 | 884.6011 | 0.0038 | DGCC 43:10 | C53H83NO8 | [M+Na]+ |
| 1069.58098 | 1069.5777 | 0.0033 | PIP 46:9 | C55H90O16P2 | [M+H]+ |
| 880.423294 | 880.4525 | 0.0292 | PS 41:10 | C47H72NO10P | [M+K]+ |
| 567.30977 | 567.3081 | 0.0016 | PA 27:6 | C30H47O8P | [M+H]+ |
| 996.73133 | 996.7263 | 0.005 | DGCC 51:10 | C61H99NO8 | [M+Na]+ |
| 968.701536 | 968.7079 | 0.0063 | PE 51:6 | C56H100NO8P | [M+Na]+ |
| 908.49158 | 908.4838 | 0.0077 | PS 43:10 | C49H76NO10P | [M+K]+ |
| 927.706895 | 927.7049 | 0.002 | PG 47:3 | C53H99O10P | [M+H]+ |
| 922.507102 | 922.4995 | 0.0076 | PS 44:10 | C50H78NO10P | [M+K]+ |
| 946.469897 | 946.4995 | 0.0296 | PS 46:12 | C52H78NO10P | [M+K]+ |
| 1131.71183 | 1131.7141 | 0.0022 | SQDG 56:11 | C65H104O12S | [M+Na]+ |
| 1089.61048 | 1089.6098 | 0.0007 | SQDG 52:12 | C61H94O12S | [M+K]+ |

*Supporting Table 2: List of annotations for signals identified as statistically different (ANOVA multiple-sample test, p-value ≤ 0.01) between the OX1, D9, C12 and E2 cell line (biological triplicates) in LPS-stimulated state in positive-ion mode from LIPID MAPS database research with the lowest mass difference for each input mass.*

| Input mass | Matched mass | Δ $m/z$ | Name | Formula | Ion |
|---|---|---|---|---|---|
| 734.56975 | 734.5694 | 0.0003 | PE 35:0 | C40H80NO8P | [M+H]+ |
| 870.58558 | 870.5854 | 0.0001 | DGCC 42:10 | C52H81NO8 | [M+Na]+ |
| 757.555242 | 757.5532 | 0.0021 | DG O-45:9 | C48H78O4 | [M+K]+ |
| 786.601077 | 786.6007 | 0.0003 | PE 39:2 | C44H84NO8P | [M+H]+ |
| 735.572987 | 735.5688 | 0.0042 | DG O-43:6 | C46H80O4 | [M+K]+ |
| 839.590695 | 839.5925 | 0.0018 | PA O-46:7 | C49H85O7P | [M+Na]+ |
| 758.570202 | 758.5694 | 0.0008 | PE 37:2 | C42H80NO8P | [M+H]+ |
| 697.477913 | 697.4779 | 0 | PA 34:1 | C37H71O8P | [M+Na]+ |
| 742.535415 | 742.5357 | 0.0003 | PE 34:0 | C39H78NO8P | [M+Na]+ |
| 667.486894 | 667.4908 | 0.0039 | TG 37:4 | C40H68O6 | [M+Na]+ |
| 871.588784 | 871.5847 | 0.004 | PG O-45:10 | C51H83O9P | [M+H]+ |

| | | | | | |
|---|---|---|---|---|---|
| 720.55378 | 720.5538 | 0 | PE 34:0 | C39H78NO8P | [M+H]+ |
| 840.59397 | 840.5902 | 0.0038 | PE O-45:10 | C50H82NO7P | [M+H]+ |
| 759.573181 | 759.5688 | 0.0044 | DG O-45:8 | C48H80O4 | [M+K]+ |
| 746.56951 | 746.5694 | 0.0001 | PE 36:1 | C41H80NO8P | [M+H]+ |
| 756.551877 | 756.5514 | 0.0005 | PE 35:0 | C40H80NO8P | [M+Na]+ |
| 676.454818 | 676.4548 | 0 | PS O-29:2 | C35H66NO9P | [M+H]+ |
| 706.538203 | 706.5381 | 0.0001 | PE 33:0 | C38H76NO8P | [M+H]+ |
| 787.604343 | 787.6001 | 0.0042 | DG O-47:8 | C50H84O4 | [M+K]+ |
| 841.607066 | 841.6082 | 0.0011 | PA O-46:6 | C49H87O7P | [M+Na]+ |
| 720.59019 | 720.5902 | 0 | PE O-35:0 | C40H82NO7P | [M+H]+ |
| 728.520166 | 728.5201 | 0.0001 | PE 33:0 | C38H76NO8P | [M+Na]+ |
| 842.609823 | 842.6058 | 0.004 | PE O-45:9 | C50H84NO7P | [M+H]+ |
| 747.57291 | 747.5688 | 0.0041 | DG O-44:7 | C47H80O4 | [M+K]+ |
| 762.591071 | 762.5878 | 0.0032 | DGCC 35:4 | C45H79NO8 | [M+H]+ |
| 730.538104 | 730.5381 | 0 | PE 35:2 | C40H76NO8P | [M+H]+ |
| 812.541415 | 812.5412 | 0.0002 | PS 36:1 | C42H80NO10P | [M+Na]+ |
| 721.557226 | 721.5532 | 0.0041 | DG O-42:6 | C45H78O4 | [M+K]+ |
| 772.584822 | 772.5851 | 0.0003 | PE 38:2 | C43H82NO8P | [M+H]+ |
| 748.585319 | 748.5851 | 0.0002 | PE 36:0 | C41H82NO8P | [M+H]+ |
| 698.481372 | 698.4755 | 0.0058 | PE 33:4 | C38H68NO8P | [M+H]+ |
| 977.621906 | 977.6242 | 0.0023 | PG 50:10 | C56H91O10P | [M+Na]+ |
| 677.458343 | 677.4542 | 0.0042 | TG O-38:7 | C41H66O5 | [M+K]+ |
| 932.559707 | 932.5566 | 0.0031 | PE 48:11 | C53H84NO8P | [M+K]+ |
| 842.55402 | 842.5541 | 0.0001 | DGCC 40:10 | C50H77NO8 | [M+Na]+ |
| 923.684503 | 923.6864 | 0.0019 | PA O-52:7 | C55H97O7P | [M+Na]+ |
| 707.54418 | 707.5375 | 0.0067 | DG O-41:6 | C44H76O4 | [M+K]+ |
| 856.605845 | 856.6062 | 0.0003 | DGTS 42:9 | C52H83NO7 | [M+Na]+ |
| 729.523732 | 729.5219 | 0.0019 | DG O-43:9 | C46H74O4 | [M+K]+ |
| 683.461549 | 683.4622 | 0.0007 | PA 33:1 | C36H69O8P | [M+Na]+ |
| 743.540718 | 743.5375 | 0.0032 | DG O-44:9 | C47H76O4 | [M+K]+ |
| 957.565559 | 957.5732 | 0.0077 | SQDG 43:7 | C52H86O12S | [M+Na]+ |
| 721.593556 | 721.5894 | 0.0042 | CE 22:5 | C49H78O2 | [M+Na]+ |
| 767.539472 | 767.5375 | 0.002 | DG O-46:11 | C49H76O4 | [M+K]+ |
| 736.575907 | 736.5722 | 0.0037 | DGCC 33:3 | C43H77NO8 | [M+H]+ |
| 675.543594 | 675.5477 | 0.0041 | CE 17:1 | C44H76O2 | [M+K]+ |
| 872.591666 | 872.593 | 0.0013 | PE O-44:6 | C49H88NO7P | [M+K]+ |
| 743.575672 | 743.5668 | 0.0089 | MGDG 33:1 | C42H78O10 | [M+H]+ |
| 771.571366 | 771.5688 | 0.0026 | DG O-46:9 | C49H80O4 | [M+K]+ |
| 867.621901 | 867.6238 | 0.0019 | PA O-48:7 | C51H89O7P | [M+Na]+ |
| 731.541639 | 731.5375 | 0.0041 | DG O-43:8 | C46H76O4 | [M+K]+ |
| 956.559706 | 956.5777 | 0.018 | PS 46:7 | C52H88NO10P | [M+K]+ |
| 728.558438 | 728.5589 | 0.0004 | PE O-36:3 | C41H78NO7P | [M+H]+ |
| 682.457135 | 682.4572 | 0.0001 | LPE O-31:3 | C36H70NO6P | [M+K]+ |
| 978.625681 | 978.6219 | 0.0038 | PS 51:12 | C57H88NO10P | [M+H]+ |
| 931.549432 | 931.546 | 0.0035 | PG 47:12 | C53H81O10P | [M+Na]+ |

| | | | | | |
|---|---|---|---|---|---|
| 714.54097 | 714.5408 | 0.0002 | PE O-33:0 | C38H78NO7P | [M+Na]+ |
| 726.50467 | 726.5044 | 0.0002 | PE 33:1 | C38H74NO8P | [M+Na]+ |
| 1069.57888 | 1069.5777 | 0.0012 | PIP 46:9 | C55H90O16P2 | [M+H]+ |
| 869.637305 | 869.6395 | 0.0022 | PA O-48:6 | C51H91O7P | [M+Na]+ |
| 843.612753 | 843.611 | 0.0018 | PG 41:3 | C47H87O10P | [M+H]+ |
| 669.446555 | 669.4466 | 0 | PA 32:1 | C35H67O8P | [M+Na]+ |
| 773.58841 | 773.5845 | 0.0039 | DG O-46:8 | C49H82O4 | [M+K]+ |
| 704.522522 | 704.5225 | 0 | PE 33:1 | C38H74NO8P | [M+H]+ |
| 788.607085 | 788.6035 | 0.0036 | DGCC 37:5 | C47H81NO8 | [M+H]+ |
| 692.558754 | 692.5589 | 0.0001 | PE O-33:0 | C38H78NO7P | [M+H]+ |
| 810.525656 | 810.5256 | 0.0001 | PS 36:2 | C42H78NO10P | [M+Na]+ |
| 813.544702 | 813.543 | 0.0017 | TG 47:9 | C50H78O6 | [M+K]+ |
| 546.257745 | 546.2593 | 0.0015 | LPS O-18:2 | C24H46NO8P | [M+K]+ |
| 738.50479 | 738.5044 | 0.0004 | PE 34:2 | C39H74NO8P | [M+Na]+ |
| 843.557386 | 843.5534 | 0.0039 | PG O-43:10 | C49H79O9P | [M+H]+ |
| 749.588689 | 749.5845 | 0.0042 | DG O-44:6 | C47H82O4 | [M+K]+ |
| 775.604256 | 775.6001 | 0.0041 | DG O-46:7 | C49H84O4 | [M+K]+ |
| 857.608949 | 857.6113 | 0.0024 | MGDG 40:4 | C49H86O10 | [M+Na]+ |
| 548.334807 | 548.3347 | 0.0001 | PE 22:2 | C27H50NO8P | [M+H]+ |
| 933.563955 | 933.5616 | 0.0024 | PG 47:11 | C53H83O10P | [M+Na]+ |
| 739.467662 | 739.4675 | 0.0002 | PA 36:2 | C39H73O8P | [M+K]+ |
| 762.503752 | 762.5044 | 0.0007 | PE 36:4 | C41H74NO8P | [M+Na]+ |
| 909.545752 | 909.5463 | 0.0006 | PI 38:4 | C47H83O13P | [M+Na]+ |
| 714.504775 | 714.5044 | 0.0004 | PE 32:0 | C37H74NO8P | [M+Na]+ |
| 764.520426 | 764.5201 | 0.0004 | PE 36:3 | C41H76NO8P | [M+Na]+ |
| 770.53063 | 770.5306 | 0 | PS O-34:1 | C40H78NO9P | [M+Na]+ |
| 835.526799 | 835.5272 | 0.0004 | PA 47:12 | C50H75O8P | [M+H]+ |
| 542.491631 | 542.4907 | 0.0009 | NAE 32:2 | C34H65NO2 | [M+Na]+ |
| 899.61828 | 899.616 | 0.0022 | PG O-47:10 | C53H87O9P | [M+H]+ |
| 784.50989 | 784.5099 | 0 | PS 34:1 | C40H76NO10P | [M+Na]+ |
| 825.559736 | 825.5616 | 0.0019 | PG 38:2 | C44H83O10P | [M+Na]+ |
| 868.625368 | 868.6215 | 0.0039 | PE O-47:10 | C52H86NO7P | [M+H]+ |
| 925.699112 | 925.7021 | 0.0029 | PA O-52:6 | C55H99O7P | [M+Na]+ |
| 898.610077 | 898.6086 | 0.0014 | PE O-46:7 | C51H90NO7P | [M+K]+ |
| 1151.6151 | 1151.5962 | 0.0189 | PIP 49:8 | C58H98O16P2 | [M+K]+ |
| 676.54713 | 676.5639 | 0.0168 | LPE O-33:1 | C38H78NO6P | [M+H]+ |
| 797.54965 | 797.5481 | 0.0016 | TG O-47:10 | C50H78O5 | [M+K]+ |
| 683.498514 | 683.4986 | 0.0001 | PA O-34:1 | C37H73O7P | [M+Na]+ |
| 799.565248 | 799.5637 | 0.0015 | TG O-47:9 | C50H80O5 | [M+K]+ |
| 592.337673 | 592.3375 | 0.0002 | LPS O-21:0 | C27H56NO8P | [M+K]+ |
| 832.507947 | 832.5099 | 0.002 | PS 38:5 | C44H76NO10P | [M+Na]+ |
| 1173.59693 | 1173.5805 | 0.0164 | PIP 51:11 | C60H96O16P2 | [M+K]+ |
| 1077.57316 | 1077.5805 | 0.0074 | PIP 43:3 | C52H96O16P2 | [M+K]+ |
| 789.521932 | 789.5195 | 0.0024 | PA O-41:5 | C44H79O7P | [M+K]+ |
| 870.640934 | 870.6371 | 0.0038 | PE O-47:9 | C52H88NO7P | [M+H]+ |

| | | | | | |
|---|---|---|---|---|---|
| 705.489613 | 705.4855 | 0.0041 | TG O-40:7 | C43H70O5 | [M+K]+ |
| 544.241977 | 544.2436 | 0.0016 | LPS O-18:3 | C24H44NO8P | [M+K]+ |
| 700.489251 | 700.4888 | 0.0005 | PE 31:0 | C36H72NO8P | [M+Na]+ |
| 774.564004 | 774.5643 | 0.0003 | PS O-36:2 | C42H80NO9P | [M+H]+ |
| 713.453091 | 713.4541 | 0.001 | PA O-39:10 | C42H65O7P | [M+H]+ |
| 729.560846 | 729.5511 | 0.0097 | MGDG 32:1 | C41H76O10 | [M+H]+ |
| 907.529322 | 907.5307 | 0.0014 | PI 38:5 | C47H81O13P | [M+Na]+ |
| 697.462327 | 697.465 | 0.0027 | LPI O-26:1 | C35H69O11P | [M+H]+ |
| 763.594411 | 763.6001 | 0.0057 | DG O-45:6 | C48H84O4 | [M+K]+ |
| 706.522742 | 706.5228 | 0.0001 | DGCC 29:1 | C39H73NO8 | [M+Na]+ |
| 688.465731 | 688.4676 | 0.0019 | LPE O-33:6 | C38H68NO6P | [M+Na]+ |
| 727.558935 | 727.5612 | 0.0023 | PA O-37:0 | C40H81O7P | [M+Na]+ |
| 711.493177 | 711.4935 | 0.0003 | PA 35:1 | C38H73O8P | [M+Na]+ |
| 845.674056 | 845.6784 | 0.0043 | DG O-51:7 | C54H94O4 | [M+K]+ |
| 507.249196 | 507.2484 | 0.0008 | LPG O-16:1 | C22H45O8P | [M+K]+ |
| 760.566716 | 760.5698 | 0.0031 | DGCC 33:2 | C43H79NO8 | [M+Na]+ |
| 944.561277 | 944.5566 | 0.0047 | PE 49:12 | C54H84NO8P | [M+K]+ |
| 1175.70202 | 1175.7007 | 0.0013 | DGDG 51:11 | C66H104O15 | [M+K]+ |
| 1153.62984 | 1153.6118 | 0.018 | PIP 49:7 | C58H100O16P2 | [M+K]+ |
| 802.595366 | 802.5956 | 0.0003 | PS O-38:2 | C44H84NO9P | [M+H]+ |
| 768.515252 | 768.515 | 0.0003 | PS O-34:2 | C40H76NO9P | [M+Na]+ |
| 712.489136 | 712.4888 | 0.0004 | PE 32:1 | C37H72NO8P | [M+Na]+ |
| 708.549286 | 708.5538 | 0.0045 | LPS O-32:0 | C38H78NO8P | [M+H]+ |
| 806.493042 | 806.4943 | 0.0012 | PS 36:4 | C42H74NO10P | [M+Na]+ |
| 811.559867 | 811.5612 | 0.0013 | PA O-44:7 | C47H81O7P | [M+Na]+ |
| 800.616174 | 800.6164 | 0.0002 | PE 40:2 | C45H86NO8P | [M+H]+ |
| 955.550793 | 955.5576 | 0.0068 | SQDG 43:8 | C52H84O12S | [M+Na]+ |
| 798.524458 | 798.5256 | 0.0011 | PS 35:1 | C41H78NO10P | [M+Na]+ |
| 705.526233 | 705.5219 | 0.0044 | DG O-41:7 | C44H74O4 | [M+K]+ |
| 1032.58997 | 1032.609 | 0.0191 | PS 52:11 | C58H92NO10P | [M+K]+ |
| 812.51978 | 812.5201 | 0.0003 | PE 40:7 | C45H76NO8P | [M+Na]+ |
| 673.409096 | 673.4075 | 0.0016 | PG 29:4 | C35H61O10P | [M+H]+ |
| 548.274177 | 548.2748 | 0.0006 | PE O-22:6 | C27H44NO7P | [M+Na]+ |
| 737.579243 | 737.5845 | 0.0052 | DG O-43:5 | C46H82O4 | [M+K]+ |
| 926.703562 | 926.6997 | 0.0038 | PE O-51:9 | C56H96NO7P | [M+H]+ |
| 692.522441 | 692.5225 | 0 | PE 32:0 | C37H74NO8P | [M+H]+ |
| 648.472903 | 648.481 | 0.0081 | DGTS 26:1 | C36H67NO7 | [M+Na]+ |
| 765.483543 | 765.4831 | 0.0004 | PA 38:3 | C41H75O8P | [M+K]+ |
| 873.594668 | 873.598 | 0.0033 | PG O-43:6 | C49H87O9P | [M+Na]+ |
| 814.631821 | 814.632 | 0.0002 | PE 41:2 | C46H88NO8P | [M+H]+ |
| 788.603307 | 788.6035 | 0.0002 | DGCC 37:5 | C47H81NO8 | [M+H]+ |
| 572.27274 | 572.2749 | 0.0022 | PE 21:2 | C26H48NO8P | [M+K]+ |
| 768.371591 | 768.3847 | 0.0131 | PS 34:9 | C40H60NO10P | [M+Na]+ |
| 811.52917 | 811.5273 | 0.0018 | TG 47:10 | C50H76O6 | [M+K]+ |
| 765.523876 | 765.5219 | 0.002 | DG O-46:12 | C49H74O4 | [M+K]+ |

| | | | | | |
|---|---|---|---|---|---|
| 760.488275 | 760.4888 | 0.0005 | PE 36:5 | C41H72NO8P | [M+Na]+ |
| 500.311477 | 500.3111 | 0.0003 | LPE O-19:2 | C24H48NO6P | [M+Na]+ |
| 740.470681 | 740.4627 | 0.008 | PE 33:2 | C38H72NO8P | [M+K]+ |
| 910.548937 | 910.5569 | 0.0079 | PS 44:8 | C50H82NO10P | [M+Na]+ |
| 1133.57352 | 1133.5492 | 0.0243 | PIP 48:10 | C57H92O16P2 | [M+K]+ |
| 1113.63729 | 1113.6379 | 0.0006 | PIP 47:5 | C56H100O16P2 | [M+Na]+ |
| 814.535272 | 814.5357 | 0.0005 | PE 40:6 | C45H78NO8P | [M+Na]+ |
| 715.544491 | 715.5355 | 0.009 | MGDG 31:1 | C40H74O10 | [M+H]+ |
| 960.58326 | 960.5879 | 0.0047 | PE 50:11 | C55H88NO8P | [M+K]+ |
| 721.53995 | 721.5402 | 0.0002 | TG 43:8 | C46H72O6 | [M+H]+ |
| 1135.58771 | 1135.5649 | 0.0228 | PIP 48:9 | C57H94O16P2 | [M+K]+ |
| 895.567802 | 895.5671 | 0.0007 | PI O-38:4 | C47H85O12P | [M+Na]+ |
| 693.562406 | 693.5581 | 0.0043 | CE 20:5 | C47H74O2 | [M+Na]+ |
| 763.507182 | 763.5062 | 0.001 | DG O-46:13 | C49H72O4 | [M+K]+ |
| 954.544877 | 954.5621 | 0.0172 | PS 46:8 | C52H86NO10P | [M+K]+ |
| 739.509274 | 739.512 | 0.0027 | LPI O-29:1 | C38H75O11P | [M+H]+ |
| 724.480592 | 724.4783 | 0.0023 | DGCC 33:9 | C43H65NO8 | [M+H]+ |
| 1006.5742 | 1006.5934 | 0.0192 | PS 50:10 | C56H90NO10P | [M+K]+ |
| 799.526229 | 799.5272 | 0.001 | PA 44:9 | C47H75O8P | [M+H]+ |
| 711.437145 | 711.4362 | 0.001 | PA 34:2 | C37H69O8P | [M+K]+ |
| 604.526543 | 604.5275 | 0.001 | CAR 29:0 | C36H71NO4 | [M+Na]+ |
| 685.548731 | 685.5532 | 0.0044 | DG O-39:3 | C42H78O4 | [M+K]+ |
| 767.521447 | 767.5221 | 0.0007 | TG 45:10 | C48H72O6 | [M+Na]+ |
| 1095.56477 | 1095.5909 | 0.0262 | PIP 46:7 | C55H94O16P2 | [M+Na]+ |
| 547.262495 | 547.2643 | 0.0018 | BMP 18:1 | C24H45O10P | [M+Na]+ |
| 688.284483 | 688.3011 | 0.0166 | PS O-28:8 | C34H52NO9P | [M+K]+ |
| 748.548421 | 748.5487 | 0.0003 | PS O-34:1 | C40H78NO9P | [M+H]+ |
| 885.545642 | 885.5463 | 0.0007 | PI 36:2 | C45H83O13P | [M+Na]+ |
| 1349.60739 | 1349.6043 | 0.0031 | PIP3 55:12 | C64H104O22P4 | [M+H]+ |
| 725.508141 | 725.5092 | 0.001 | PA 36:1 | C39H75O8P | [M+Na]+ |
| 858.578295 | 858.5773 | 0.0009 | PE O-43:6 | C48H86NO7P | [M+K]+ |
| 737.473087 | 737.4728 | 0.0003 | PG O-33:4 | C39H71O9P | [M+Na]+ |
| 533.38227 | 533.3755 | 0.0067 | FA 34:7 | C34H54O2 | [M+K]+ |
| 1183.61346 | 1183.6199 | 0.0064 | PIP2 46:3 | C55H103O19P3 | [M+Na]+ |
| 647.448244 | 647.4436 | 0.0046 | DG O-37:8 | C40H64O4 | [M+K]+ |
| 979.628999 | 979.627 | 0.002 | PI 45:7 | C54H91O13P | [M+H]+ |
| 813.575563 | 813.5769 | 0.0013 | PA O-44:6 | C47H83O7P | [M+Na]+ |
| 856.505327 | 856.5099 | 0.0046 | PS 40:7 | C46H76NO10P | [M+Na]+ |
| 560.501548 | 560.4803 | 0.0212 | NAE 32:1 | C34H67NO2 | [M+K]+ |
| 722.564724 | 722.5694 | 0.0047 | LPS O-33:0 | C39H80NO8P | [M+H]+ |
| 699.48401 | 699.4807 | 0.0033 | LPI O-26:0 | C35H71O11P | [M+H]+ |
| 504.306458 | 504.3061 | 0.0004 | LPE 18:0 | C23H48NO7P | [M+Na]+ |
| 737.451828 | 737.4518 | 0 | PA 36:3 | C39H71O8P | [M+K]+ |
| 543.494927 | 543.4772 | 0.0178 | MG O-33:6 | C36H62O3 | [M+H]+ |
| 798.465841 | 798.468 | 0.0022 | PS O-37:8 | C43H70NO9P | [M+Na]+ |

| | | | | | |
|---|---|---|---|---|---|
| 739.488607 | 739.4884 | 0.0002 | PG O-33:3 | C39H73O9P | [M+Na]+ |
| 753.503733 | 753.5041 | 0.0004 | PG O-34:3 | C40H75O9P | [M+Na]+ |
| 771.534285 | 771.5324 | 0.0019 | TG O-45:9 | C48H76O5 | [M+K]+ |
| 733.521979 | 733.5168 | 0.0052 | TG O-42:7 | C45H74O5 | [M+K]+ |
| 833.511473 | 833.5117 | 0.0002 | TG 49:13 | C52H74O6 | [M+K]+ |
| 904.529469 | 904.5253 | 0.0042 | PE 46:11 | C51H80NO8P | [M+K]+ |
| 1107.55838 | 1107.5336 | 0.0248 | PIP 46:9 | C55H90O16P2 | [M+K]+ |
| 758.423668 | 758.4158 | 0.0079 | PE 35:7 | C40H66NO8P | [M+K]+ |
| 746.532892 | 746.533 | 0.0002 | PS O-34:2 | C40H76NO9P | [M+H]+ |
| 785.513337 | 785.5117 | 0.0016 | TG 45:9 | C48H74O6 | [M+K]+ |
| 722.596922 | 722.5929 | 0.004 | DGTS 33:2 | C43H79NO7 | [M+H]+ |
| 722.555096 | 722.5541 | 0.001 | DGCC 30:0 | C40H77NO8 | [M+Na]+ |
| 846.677519 | 846.6793 | 0.0018 | DGCC 39:1 | C49H93NO8 | [M+Na]+ |
| 715.510402 | 715.5062 | 0.0042 | DG O-42:9 | C45H72O4 | [M+K]+ |
| 770.494016 | 770.4943 | 0.0002 | PS 33:1 | C39H74NO10P | [M+Na]+ |
| 945.564504 | 945.5616 | 0.0029 | PG 48:12 | C54H83O10P | [M+Na]+ |
| 655.426677 | 655.4309 | 0.0042 | PA 31:1 | C34H65O8P | [M+Na]+ |
| 661.591144 | 661.5918 | 0.0007 | CE 19:3 | C46H76O2 | [M+H]+ |
| 745.543582 | 745.5402 | 0.0034 | TG 45:10 | C48H72O6 | [M+H]+ |
| 1009.58567 | 1009.5859 | 0.0002 | DGDG 40:9 | C55H86O15 | [M+Na]+ |
| 666.301129 | 666.3168 | 0.0156 | PS O-26:5 | C32H54NO9P | [M+K]+ |
| 775.56769 | 775.5637 | 0.004 | TG O-45:7 | C48H80O5 | [M+K]+ |
| 900.622914 | 900.6243 | 0.0014 | PE O-46:6 | C51H92NO7P | [M+K]+ |
| 796.509078 | 796.5099 | 0.0008 | PS 35:2 | C41H76NO10P | [M+Na]+ |
| 694.449826 | 694.4442 | 0.0056 | PE 33:6 | C38H64NO8P | [M+H]+ |
| 803.598944 | 803.595 | 0.0039 | TG O-47:7 | C50H84O5 | [M+K]+ |
| 734.605646 | 734.6058 | 0.0002 | PE O-36:0 | C41H84NO7P | [M+H]+ |
| 1271.60558 | 1271.5938 | 0.0118 | PIP2 52:9 | C61H103O19P3 | [M+K]+ |
| 977.619043 | 977.6196 | 0.0006 | DGDG 39:7 | C54H88O15 | [M+H]+ |
| 1181.59812 | 1181.6042 | 0.0061 | PIP2 46:4 | C55H101O19P3 | [M+Na]+ |
| 643.378315 | 643.3759 | 0.0024 | TG O-36:10 | C39H56O5 | [M+K]+ |
| 855.621178 | 855.6238 | 0.0026 | PA O-47:6 | C50H89O7P | [M+Na]+ |
| 644.531218 | 644.5096 | 0.0216 | DGCC 26:0 | C36H69NO8 | [M+H]+ |
| 524.262693 | 524.2538 | 0.0089 | LPE O-20:5 | C25H44NO6P | [M+K]+ |
| 576.304015 | 576.3061 | 0.002 | PE O-24:6 | C29H48NO7P | [M+Na]+ |
| 856.587887 | 856.5851 | 0.0028 | PE 45:9 | C50H82NO8P | [M+H]+ |
| 818.494685 | 818.4943 | 0.0004 | PS 37:5 | C43H74NO10P | [M+Na]+ |
| 795.533901 | 795.5324 | 0.0015 | TG O-47:11 | C50H76O5 | [M+K]+ |
| 616.563589 | 616.5333 | 0.0303 | NAT 34:0 | C36H73NO4S | [M+H]+ |
| 908.532749 | 908.5412 | 0.0085 | PS 44:9 | C50H80NO10P | [M+Na]+ |
| 1109.57371 | 1109.5492 | 0.0245 | PIP 46:8 | C55H92O16P2 | [M+K]+ |
| 681.357981 | 681.3609 | 0.0029 | MGDG 28:8 | C37H54O10 | [M+Na]+ |
| 810.503599 | 810.5044 | 0.0008 | PE 40:8 | C45H74NO8P | [M+Na]+ |
| 745.458352 | 745.4569 | 0.0015 | PA O-38:6 | C41H71O7P | [M+K]+ |
| 662.450596 | 662.452 | 0.0014 | LPE O-31:5 | C36H66NO6P | [M+Na]+ |

| | | | | | |
|---|---|---|---|---|---|
| 815.635385 | 815.6314 | 0.004 | DG O-49:8 | C52H88O4 | [M+K]+ |
| 1289.64865 | 1289.6408 | 0.0079 | PIP2 53:7 | C62H109O19P3 | [M+K]+ |
| 808.510325 | 808.5099 | 0.0004 | PS 36:3 | C42H76NO10P | [M+Na]+ |
| 684.502035 | 684.4963 | 0.0058 | PE O-33:4 | C38H70NO7P | [M+H]+ |
| 1159.58997 | 1159.5649 | 0.0251 | PIP 50:11 | C59H94O16P2 | [M+K]+ |
| 1269.5931 | 1269.5782 | 0.0149 | PIP2 52:10 | C61H101O19P3 | [M+K]+ |
| 594.352661 | 594.353 | 0.0003 | PE O-25:4 | C30H54NO7P | [M+Na]+ |
| 981.565329 | 981.5732 | 0.0079 | SQDG 45:9 | C54H86O12S | [M+Na]+ |
| 850.497048 | 850.4993 | 0.0023 | PS O-41:10 | C47H74NO9P | [M+Na]+ |
| 1244.58658 | 1244.6182 | 0.0316 | CoA 33:0 | C54H100N7O17P3S | [M+H]+ |
| 690.506476 | 690.5068 | 0.0004 | PE 32:1 | C37H72NO8P | [M+H]+ |
| 509.265526 | 509.2664 | 0.0009 | DG 26:7 | C29H42O5 | [M+K]+ |
| 1287.82938 | 1287.8259 | 0.0035 | DGDG 59:11 | C74H120O15 | [M+K]+ |
| 897.668305 | 897.6708 | 0.0025 | PA O-50:6 | C53H95O7P | [M+Na]+ |
| 1099.62947 | 1099.6246 | 0.0048 | PIP 48:8 | C57H96O16P2 | [M+H]+ |
| 903.51881 | 903.5147 | 0.0042 | PG 45:12 | C51H77O10P | [M+Na]+ |
| 1311.72122 | 1311.7214 | 0.0002 | PIP 61:12 | C70H114O16P2 | [M+K]+ |
| 813.52286 | 813.5252 | 0.0024 | LPI O-33:3 | C42H79O11P | [M+Na]+ |
| 815.539143 | 815.5409 | 0.0017 | LPI O-33:2 | C42H81O11P | [M+Na]+ |
| 788.434352 | 788.4263 | 0.008 | PS O-35:7 | C41H68NO9P | [M+K]+ |
| 1161.6009 | 1161.5805 | 0.0204 | PIP 50:10 | C59H96O16P2 | [M+K]+ |
| 801.543377 | 801.543 | 0.0004 | TG 46:8 | C49H78O6 | [M+K]+ |
| 1005.65362 | 1005.6555 | 0.0019 | PG 52:10 | C58H95O10P | [M+Na]+ |
| 844.561847 | 844.5617 | 0.0002 | PE O-42:6 | C47H84NO7P | [M+K]+ |
| 1268.58615 | 1268.5584 | 0.0277 | CoA 32:0 | C53H98N7O17P3S | [M+K]+ |
| 828.515623 | 828.5151 | 0.0005 | PS 36:1 | C42H80NO10P | [M+K]+ |
| 765.504212 | 765.5041 | 0.0001 | PG O-35:4 | C41H75O9P | [M+Na]+ |
| 782.495025 | 782.4943 | 0.0008 | PS 34:2 | C40H74NO10P | [M+Na]+ |
| 760.557717 | 760.5488 | 0.0089 | DGTS 33:2 | C43H79NO7 | [M+K]+ |
| 500.290668 | 500.2805 | 0.0102 | NAT 25:6 | C27H43NO4S | [M+Na]+ |
| 622.45719 | 622.4653 | 0.0081 | DGTS 24:0 | C34H65NO7 | [M+Na]+ |
| 756.47917 | 756.4786 | 0.0006 | PS 32:1 | C38H72NO10P | [M+Na]+ |
| 801.619519 | 801.6158 | 0.0038 | DG O-48:8 | C51H86O4 | [M+K]+ |
| 1091.56485 | 1091.562 | 0.0028 | PIP 48:12 | C57H88O16P2 | [M+H]+ |
| 843.658343 | 843.6627 | 0.0044 | DG O-51:8 | C54H92O4 | [M+K]+ |
| 827.591684 | 827.5925 | 0.0008 | PA O-45:6 | C48H85O7P | [M+Na]+ |
| 769.518501 | 769.5168 | 0.0017 | TG O-45:10 | C48H74O5 | [M+K]+ |
| 554.309506 | 554.3007 | 0.0088 | LPE O-22:4 | C27H50NO6P | [M+K]+ |
| 700.525404 | 700.5252 | 0.0002 | PE O-32:0 | C37H76NO7P | [M+Na]+ |
| 1053.57826 | 1053.5805 | 0.0023 | PIP 41:1 | C50H96O16P2 | [M+K]+ |
| 671.461652 | 671.4622 | 0.0006 | PA 32:0 | C35H69O8P | [M+Na]+ |
| 804.493521 | 804.4939 | 0.0003 | PE O-41:11 | C46H72NO7P | [M+Na]+ |
| 1035.56529 | 1035.5628 | 0.0025 | SQDG 48:11 | C57H88O12S | [M+K]+ |
| 810.542808 | 810.5432 | 0.0004 | PE O-43:11 | C48H76NO7P | [M+H]+ |
| 575.2552 | 575.2534 | 0.0018 | PA O-26:7 | C29H45O7P | [M+K]+ |

| | | | | | |
|---|---|---|---|---|---|
| 1114.64176 | 1114.6873 | 0.0455 | PS 58:12 | C64H102NO10P | [M+K]+ |
| 545.247331 | 545.2486 | 0.0013 | BMP 18:2 | C24H43O10P | [M+Na]+ |
| 663.479066 | 663.4749 | 0.0042 | DG O-38:7 | C41H68O4 | [M+K]+ |
| 784.489056 | 784.4889 | 0.0001 | PS O-34:2 | C40H76NO9P | [M+K]+ |
| 769.375108 | 769.3689 | 0.0062 | PI O-28:5 | C37H63O12P | [M+K]+ |
| 615.353105 | 615.3505 | 0.0026 | MGDG 21:0 | C30H56O10 | [M+K]+ |
| 807.496341 | 807.496 | 0.0003 | TG 47:12 | C50H72O6 | [M+K]+ |
| 927.650228 | 927.6473 | 0.0029 | TG 57:14 | C60H88O6 | [M+Na]+ |
| 502.48631 | 502.4618 | 0.0245 | NAE 31:4 | C33H59NO2 | [M+H]+ |
| 841.601837 | 841.6082 | 0.0063 | PA O-46:6 | C49H87O7P | [M+Na]+ |
| 692.433612 | 692.4286 | 0.005 | PE 33:7 | C38H62NO8P | [M+H]+ |
| 766.486619 | 766.4784 | 0.0083 | PE 35:3 | C40H74NO8P | [M+K]+ |
| 767.559684 | 767.5586 | 0.001 | TG 43:4 | C46H80O6 | [M+K]+ |
| 714.45471 | 714.4493 | 0.0054 | PE O-36:10 | C41H64NO7P | [M+H]+ |
| 511.280909 | 511.2819 | 0.001 | PA O-24:6 | C27H43O7P | [M+H]+ |
| 1067.59946 | 1067.5985 | 0.0009 | PI 49:10 | C58H93O13P | [M+K]+ |
| 768.443572 | 768.4365 | 0.0071 | PE O-37:9 | C42H68NO7P | [M+K]+ |
| 787.50697 | 787.5096 | 0.0026 | LPI O-31:2 | C40H77O11P | [M+Na]+ |
| 533.266482 | 533.264 | 0.0025 | PA 21:0 | C24H47O8P | [M+K]+ |
| 819.658875 | 819.6627 | 0.0038 | DG O-49:6 | C52H92O4 | [M+K]+ |
| 759.46345 | 759.4595 | 0.0039 | PG O-37:10 | C43H67O9P | [M+H]+ |
| 689.469651 | 689.4728 | 0.0031 | PG O-29:0 | C35H71O9P | [M+Na]+ |
| 510.230372 | 510.2381 | 0.0078 | LPE O-19:5 | C24H42NO6P | [M+K]+ |
| 761.490004 | 761.4882 | 0.0018 | PA O-39:5 | C42H75O7P | [M+K]+ |
| 810.490817 | 810.4834 | 0.0074 | PE O-40:9 | C45H74NO7P | [M+K]+ |
| 693.527943 | 693.5219 | 0.0061 | DG O-40:6 | C43H74O4 | [M+K]+ |
| 859.529583 | 859.5307 | 0.0011 | PI 34:1 | C43H81O13P | [M+Na]+ |
| 858.612404 | 858.6218 | 0.0094 | DGTS 42:8 | C52H85NO7 | [M+Na]+ |
| 1093.55057 | 1093.5753 | 0.0247 | PIP 46:8 | C55H92O16P2 | [M+Na]+ |
| 946.576532 | 946.5723 | 0.0043 | PE 49:11 | C54H86NO8P | [M+K]+ |
| 783.513314 | 783.5147 | 0.0013 | PG 35:2 | C41H77O10P | [M+Na]+ |
| 629.362028 | 629.3603 | 0.0017 | TG O-35:10 | C38H54O5 | [M+K]+ |
| 655.467525 | 655.4673 | 0.0002 | PA O-32:1 | C35H69O7P | [M+Na]+ |
| 763.467197 | 763.4673 | 0.0001 | PA O-41:10 | C44H69O7P | [M+Na]+ |
| 978.565871 | 978.5621 | 0.0038 | PS 48:10 | C54H86NO10P | [M+K]+ |
| 1125.70803 | 1125.7037 | 0.0044 | SQDG 54:8 | C63H106O12S | [M+K]+ |
| 770.459767 | 770.4521 | 0.0076 | PE O-37:8 | C42H70NO7P | [M+K]+ |
| 783.533755 | 783.5324 | 0.0013 | TG O-46:10 | C49H76O5 | [M+K]+ |
| 796.475353 | 796.4678 | 0.0076 | PE O-39:9 | C44H72NO7P | [M+K]+ |
| 896.656725 | 896.6528 | 0.004 | PE O-49:10 | C54H90NO7P | [M+H]+ |
| 686.270344 | 686.3065 | 0.0361 | PS 28:8 | C34H50NO10P | [M+Na]+ |
| 679.51192 | 679.5062 | 0.0057 | DG O-39:6 | C42H72O4 | [M+K]+ |
| 677.494191 | 677.4906 | 0.0036 | DG O-39:7 | C42H70O4 | [M+K]+ |
| 738.454818 | 738.4471 | 0.0078 | PE 33:3 | C38H70NO8P | [M+K]+ |
| 812.578738 | 812.5776 | 0.0012 | PS O-37:1 | C43H84NO9P | [M+Na]+ |

| | | | | | |
|---|---|---|---|---|---|
| 571.471321 | 571.4721 | 0.0008 | MG 34:6 | C37H62O4 | [M+H]+ |
| 1243.57724 | 1243.5625 | 0.0147 | PIP2 50:9 | C59H99O19P3 | [M+K]+ |
| 947.522766 | 947.5199 | 0.0029 | PG 47:12 | C53H81O10P | [M+K]+ |
| 723.557559 | 723.5558 | 0.0017 | TG 43:7 | C46H74O6 | [M+H]+ |
| 1009.5022 | 1009.5179 | 0.0157 | PIP 38:2 | C47H88O16P2 | [M+K]+ |
| 1225.58619 | 1225.573 | 0.0132 | PIP3 45:4 | C54H100O22P4 | [M+H]+ |
| 978.544629 | 978.5621 | 0.0175 | PS 48:10 | C54H86NO10P | [M+K]+ |
| 562.230006 | 562.254 | 0.024 | PE 22:6 | C27H42NO8P | [M+Na]+ |
| 881.514061 | 881.515 | 0.001 | PI 36:4 | C45H79O13P | [M+Na]+ |
| 1250.56496 | 1250.5688 | 0.0039 | CoA 32:1 | C53H96N7O17P3S | [M+Na]+ |
| 705.447215 | 705.4467 | 0.0005 | PG O-29:0 | C35H71O9P | [M+K]+ |
| 905.533429 | 905.5303 | 0.0031 | PG 45:11 | C51H79O10P | [M+Na]+ |
| 755.501247 | 755.5011 | 0.0001 | TG O-44:10 | C47H72O5 | [M+K]+ |
| 726.431687 | 726.4317 | 0 | PS 30:2 | C36H66NO10P | [M+Na]+ |
| 574.289662 | 574.2906 | 0.0009 | PE 21:1 | C26H50NO8P | [M+K]+ |
| 739.449874 | 739.4521 | 0.0022 | PG 32:3 | C38H69O10P | [M+Na]+ |
| 1075.55754 | 1075.5649 | 0.0073 | PIP 43:4 | C52H94O16P2 | [M+K]+ |
| 737.418056 | 737.4178 | 0.0003 | TG 42:12 | C45H62O6 | [M+K]+ |
| 735.434692 | 735.4348 | 0.0001 | SQDG 28:2 | C37H66O12S | [M+H]+ |
| 749.552563 | 749.5481 | 0.0045 | TG O-43:6 | C46H78O5 | [M+K]+ |
| 549.376783 | 549.3914 | 0.0146 | MG 31:7 | C34H54O4 | [M+Na]+ |
| 804.477828 | 804.4786 | 0.0008 | PS 36:5 | C42H72NO10P | [M+Na]+ |
| 857.512419 | 857.515 | 0.0026 | PI 34:2 | C43H79O13P | [M+Na]+ |
| 701.495981 | 701.4906 | 0.0054 | DG O-41:9 | C44H70O4 | [M+K]+ |
| 772.617936 | 772.6215 | 0.0035 | PE O-39:2 | C44H86NO7P | [M+H]+ |
| 747.536783 | 747.5324 | 0.0044 | TG O-43:7 | C46H76O5 | [M+K]+ |
| 659.279401 | 659.2803 | 0.0009 | PI 20:3 | C29H49O13P | [M+Na]+ |
| 750.592578 | 750.5878 | 0.0047 | DGCC 34:3 | C44H79NO8 | [M+H]+ |
| 925.517316 | 925.5201 | 0.0028 | PI O-41:10 | C50H79O12P | [M+Na]+ |
| 1131.56121 | 1131.5886 | 0.0274 | PIP2 42:1 | C51H99O19P3 | [M+Na]+ |
| 576.354632 | 576.3531 | 0.0015 | DGCC 22:6 | C32H49NO8 | [M+H]+ |
| 934.567277 | 934.5723 | 0.005 | PE 48:10 | C53H86NO8P | [M+K]+ |
| 871.644404 | 871.6423 | 0.0021 | PG 43:3 | C49H91O10P | [M+H]+ |
| 886.550387 | 886.5569 | 0.0065 | PS 42:6 | C48H82NO10P | [M+Na]+ |
| 501.258781 | 501.2588 | 0 | PA 20:1 | C23H43O8P | [M+Na]+ |
| 508.253457 | 508.2436 | 0.0098 | LPS O-15:0 | C21H44NO8P | [M+K]+ |
| 664.357577 | 664.3585 | 0.0009 | PS O-27:5 | C33H56NO9P | [M+Na]+ |
| 884.604911 | 884.6011 | 0.0038 | DGCC 43:10 | C53H83NO8 | [M+Na]+ |
| 687.57036 | 687.5688 | 0.0015 | DG O-39:2 | C42H80O4 | [M+K]+ |
| 1101.54566 | 1101.544 | 0.0016 | PIP2 42:5 | C51H91O19P3 | [M+H]+ |
| 1085.55141 | 1085.5492 | 0.0022 | PIP 44:6 | C53H92O16P2 | [M+K]+ |
| 1259.79771 | 1259.7946 | 0.0031 | DGDG 57:11 | C72H116O15 | [M+K]+ |
| 536.244505 | 536.2385 | 0.006 | LPS 16:0 | C22H44NO9P | [M+K]+ |
| 1069.58098 | 1069.5777 | 0.0033 | PIP 46:9 | C55H90O16P2 | [M+H]+ |
| 947.502184 | 947.5046 | 0.0025 | PI 40:7 | C49H81O13P | [M+K]+ |

| | | | | | |
|---|---|---|---|---|---|
| 972.589957 | 972.5879 | 0.002 | PE 51:12 | C56H88NO8P | [M+K]+ |
| 689.28944 | 689.2851 | 0.0043 | PG 28:8 | C34H51O10P | [M+K]+ |
| 708.545687 | 708.5409 | 0.0048 | DGCC 31:3 | C41H73NO8 | [M+H]+ |
| 735.458232 | 735.4571 | 0.0011 | PG O-33:5 | C39H69O9P | [M+Na]+ |
| 566.35794 | 566.3581 | 0.0002 | LPE O-24:4 | C29H54NO6P | [M+Na]+ |
| 1381.60619 | 1381.6071 | 0.0009 | PIP3 54:8 | C63H110O22P4 | [M+K]+ |
| 683.40916 | 683.4072 | 0.0019 | TG O-39:11 | C42H60O5 | [M+K]+ |
| 649.476411 | 649.4802 | 0.0038 | TG O-37:6 | C40H66O5 | [M+Na]+ |
| 524.29898 | 524.2983 | 0.0007 | LPS 18:1 | C24H46NO9P | [M+H]+ |
| 923.502748 | 923.5045 | 0.0017 | PI O-41:11 | C50H77O12P | [M+Na]+ |
| 620.221741 | 620.2595 | 0.0378 | PS 23:6 | C29H44NO10P | [M+Na]+ |
| 1245.59069 | 1245.5782 | 0.0125 | PIP2 50:8 | C59H101O19P3 | [M+K]+ |
| 897.487179 | 897.4888 | 0.0017 | PI O-39:10 | C48H75O12P | [M+Na]+ |
| 879.497583 | 879.4994 | 0.0018 | PI 36:5 | C45H77O13P | [M+Na]+ |
| 644.43997 | 644.4416 | 0.0016 | LPE O-28:1 | C33H68NO6P | [M+K]+ |
| 708.530682 | 708.5302 | 0.0004 | LPE O-34:3 | C39H76NO6P | [M+Na]+ |
| 854.490039 | 854.4943 | 0.0042 | PS 40:8 | C46H74NO10P | [M+Na]+ |
| 855.552936 | 855.5534 | 0.0005 | PG O-44:11 | C50H79O9P | [M+H]+ |
| 600.349872 | 600.3507 | 0.0008 | DGCC 22:5 | C32H51NO8 | [M+Na]+ |
| 850.665858 | 850.666 | 0.0001 | PE O-43:2 | C48H94NO7P | [M+Na]+ |
| 1105.54513 | 1105.5179 | 0.0272 | PIP 46:10 | C55H88O16P2 | [M+K]+ |
| 703.510647 | 703.5062 | 0.0044 | DG O-41:8 | C44H72O4 | [M+K]+ |
| 749.38484 | 749.3907 | 0.0058 | SQDG 26:0 | C35H66O12S | [M+K]+ |
| 577.350027 | 577.35 | 0 | PG O-23:3 | C29H53O9P | [M+H]+ |
| 502.254344 | 502.254 | 0.0003 | LPS O-16:2 | C22H42NO8P | [M+Na]+ |
| 681.263931 | 681.2646 | 0.0007 | PI 22:6 | C31H47O13P | [M+Na]+ |
| 501.315008 | 501.3315 | 0.0165 | LPA O-22:1 | C25H51O6P | [M+Na]+ |
| 769.352855 | 769.3594 | 0.0065 | SQDG 28:4 | C37H62O12S | [M+K]+ |
| 679.342156 | 679.3429 | 0.0007 | PI 21:0 | C30H57O13P | [M+Na]+ |
| 924.510073 | 924.5151 | 0.0051 | PS 44:9 | C50H80NO10P | [M+K]+ |
| 971.580752 | 971.5889 | 0.0081 | SQDG 44:7 | C53H88O12S | [M+Na]+ |
| 1033.50408 | 1033.5179 | 0.0138 | PIP 40:4 | C49H88O16P2 | [M+K]+ |
| 548.429139 | 548.4438 | 0.0147 | NAE 33:6 | C35H59NO2 | [M+Na]+ |
| 1059.56163 | 1059.5336 | 0.0281 | PIP 42:5 | C51H90O16P2 | [M+K]+ |
| 676.32208 | 676.3221 | 0 | PS 27:6 | C33H52NO10P | [M+Na]+ |
| 1135.55087 | 1135.5625 | 0.0116 | PIP2 41:0 | C50H99O19P3 | [M+K]+ |
| 1115.56461 | 1115.5597 | 0.0049 | PIP2 43:5 | C52H93O19P3 | [M+H]+ |
| 687.275049 | 687.2811 | 0.0061 | SQDG 22:3 | C31H52O12S | [M+K]+ |
| 996.73133 | 996.7263 | 0.005 | DGCC 51:10 | C61H99NO8 | [M+Na]+ |
| 755.482961 | 755.4834 | 0.0004 | PG 33:2 | C39H73O10P | [M+Na]+ |
| 902.51379 | 902.5097 | 0.0041 | PE 46:12 | C51H78NO8P | [M+K]+ |
| 1253.58308 | 1253.6018 | 0.0188 | PIP3 45:1 | C54H106O22P4 | [M+Na]+ |
| 515.36961 | 515.3707 | 0.0011 | DG 27:3 | C30H52O5 | [M+Na]+ |
| 693.428409 | 693.4256 | 0.0028 | PA O-34:4 | C37H67O7P | [M+K]+ |
| 729.451178 | 729.4491 | 0.0021 | TG 41:9 | C44H66O6 | [M+K]+ |

| | | | | | |
|---|---|---|---|---|---|
| 540.247735 | 540.2487 | 0.001 | PE O-20:4 | C25H44NO7P | [M+K]+ |
| 747.369644 | 747.375 | 0.0054 | SQDG 26:1 | C35H64O12S | [M+K]+ |
| 719.337359 | 719.3321 | 0.0053 | PG 30:7 | C36H57O10P | [M+K]+ |
| 734.473498 | 734.4731 | 0.0004 | PE 34:4 | C39H70NO8P | [M+Na]+ |
| 797.606 | 797.6056 | 0.0004 | TG 45:3 | C48H86O6 | [M+K]+ |
| 957.453702 | 957.4525 | 0.0012 | PIP 38:9 | C47H74O16P2 | [M+H]+ |
| 1145.60194 | 1145.6042 | 0.0023 | PIP2 43:1 | C52H101O19P3 | [M+Na]+ |
| 909.582387 | 909.5827 | 0.0003 | PI O-39:4 | C48H87O12P | [M+Na]+ |
| 807.478358 | 807.4784 | 0.0001 | PI O-30:0 | C39H77O12P | [M+K]+ |
| 1083.53672 | 1083.5336 | 0.0032 | PIP 44:7 | C53H90O16P2 | [M+K]+ |
| 847.453853 | 847.445 | 0.0088 | DGDG 28:6 | C43H68O15 | [M+Na]+ |
| 789.438893 | 789.4315 | 0.0074 | PI O-29:2 | C38H71O12P | [M+K]+ |
| 927.706895 | 927.7049 | 0.002 | PG 47:3 | C53H99O10P | [M+H]+ |
| 667.305042 | 667.3008 | 0.0043 | PG 26:5 | C32H53O10P | [M+K]+ |
| 872.477956 | 872.4838 | 0.0059 | PS 40:7 | C46H76NO10P | [M+K]+ |
| 817.642921 | 817.6471 | 0.0041 | DG O-49:7 | C52H90O4 | [M+K]+ |
| 787.529819 | 787.5273 | 0.0025 | TG 45:8 | C48H76O6 | [M+K]+ |
| 589.572973 | 589.553 | 0.02 | MG O-34:1 | C37H74O3 | [M+Na]+ |
| 836.529734 | 836.5225 | 0.0073 | PE 44:12 | C49H74NO8P | [M+H]+ |
| 531.365867 | 531.3785 | 0.0126 | LPA O-24:0 | C27H57O6P | [M+Na]+ |
| 767.49925 | 767.4988 | 0.0005 | PA 38:2 | C41H77O8P | [M+K]+ |
| 838.536534 | 838.5359 | 0.0007 | PS O-38:3 | C44H82NO9P | [M+K]+ |
| 611.427908 | 611.4282 | 0.0003 | TG 33:4 | C36H60O6 | [M+Na]+ |
| 679.375197 | 679.3736 | 0.0016 | PA 32:4 | C35H61O8P | [M+K]+ |
| 1309.81006 | 1309.7996 | 0.0104 | PIP 60:6 | C69H124O16P2 | [M+K]+ |
| 723.346712 | 723.348 | 0.0013 | PI O-26:6 | C35H57O12P | [M+Na]+ |
| 762.454284 | 762.4471 | 0.0072 | PE 35:5 | C40H70NO8P | [M+K]+ |
| 848.481903 | 848.4837 | 0.0018 | PS O-41:11 | C47H72NO9P | [M+Na]+ |
| 1093.61294 | 1093.6118 | 0.0011 | PIP 44:2 | C53H100O16P2 | [M+K]+ |
| 857.589661 | 857.5821 | 0.0076 | PA O-46:6 | C49H87O7P | [M+K]+ |
| 543.457914 | 543.4538 | 0.0041 | FA 34:2 | C34H64O2 | [M+K]+ |
| 851.501341 | 851.4988 | 0.0026 | PA 45:9 | C48H77O8P | [M+K]+ |
| 589.466101 | 589.4593 | 0.0068 | MG 32:2 | C35H66O4 | [M+K]+ |
| 677.327393 | 677.3272 | 0.0001 | PI 21:1 | C30H55O13P | [M+Na]+ |
| 1071.58678 | 1071.5909 | 0.0041 | PIP 44:5 | C53H94O16P2 | [M+Na]+ |
| 533.233206 | 533.2276 | 0.0056 | LPG 17:2 | C23H43O9P | [M+K]+ |
| 1100.63848 | 1100.6716 | 0.0332 | PS 57:12 | C63H100NO10P | [M+K]+ |
| 754.478839 | 754.4784 | 0.0005 | PE 34:2 | C39H74NO8P | [M+K]+ |
| 1103.56165 | 1103.5596 | 0.002 | PIP 47:10 | C56H90O16P2 | [M+Na]+ |
| 805.497091 | 805.499 | 0.0019 | PG 37:5 | C43H75O10P | [M+Na]+ |
| 649.387195 | 649.3865 | 0.0007 | TG 35:7 | C38H58O6 | [M+K]+ |
| 963.518443 | 963.5263 | 0.0078 | SQDG 44:11 | C53H80O12S | [M+Na]+ |
| 1379.59153 | 1379.5914 | 0.0001 | PIP3 54:9 | C63H108O22P4 | [M+K]+ |
| 596.294928 | 596.2959 | 0.001 | PS O-22:4 | C28H48NO9P | [M+Na]+ |
| 1251.56902 | 1251.5862 | 0.0172 | PIP3 45:2 | C54H104O22P4 | [M+Na]+ |

| | | | | | |
|---|---|---|---|---|---|
| 938.57173 | 938.5672 | 0.0046 | PS O-46:9 | C52H86NO9P | [M+K]+ |
| 581.26807 | 581.2697 | 0.0017 | LPI 15:0 | C24H47O12P | [M+Na]+ |
| 955.438626 | 955.4368 | 0.0018 | PIP 38:10 | C47H72O16P2 | [M+H]+ |
| 713.545748 | 713.5456 | 0.0002 | PA O-36:0 | C39H79O7P | [M+Na]+ |
| 1058.59518 | 1058.6247 | 0.0295 | PS 54:12 | C60H94NO10P | [M+K]+ |
| 811.506128 | 811.5038 | 0.0023 | PA O-43:8 | C46H77O7P | [M+K]+ |
| 549.230316 | 549.2378 | 0.0075 | PA O-24:6 | C27H43O7P | [M+K]+ |
| 803.48306 | 803.4834 | 0.0003 | PG 37:6 | C43H73O10P | [M+Na]+ |
| 741.450257 | 741.4491 | 0.0012 | TG 42:10 | C45H66O6 | [M+K]+ |
| 827.691175 | 827.6912 | 0 | DG O-53:11 | C56H90O4 | [M+H]+ |
| 830.418867 | 830.4369 | 0.018 | PS 37:7 | C43H70NO10P | [M+K]+ |
| 1211.60591 | 1211.5938 | 0.0121 | PIP2 47:4 | C56H103O19P3 | [M+K]+ |
| 690.29953 | 690.3168 | 0.0172 | PS O-28:7 | C34H54NO9P | [M+K]+ |
| 761.471217 | 761.4728 | 0.0016 | PG O-35:6 | C41H71O9P | [M+Na]+ |
| 512.246338 | 512.2538 | 0.0074 | LPE O-19:4 | C24H44NO6P | [M+K]+ |
| 1283.56947 | 1283.5573 | 0.0122 | PIP3 50:10 | C59H98O22P4 | [M+H]+ |
| 676.285221 | 676.3011 | 0.0159 | PS O-27:7 | C33H52NO9P | [M+K]+ |
| 802.388234 | 802.4056 | 0.0174 | PS 35:7 | C41H66NO10P | [M+K]+ |
| 727.26985 | 727.2855 | 0.0157 | PI 24:5 | C33H53O13P | [M+K]+ |
| 657.336665 | 657.3374 | 0.0008 | LPI O-22:4 | C31H55O11P | [M+Na]+ |
| 1089.54806 | 1089.544 | 0.004 | PIP2 41:4 | C50H91O19P3 | [M+H]+ |
| 551.237485 | 551.2382 | 0.0007 | BMP 17:0 | C23H45O10P | [M+K]+ |
| 1267.57786 | 1267.5625 | 0.0154 | PIP2 52:11 | C61H99O19P3 | [M+K]+ |
| 1309.58522 | 1309.573 | 0.0123 | PIP3 52:11 | C61H100O22P4 | [M+H]+ |
| 537.295236 | 537.2953 | 0.0001 | LPG O-18:0 | C24H51O8P | [M+K]+ |
| 1168.52932 | 1168.493 | 0.0363 | CoA 28:3 | C49H84N7O17P3S | [M+H]+ |
| 960.572604 | 960.5725 | 0.0001 | PS 48:11 | C54H84NO10P | [M+Na]+ |
| 1079.52564 | 1079.5023 | 0.0234 | PIP 44:9 | C53H86O16P2 | [M+K]+ |
| 673.368017 | 673.3687 | 0.0007 | LPI O-23:3 | C32H59O11P | [M+Na]+ |
| 1031.48601 | 1031.5023 | 0.0163 | PIP 40:5 | C49H86O16P2 | [M+K]+ |
| 1202.599 | 1202.5713 | 0.0277 | CoA 30:0 | C51H94N7O17P3S | [M+H]+ |
| 652.286017 | 652.3011 | 0.0151 | PS O-25:5 | C31H52NO9P | [M+K]+ |
| 1389.59872 | 1389.5758 | 0.0229 | PIP3 55:11 | C64H106O22P4 | [M+K]+ |
| 1485.62142 | 1485.6697 | 0.0483 | PIP3 62:12 | C71H118O22P4 | [M+K]+ |
| 571.261978 | 571.2643 | 0.0023 | PG 20:3 | C26H45O10P | [M+Na]+ |
| 849.486619 | 849.4888 | 0.0022 | PI O-35:6 | C44H75O12P | [M+Na]+ |
| 727.24861 | 727.249 | 0.0004 | PIP 21:5 | C30H48O16P2 | [M+H]+ |
| 757.481933 | 757.4804 | 0.0015 | TG 43:9 | C46H70O6 | [M+K]+ |

*Supporting Table 3: List of annotations for signals of the OX1 cell line identified as statistically different (ANOVA multiple-sample test, p-value ≤ 0.05) between the naïve and LPS-stimulated state (biological triplicates, each consisting of 22,500 individual mass spectra of multiple individual cells) in positive-ion mode from LIPID MAPS database research with the lowest mass difference for each input mass.*

| Input mass | Matched mass | Δ $m/z$ | Name | Formula | Ion |
|---|---|---|---|---|---|
| 593.3447 | 593.3449 | 0.0002 | PG 23:2 | C29H53O10P | [M+H]+ |

| | | | | | |
|---|---|---|---|---|---|
| 897.58075 | 897.5827 | 0.002 | PI O-38:3 | C47H87O12P | [M+Na]+ |
| 661.464157 | 661.4593 | 0.0049 | DG O-38:8 | C41H66O4 | [M+K]+ |
| 809.630324 | 809.6395 | 0.0091 | PA O-43:1 | C46H91O7P | [M+Na]+ |
| 518.28621 | 518.2853 | 0.0009 | LPS O-17:1 | C23H46NO8P | [M+Na]+ |
| 717.562395 | 717.5511 | 0.0113 | MGDG 31:0 | C40H76O10 | [M+H]+ |
| 587.552039 | 587.5374 | 0.0147 | MG O-34:2 | C37H72O3 | [M+Na]+ |
| 689.495009 | 689.4906 | 0.0044 | DG O-40:8 | C43H70O4 | [M+K]+ |
| 755.588684 | 755.5925 | 0.0038 | PA O-39:0 | C42H85O7P | [M+Na]+ |
| 731.578618 | 731.5668 | 0.0118 | MGDG 32:0 | C41H78O10 | [M+H]+ |
| 728.411316 | 728.4052 | 0.0061 | PE O-34:8 | C39H64NO7P | [M+K]+ |
| 628.520996 | 628.5147 | 0.0063 | DGTS 26:0 | C36H69NO7 | [M+H]+ |
| 757.592101 | 757.5824 | 0.0097 | MGDG 34:1 | C43H80O10 | [M+H]+ |
| 770.423285 | 770.4158 | 0.0075 | PE 36:8 | C41H66NO8P | [M+K]+ |
| 717.575714 | 717.5792 | 0.0035 | DG O-43:7 | C46H78O4 | [M+Na]+ |
| 772.617936 | 772.6215 | 0.0035 | PE O-39:2 | C44H86NO7P | [M+H]+ |
| 634.394709 | 634.3867 | 0.008 | PE O-30:8 | C35H56NO7P | [M+H]+ |
| 1085.68702 | 1085.6819 | 0.0052 | PI O-51:8 | C60H103O12P | [M+K]+ |
| 577.350027 | 577.35 | 0 | PG O-23:3 | C29H53O9P | [M+H]+ |
| 675.517012 | 675.5042 | 0.0128 | MGDG 28:0 | C37H70O10 | [M+H]+ |
| 605.268576 | 605.2697 | 0.0012 | LPI 17:2 | C26H47O12P | [M+Na]+ |
| 503.20296 | 503.2017 | 0.0013 | BMP 15:2 | C21H37O10P | [M+Na]+ |
| 827.691175 | 827.6912 | 0 | DG O-53:11 | C56H90O4 | [M+H]+ |
| 767.395001 | 767.3896 | 0.0054 | LPI O-29:6 | C38H65O11P | [M+K]+ |
| 587.280258 | 587.2746 | 0.0057 | PG O-21:3 | C27H49O9P | [M+K]+ |

*Supporting Table 4: List of annotations for signals of the D9 cell line identified as statistically different (ANOVA multiple-sample test, p-value ≤ 0.01) between the naïve and LPS-stimulated state (biological triplicates, each consisting of 22,500 individual mass spectra of multiple individual cells) in positive-ion mode from LIPID MAPS database research with the lowest mass difference for each input mass.*

| Input mass | Matched mass | Δ $m/z$ | Name | Formula | Ion |
|---|---|---|---|---|---|
| 742.535415 | 742.5357 | 0.0003 | PE 34:0 | C39H78NO8P | [M+Na]+ |
| 741.530655 | 741.5276 | 0.003 | LPI O-29:0 | C38H77O11P | [M+H]+ |
| 728.520166 | 728.5201 | 0.0001 | PE 33:0 | C38H76NO8P | [M+Na]+ |
| 772.525534 | 772.5253 | 0.0002 | PE 35:0 | C40H80NO8P | [M+K]+ |
| 770.56746 | 770.567 | 0.0004 | PE 36:0 | C41H82NO8P | [M+Na]+ |
| 707.54418 | 707.5375 | 0.0067 | DG O-41:6 | C44H76O4 | [M+K]+ |
| 729.523732 | 729.5219 | 0.0019 | DG O-43:9 | C46H74O4 | [M+K]+ |
| 743.545473 | 743.5375 | 0.008 | DG O-44:9 | C47H76O4 | [M+K]+ |
| 787.458708 | 787.4545 | 0.0043 | PG 38:10 | C44H67O10P | [M+H]+ |
| 714.504775 | 714.5044 | 0.0004 | PE 32:0 | C37H74NO8P | [M+Na]+ |
| 685.435813 | 685.4415 | 0.0057 | PG O-29:2 | C35H67O9P | [M+Na]+ |
| 649.427011 | 649.4229 | 0.0041 | TG O-36:7 | C39H62O5 | [M+K]+ |
| 883.650475 | 883.6551 | 0.0046 | PA O-49:6 | C52H93O7P | [M+Na]+ |
| 828.536293 | 828.5385 | 0.0022 | DGCC 39:10 | C49H75NO8 | [M+Na]+ |
| 700.489251 | 700.4888 | 0.0005 | PE 31:0 | C36H72NO8P | [M+Na]+ |

| | | | | | |
|---|---|---|---|---|---|
| 713.453091 | 713.4541 | 0.001 | PA O-39:10 | C42H65O7P | [M+H]+ |
| 703.558242 | 703.5636 | 0.0053 | DG O-42:7 | C45H76O4 | [M+Na]+ |
| 922.593394 | 922.5934 | 0 | PS 43:3 | C49H90NO10P | [M+K]+ |
| 788.541911 | 788.5436 | 0.0017 | DGTS 37:8 | C47H75NO7 | [M+Na]+ |
| 663.442264 | 663.4385 | 0.0037 | TG O-37:7 | C40H64O5 | [M+K]+ |
| 752.502883 | 752.4991 | 0.0038 | PE O-35:3 | C40H76NO7P | [M+K]+ |
| 746.551194 | 746.5541 | 0.0029 | DGCC 32:2 | C42H77NO8 | [M+Na]+ |
| 658.388185 | 658.3845 | 0.0037 | PE 27:1 | C32H62NO8P | [M+K]+ |
| 790.515954 | 790.5147 | 0.0012 | PE O-38:5 | C43H78NO7P | [M+K]+ |
| 811.581108 | 811.5823 | 0.0012 | PG O-38:2 | C44H85O9P | [M+Na]+ |
| 794.586889 | 794.5905 | 0.0036 | DGTS 37:5 | C47H81NO7 | [M+Na]+ |
| 786.525867 | 786.5256 | 0.0003 | PS 34:0 | C40H78NO10P | [M+Na]+ |
| 789.601683 | 789.6028 | 0.0011 | TG 48:9 | C51H80O6 | [M+H]+ |
| 856.549104 | 856.5488 | 0.0003 | DGTS 41:10 | C51H79NO7 | [M+K]+ |
| 917.537296 | 917.5419 | 0.0046 | SQDG 40:6 | C49H82O12S | [M+Na]+ |
| 748.548421 | 748.5487 | 0.0003 | PS O-34:1 | C40H78NO9P | [M+H]+ |
| 1053.55399 | 1053.5464 | 0.0076 | PIP 45:10 | C54H86O16P2 | [M+H]+ |
| 633.358186 | 633.3609 | 0.0027 | MGDG 24:4 | C33H54O10 | [M+Na]+ |
| 686.421992 | 686.4158 | 0.0062 | PE 29:1 | C34H66NO8P | [M+K]+ |
| 549.338281 | 549.3341 | 0.0042 | DG O-30:8 | C33H50O4 | [M+K]+ |
| 1055.56656 | 1055.562 | 0.0045 | PIP 45:9 | C54H88O16P2 | [M+H]+ |
| 884.65462 | 884.6528 | 0.0019 | PE O-48:9 | C53H90NO7P | [M+H]+ |
| 918.562171 | 918.5621 | 0.0001 | PS 43:5 | C49H86NO10P | [M+K]+ |
| 904.529469 | 904.5253 | 0.0042 | PE 46:11 | C51H80NO8P | [M+K]+ |
| 758.423668 | 758.4158 | 0.0079 | PE 35:7 | C40H66NO8P | [M+K]+ |
| 816.572779 | 816.5749 | 0.0021 | DGTS 39:8 | C49H79NO7 | [M+Na]+ |
| 829.539966 | 829.5378 | 0.0022 | TG 50:14 | C53H74O6 | [M+Na]+ |
| 715.510402 | 715.5062 | 0.0042 | DG O-42:9 | C45H72O4 | [M+K]+ |
| 770.494016 | 770.4943 | 0.0002 | PS 33:1 | C39H74NO10P | [M+Na]+ |
| 684.404928 | 684.4001 | 0.0048 | PE 29:2 | C34H64NO8P | [M+K]+ |
| 796.509078 | 796.5099 | 0.0008 | PS 35:2 | C41H76NO10P | [M+Na]+ |
| 803.598944 | 803.595 | 0.0039 | TG O-47:7 | C50H84O5 | [M+K]+ |
| 643.378315 | 643.3759 | 0.0024 | TG O-36:10 | C39H56O5 | [M+K]+ |
| 713.497867 | 713.4963 | 0.0015 | LPI O-27:0 | C36H73O11P | [M+H]+ |
| 1230.21577 | 1230.1712 | 0.0445 | TG O-80:6 | C83H152O5 | [M+H]+ |
| 884.564454 | 884.5566 | 0.0078 | PE 44:7 | C49H84NO8P | [M+K]+ |
| 897.668305 | 897.6708 | 0.0025 | PA O-50:6 | C53H95O7P | [M+Na]+ |
| 855.589797 | 855.5898 | 0 | TG O-53:15 | C56H80O5 | [M+Na]+ |
| 1430.2264 | 1430.2314 | 0.005 | TG O-94:15 | C97H162O5 | [M+Na]+ |
| 844.561847 | 844.5617 | 0.0002 | PE O-42:6 | C47H84NO7P | [M+K]+ |
| 931.573644 | 931.5755 | 0.0018 | DGDG 32:0 | C47H88O15 | [M+K]+ |
| 782.495025 | 782.4943 | 0.0008 | PS 34:2 | C40H74NO10P | [M+Na]+ |
| 842.517474 | 842.5097 | 0.0078 | PE 41:7 | C46H78NO8P | [M+K]+ |
| 756.47917 | 756.4786 | 0.0006 | PS 32:1 | C38H72NO10P | [M+Na]+ |
| 810.542808 | 810.5432 | 0.0004 | PE O-43:11 | C48H76NO7P | [M+H]+ |

| | | | | | |
|---|---|---|---|---|---|
| 818.590234 | 818.5905 | 0.0003 | DGTS 39:7 | C49H81NO7 | [M+Na]+ |
| 581.365126 | 581.3603 | 0.0048 | TG O-31:6 | C34H54O5 | [M+K]+ |
| 635.447406 | 635.4436 | 0.0038 | DG O-36:7 | C39H64O4 | [M+K]+ |
| 880.550452 | 880.5488 | 0.0016 | DGTS 43:12 | C53H79NO7 | [M+K]+ |
| 800.505862 | 800.5072 | 0.0013 | DGCC 37:10 | C47H71NO8 | [M+Na]+ |
| 759.46345 | 759.4595 | 0.0039 | PG O-37:10 | C43H67O9P | [M+H]+ |
| 603.326367 | 603.3269 | 0.0005 | PG 22:1 | C28H53O10P | [M+Na]+ |
| 690.373948 | 690.3741 | 0.0002 | PS O-29:6 | C35H58NO9P | [M+Na]+ |
| 774.527324 | 774.5279 | 0.0006 | DGTS 36:8 | C46H73NO7 | [M+Na]+ |
| 629.362028 | 629.3603 | 0.0017 | TG O-35:10 | C38H54O5 | [M+K]+ |
| 964.553109 | 964.5464 | 0.0067 | PS 47:10 | C53H84NO10P | [M+K]+ |
| 679.51192 | 679.5062 | 0.0057 | DG O-39:6 | C42H72O4 | [M+K]+ |
| 1093.54596 | 1093.5179 | 0.028 | PIP 45:9 | C54H88O16P2 | [M+K]+ |
| 698.478021 | 698.4755 | 0.0025 | PE 33:4 | C38H68NO8P | [M+H]+ |
| 905.533429 | 905.5303 | 0.0031 | PG 45:11 | C51H79O10P | [M+Na]+ |
| 795.426463 | 795.4209 | 0.0056 | LPI O-31:6 | C40H69O11P | [M+K]+ |
| 823.457333 | 823.4522 | 0.0051 | LPI O-33:6 | C42H73O11P | [M+K]+ |
| 658.425311 | 658.4289 | 0.0036 | DGCC 26:4 | C36H61NO8 | [M+Na]+ |
| 749.552563 | 749.5481 | 0.0045 | TG O-43:6 | C46H78O5 | [M+K]+ |
| 676.35837 | 676.3585 | 0.0001 | PS O-28:6 | C34H56NO9P | [M+Na]+ |
| 662.342933 | 662.343 | 0.0001 | PS 24:0 | C30H58NO10P | [M+K]+ |
| 1000.5659 | 1000.6038 | 0.0379 | PS 51:12 | C57H88NO10P | [M+Na]+ |
| 631.342988 | 631.3453 | 0.0023 | MGDG 24:5 | C33H52O10 | [M+Na]+ |
| 925.517316 | 925.5201 | 0.0028 | PI O-41:10 | C50H79O12P | [M+Na]+ |
| 1007.567 | 1007.5702 | 0.0032 | DGDG 40:10 | C55H84O15 | [M+Na]+ |
| 1386.22278 | 1386.2264 | 0.0036 | TG 89:9 | C92H162O6 | [M+Na]+ |
| 1085.55141 | 1085.5492 | 0.0022 | PIP 44:6 | C53H92O16P2 | [M+K]+ |
| 1236.21401 | 1236.2158 | 0.0018 | TG O-78:0 | C81H160O5 | [M+Na]+ |
| 828.535228 | 828.5385 | 0.0033 | DGCC 39:10 | C49H75NO8 | [M+Na]+ |
| 760.510459 | 760.5123 | 0.0018 | DGTS 35:8 | C45H71NO7 | [M+Na]+ |
| 907.551129 | 907.546 | 0.0052 | PG 45:10 | C51H81O10P | [M+Na]+ |
| 1032.57714 | 1032.609 | 0.0319 | PS 52:11 | C58H92NO10P | [M+K]+ |
| 687.368191 | 687.3634 | 0.0048 | PG 27:2 | C33H61O10P | [M+K]+ |
| 568.241917 | 568.2436 | 0.0017 | PE 21:4 | C26H44NO8P | [M+K]+ |
| 852.501128 | 852.494 | 0.0071 | PE 42:9 | C47H76NO8P | [M+K]+ |
| 1095.56137 | 1095.5336 | 0.0278 | PIP 45:8 | C54H90O16P2 | [M+K]+ |
| 759.427249 | 759.4232 | 0.0041 | PG 36:10 | C42H63O10P | [M+H]+ |
| 867.592243 | 867.5899 | 0.0023 | TG 51:10 | C54H84O6 | [M+K]+ |
| 615.347016 | 615.3504 | 0.0034 | LPI 19:0 | C28H55O12P | [M+H]+ |
| 753.449868 | 753.4491 | 0.0008 | TG 43:11 | C46H66O6 | [M+K]+ |
| 902.51379 | 902.5097 | 0.0041 | PE 46:12 | C51H78NO8P | [M+K]+ |
| 794.49494 | 794.4943 | 0.0007 | PS 35:3 | C41H74NO10P | [M+Na]+ |
| 729.451178 | 729.4491 | 0.0021 | TG 41:9 | C44H66O6 | [M+K]+ |
| 747.369644 | 747.375 | 0.0054 | SQDG 26:1 | C35H64O12S | [M+K]+ |
| 962.537757 | 962.5308 | 0.007 | PS 47:11 | C53H82NO10P | [M+K]+ |

| | | | | | |
|---|---|---|---|---|---|
| 718.405347 | 718.4054 | 0.0001 | PS O-31:6 | C37H62NO9P | [M+Na]+ |
| 886.578989 | 886.5723 | 0.0067 | PE 44:6 | C49H86NO8P | [M+K]+ |
| 847.453853 | 847.445 | 0.0088 | DGDG 28:6 | C43H68O15 | [M+Na]+ |
| 1418.22952 | 1418.2314 | 0.0019 | TG O-93:14 | C96H162O5 | [M+Na]+ |
| 966.569076 | 966.5621 | 0.007 | PS 47:9 | C53H86NO10P | [M+K]+ |
| 978.531814 | 978.5621 | 0.0303 | PS 48:10 | C54H86NO10P | [M+K]+ |
| 787.529819 | 787.5273 | 0.0025 | TG 45:8 | C48H76O6 | [M+K]+ |
| 890.514832 | 890.5097 | 0.0052 | PE 45:11 | C50H78NO8P | [M+K]+ |
| 960.485562 | 960.5151 | 0.0296 | PS 47:12 | C53H80NO10P | [M+K]+ |
| 531.365867 | 531.3785 | 0.0126 | LPA O-24:0 | C27H57O6P | [M+Na]+ |
| 762.528361 | 762.5281 | 0.0003 | DGCC 32:2 | C42H77NO8 | [M+K]+ |
| 827.525279 | 827.5221 | 0.0031 | PG O-42:11 | C48H75O9P | [M+H]+ |
| 1055.57058 | 1055.562 | 0.0085 | PIP 45:9 | C54H88O16P2 | [M+H]+ |
| 952.551578 | 952.5464 | 0.0051 | PS 46:9 | C52H84NO10P | [M+K]+ |
| 886.608294 | 886.6086 | 0.0004 | PE O-45:6 | C50H90NO7P | [M+K]+ |
| 822.524033 | 822.5256 | 0.0015 | PS 37:3 | C43H78NO10P | [M+Na]+ |
| 922.541232 | 922.5359 | 0.0054 | PS O-45:10 | C51H82NO9P | [M+K]+ |
| 1073.57091 | 1073.5492 | 0.0217 | PIP 43:5 | C52H92O16P2 | [M+K]+ |
| 745.447205 | 745.4439 | 0.0033 | PG O-36:10 | C42H65O9P | [M+H]+ |
| 743.508611 | 743.5011 | 0.0075 | TG O-43:9 | C46H72O5 | [M+K]+ |
| 819.389665 | 819.3845 | 0.0051 | PI O-32:8 | C41H65O12P | [M+K]+ |
| 1191.58165 | 1191.5886 | 0.0069 | PIP2 47:6 | C56H99O19P3 | [M+Na]+ |
| 669.318277 | 669.3164 | 0.0018 | PG 26:4 | C32H55O10P | [M+K]+ |
| 871.55272 | 871.5484 | 0.0044 | PG 44:10 | C50H79O10P | [M+H]+ |
| 667.342195 | 667.3453 | 0.0031 | PI 22:2 | C31H55O13P | [M+H]+ |
| 767.395001 | 767.3896 | 0.0054 | LPI O-29:6 | C38H65O11P | [M+K]+ |
| 845.437568 | 845.4294 | 0.0082 | DGDG 28:7 | C43H66O15 | [M+Na]+ |
| 1087.70813 | 1087.7128 | 0.0046 | PG O-58:12 | C64H105O9P | [M+K]+ |
| 882.510303 | 882.5046 | 0.0057 | PS O-42:9 | C48H78NO9P | [M+K]+ |
| 1185.59395 | 1185.5805 | 0.0134 | PIP 52:12 | C61H96O16P2 | [M+K]+ |
| 625.331477 | 625.3347 | 0.0033 | PI O-20:2 | C29H53O12P | [M+H]+ |

*Supporting Table 5: List of annotations for signals of the C12 cell line identified as statistically different (ANOVA multiple-sample test, p-value ≤ 0.05) between the naïve and LPS-stimulated state (biological triplicates, each consisting of 22,500 individual mass spectra of multiple individual cells) in positive-ion mode from LIPID MAPS database research with the lowest mass difference for each input mass.*

| Input mass | Matched mass | Δ *m/z* | Name | Formula | Ion |
|---|---|---|---|---|---|
| 757.555242 | 757.5532 | 0.0021 | DG O-45:9 | C48H78O4 | [M+K]+ |
| 742.535415 | 742.5357 | 0.0003 | PE 34:0 | C39H78NO8P | [M+Na]+ |
| 756.551877 | 756.5514 | 0.0005 | PE 35:0 | C40H80NO8P | [M+Na]+ |
| 728.520166 | 728.5201 | 0.0001 | PE 33:0 | C38H76NO8P | [M+Na]+ |
| 772.525534 | 772.5253 | 0.0002 | PE 35:0 | C40H80NO8P | [M+K]+ |
| 901.564602 | 901.5718 | 0.0072 | PA 50:11 | C53H83O8P | [M+Na]+ |
| 770.56746 | 770.567 | 0.0004 | PE 36:0 | C41H82NO8P | [M+Na]+ |
| 932.559707 | 932.5566 | 0.0031 | PE 48:11 | C53H84NO8P | [M+K]+ |

| | | | | | |
|---|---|---|---|---|---|
| 842.55402 | 842.5541 | 0.0001 | DGCC 40:10 | C50H77NO8 | [M+Na]+ |
| 729.523732 | 729.5219 | 0.0019 | DG O-43:9 | C46H74O4 | [M+K]+ |
| 743.545473 | 743.5375 | 0.008 | DG O-44:9 | C47H76O4 | [M+K]+ |
| 884.600719 | 884.6011 | 0.0004 | DGCC 43:10 | C53H83NO8 | [M+Na]+ |
| 726.50467 | 726.5044 | 0.0002 | PE 33:1 | C38H74NO8P | [M+Na]+ |
| 753.523776 | 753.5219 | 0.0019 | DG O-45:11 | C48H74O4 | [M+K]+ |
| 704.522522 | 704.5225 | 0 | PE 33:1 | C38H74NO8P | [M+H]+ |
| 773.528812 | 773.5303 | 0.0015 | PG 34:0 | C40H79O10P | [M+Na]+ |
| 843.557386 | 843.5534 | 0.0039 | PG O-43:10 | C49H79O9P | [M+H]+ |
| 903.579749 | 903.5723 | 0.0074 | PI O-37:1 | C46H89O12P | [M+K]+ |
| 933.563955 | 933.5616 | 0.0024 | PG 47:11 | C53H83O10P | [M+Na]+ |
| 714.504775 | 714.5044 | 0.0004 | PE 32:0 | C37H74NO8P | [M+Na]+ |
| 770.53063 | 770.5306 | 0 | PS O-34:1 | C40H78NO9P | [M+Na]+ |
| 814.557138 | 814.5569 | 0.0003 | PS 36:0 | C42H82NO10P | [M+Na]+ |
| 794.530737 | 794.5306 | 0.0001 | PS O-36:3 | C42H78NO9P | [M+Na]+ |
| 700.489251 | 700.4888 | 0.0005 | PE 31:0 | C36H72NO8P | [M+Na]+ |
| 783.635194 | 783.6416 | 0.0064 | CE 25:3 | C52H88O2 | [M+K]+ |
| 697.462327 | 697.465 | 0.0027 | LPI O-26:1 | C35H69O11P | [M+H]+ |
| 768.515252 | 768.515 | 0.0003 | PS O-34:2 | C40H76NO9P | [M+Na]+ |
| 745.487333 | 745.4861 | 0.0012 | MGDG 32:4 | C41H70O10 | [M+Na]+ |
| 708.549286 | 708.5538 | 0.0045 | LPS O-32:0 | C38H78NO8P | [M+H]+ |
| 840.53823 | 840.5385 | 0.0003 | DGCC 40:11 | C50H75NO8 | [M+Na]+ |
| 838.663779 | 838.666 | 0.0022 | PE O-42:1 | C47H94NO7P | [M+Na]+ |
| 790.515954 | 790.5147 | 0.0012 | PE O-38:5 | C43H78NO7P | [M+K]+ |
| 658.540476 | 658.5252 | 0.0152 | DGCC 27:0 | C37H71NO8 | [M+H]+ |
| 1013.68922 | 1013.6819 | 0.0074 | PI O-45:2 | C54H103O12P | [M+K]+ |
| 951.694 | 951.692 | 0.002 | MGDG 49:9 | C58H94O10 | [M+H]+ |
| 786.525867 | 786.5256 | 0.0003 | PS 34:0 | C40H78NO10P | [M+Na]+ |
| 917.537296 | 917.5419 | 0.0046 | SQDG 40:6 | C49H82O12S | [M+Na]+ |
| 1055.56656 | 1055.562 | 0.0045 | PIP 45:9 | C54H88O16P2 | [M+H]+ |
| 771.534285 | 771.5324 | 0.0019 | TG O-45:9 | C48H76O5 | [M+K]+ |
| 715.510402 | 715.5062 | 0.0042 | DG O-42:9 | C45H72O4 | [M+K]+ |
| 702.505941 | 702.5068 | 0.0009 | PE 33:2 | C38H72NO8P | [M+H]+ |
| 771.556067 | 771.5558 | 0.0003 | TG 47:11 | C50H74O6 | [M+H]+ |
| 1224.58114 | 1224.5556 | 0.0255 | CoA 32:3 | C53H92N7O17P3S | [M+H]+ |
| 784.63865 | 784.6215 | 0.0172 | PE O-40:3 | C45H86NO7P | [M+H]+ |
| 864.537095 | 864.5304 | 0.0067 | PE O-44:10 | C49H80NO7P | [M+K]+ |
| 985.658927 | 985.6657 | 0.0067 | PA 56:11 | C59H95O8P | [M+Na]+ |
| 795.533901 | 795.5324 | 0.0015 | TG O-47:11 | C50H76O5 | [M+K]+ |
| 713.497867 | 713.4963 | 0.0015 | LPI O-27:0 | C36H73O11P | [M+H]+ |
| 841.541849 | 841.5378 | 0.0041 | PG O-43:11 | C49H77O9P | [M+H]+ |
| 853.463086 | 853.4628 | 0.0003 | PI O-34:5 | C43H75O12P | [M+K]+ |
| 604.29959 | 604.301 | 0.0014 | PE 25:6 | C30H48NO8P | [M+Na]+ |
| 600.267418 | 600.2698 | 0.0024 | PS O-21:3 | C27H48NO9P | [M+K]+ |
| 580.30083 | 580.301 | 0.0001 | PE 23:4 | C28H48NO8P | [M+Na]+ |

| | | | | | |
|---|---|---|---|---|---|
| 769.518501 | 769.5168 | 0.0017 | TG O-45:10 | C48H74O5 | [M+K]+ |
| 575.2552 | 575.2534 | 0.0018 | PA O-26:7 | C29H45O7P | [M+K]+ |
| 502.48631 | 502.4618 | 0.0245 | NAE 31:4 | C33H59NO2 | [M+H]+ |
| 691.430481 | 691.4309 | 0.0004 | PA 34:4 | C37H65O8P | [M+Na]+ |
| 692.433612 | 692.4286 | 0.005 | PE 33:7 | C38H62NO8P | [M+H]+ |
| 1045.53824 | 1045.5179 | 0.0203 | PIP 41:5 | C50H88O16P2 | [M+K]+ |
| 719.442306 | 719.4412 | 0.0011 | PA O-36:5 | C39H69O7P | [M+K]+ |
| 666.410235 | 666.4105 | 0.0003 | PE 29:3 | C34H62NO8P | [M+Na]+ |
| 665.377392 | 665.3789 | 0.0015 | PG O-28:5 | C34H59O9P | [M+Na]+ |
| 941.534502 | 941.5419 | 0.0074 | SQDG 42:8 | C51H82O12S | [M+Na]+ |
| 1135.58461 | 1135.5649 | 0.0197 | PIP 48:9 | C57H94O16P2 | [M+K]+ |
| 875.50564 | 875.5046 | 0.001 | PI 34:1 | C43H81O13P | [M+K]+ |
| 1247.56896 | 1247.5573 | 0.0117 | PIP3 47:7 | C56H98O22P4 | [M+H]+ |
| 679.51192 | 679.5062 | 0.0057 | DG O-39:6 | C42H72O4 | [M+K]+ |
| 738.454818 | 738.4471 | 0.0078 | PE 33:3 | C38H70NO8P | [M+K]+ |
| 773.551193 | 773.5481 | 0.0031 | TG O-45:8 | C48H78O5 | [M+K]+ |
| 1225.58619 | 1225.573 | 0.0132 | PIP3 45:4 | C54H100O22P4 | [M+H]+ |
| 686.509771 | 686.5095 | 0.0003 | PE O-31:0 | C36H74NO7P | [M+Na]+ |
| 705.447215 | 705.4467 | 0.0005 | PG O-29:0 | C35H71O9P | [M+K]+ |
| 881.536795 | 881.5327 | 0.0041 | PG 45:12 | C51H77O10P | [M+H]+ |
| 718.330666 | 718.3481 | 0.0174 | PS O-30:7 | C36H58NO9P | [M+K]+ |
| 623.37099 | 623.3708 | 0.0001 | TG 33:6 | C36H56O6 | [M+K]+ |
| 766.499772 | 766.4993 | 0.0004 | PS O-34:3 | C40H74NO9P | [M+Na]+ |
| 659.544353 | 659.5375 | 0.0068 | DG O-37:2 | C40H76O4 | [M+K]+ |
| 674.326034 | 674.3219 | 0.0042 | PE 29:7 | C34H54NO8P | [M+K]+ |
| 701.495981 | 701.4906 | 0.0054 | DG O-41:9 | C44H70O4 | [M+K]+ |
| 1000.5659 | 1000.6038 | 0.0379 | PS 51:12 | C57H88NO10P | [M+Na]+ |
| 747.536783 | 747.5324 | 0.0044 | TG O-43:7 | C46H76O5 | [M+K]+ |
| 751.508337 | 751.5062 | 0.0021 | DG O-45:12 | C48H72O4 | [M+K]+ |
| 934.567277 | 934.5723 | 0.005 | PE 48:10 | C53H86NO8P | [M+K]+ |
| 664.357577 | 664.3585 | 0.0009 | PS O-27:5 | C33H56NO9P | [M+Na]+ |
| 579.289416 | 579.2905 | 0.0011 | LPI O-16:1 | C25H49O11P | [M+Na]+ |
| 718.451033 | 718.4572 | 0.0062 | LPE O-34:6 | C39H70NO6P | [M+K]+ |
| 885.568516 | 885.564 | 0.0045 | PG 45:10 | C51H81O10P | [M+H]+ |
| 689.28944 | 689.2851 | 0.0043 | PG 28:8 | C34H51O10P | [M+K]+ |
| 679.430327 | 679.4309 | 0.0006 | PA 33:3 | C36H65O8P | [M+Na]+ |
| 735.458232 | 735.4571 | 0.0011 | PG O-33:5 | C39H69O9P | [M+Na]+ |
| 760.510459 | 760.5123 | 0.0018 | DGTS 35:8 | C45H71NO7 | [M+Na]+ |
| 1141.60161 | 1141.6118 | 0.0102 | PIP 48:6 | C57H100O16P2 | [M+K]+ |
| 1245.59069 | 1245.5782 | 0.0125 | PIP2 50:8 | C59H101O19P3 | [M+K]+ |
| 897.487179 | 897.4888 | 0.0017 | PI O-39:10 | C48H75O12P | [M+Na]+ |
| 637.386649 | 637.3865 | 0.0002 | TG 34:6 | C37H58O6 | [M+K]+ |
| 583.320838 | 583.3242 | 0.0033 | LPI O-18:2 | C27H51O11P | [M+H]+ |
| 1230.57144 | 1230.6026 | 0.0311 | CoA 32:0 | C53H98N7O17P3S | [M+H]+ |
| 650.342814 | 650.3428 | 0 | PS O-26:5 | C32H54NO9P | [M+Na]+ |

| | | | | | |
|---|---|---|---|---|---|
| 924.510073 | 924.5151 | 0.0051 | PS 44:9 | C50H80NO10P | [M+K]+ |
| 1059.56163 | 1059.5336 | 0.0281 | PIP 42:5 | C51H90O16P2 | [M+K]+ |
| 986.662744 | 986.6611 | 0.0017 | PS O-49:6 | C55H98NO9P | [M+K]+ |
| 687.275049 | 687.2811 | 0.0061 | SQDG 22:3 | C31H52O12S | [M+K]+ |
| 996.73133 | 996.7263 | 0.005 | DGCC 51:10 | C61H99NO8 | [M+Na]+ |
| 1253.58308 | 1253.6018 | 0.0188 | PIP3 45:1 | C54H106O22P4 | [M+Na]+ |
| 760.340902 | 760.3586 | 0.0177 | PS 32:7 | C38H60NO10P | [M+K]+ |
| 1209.58932 | 1209.5782 | 0.0112 | PIP2 47:5 | C56H101O19P3 | [M+K]+ |
| 719.337359 | 719.3321 | 0.0053 | PG 30:7 | C36H57O10P | [M+K]+ |
| 731.463648 | 731.4646 | 0.001 | PA 39:8 | C42H67O8P | [M+H]+ |
| 991.494168 | 991.5002 | 0.006 | SQDG 45:12 | C54H80O12S | [M+K]+ |
| 901.524906 | 901.5227 | 0.0023 | MGDG 43:11 | C52H78O10 | [M+K]+ |
| 915.518523 | 915.5263 | 0.0077 | SQDG 40:7 | C49H80O12S | [M+Na]+ |
| 764.409459 | 764.4262 | 0.0167 | PE 37:10 | C42H64NO8P | [M+Na]+ |
| 822.416284 | 822.4317 | 0.0154 | PS 38:10 | C44H66NO10P | [M+Na]+ |
| 789.438893 | 789.4315 | 0.0074 | PI O-29:2 | C38H71O12P | [M+K]+ |
| 919.514291 | 919.5097 | 0.0046 | PI O-39:7 | C48H81O12P | [M+K]+ |
| 1046.66181 | 1046.6611 | 0.0007 | PS O-54:11 | C60H98NO9P | [M+K]+ |
| 649.41299 | 649.4075 | 0.0055 | PG 27:2 | C33H61O10P | [M+H]+ |
| 801.560209 | 801.5616 | 0.0014 | PG 36:0 | C42H83O10P | [M+Na]+ |
| 810.34142 | 810.3743 | 0.0329 | PS 36:10 | C42H62NO10P | [M+K]+ |
| 688.343685 | 688.3375 | 0.0062 | PE 30:7 | C35H56NO8P | [M+K]+ |
| 1055.57058 | 1055.562 | 0.0085 | PIP 45:9 | C54H88O16P2 | [M+H]+ |
| 1087.55371 | 1087.5649 | 0.0112 | PIP 44:5 | C53H94O16P2 | [M+K]+ |
| 1224.58721 | 1224.5556 | 0.0316 | CoA 32:3 | C53H92N7O17P3S | [M+H]+ |
| 952.551578 | 952.5464 | 0.0051 | PS 46:9 | C52H84NO10P | [M+K]+ |
| 1252.60546 | 1252.5869 | 0.0186 | CoA 34:3 | C55H96N7O17P3S | [M+H]+ |
| 602.28301 | 602.2853 | 0.0023 | PE 25:7 | C30H46NO8P | [M+Na]+ |
| 581.305336 | 581.3061 | 0.0008 | LPI O-16:0 | C25H51O11P | [M+Na]+ |
| 1231.58089 | 1231.5625 | 0.0184 | PIP2 49:8 | C58H99O19P3 | [M+K]+ |
| 662.377971 | 662.3792 | 0.0013 | PE 29:5 | C34H58NO8P | [M+Na]+ |
| 981.528365 | 981.5254 | 0.003 | PI O-44:11 | C53H83O12P | [M+K]+ |
| 709.423942 | 709.4205 | 0.0034 | PA 34:3 | C37H67O8P | [M+K]+ |
| 1361.60306 | 1361.6018 | 0.0012 | PIP3 54:10 | C63H106O22P4 | [M+Na]+ |
| 700.342365 | 700.3375 | 0.0049 | PE 31:8 | C36H56NO8P | [M+K]+ |
| 1073.57091 | 1073.5492 | 0.0217 | PIP 43:5 | C52H92O16P2 | [M+K]+ |
| 653.360939 | 653.3579 | 0.003 | PA 30:3 | C33H59O8P | [M+K]+ |
| 827.64001 | 827.6314 | 0.0086 | DG O-50:9 | C53H88O4 | [M+K]+ |
| 964.522634 | 964.5464 | 0.0238 | PS 47:10 | C53H84NO10P | [M+K]+ |
| 1274.59351 | 1274.5688 | 0.0247 | CoA 34:3 | C55H96N7O17P3S | [M+Na]+ |
| 723.329222 | 723.327 | 0.0022 | LPI O-26:7 | C35H57O11P | [M+K]+ |
| 885.581648 | 885.5827 | 0.0011 | PI O-37:2 | C46H87O12P | [M+Na]+ |
| 830.418867 | 830.4369 | 0.018 | PS 37:7 | C43H70NO10P | [M+K]+ |
| 761.471217 | 761.4728 | 0.0016 | PG O-35:6 | C41H71O9P | [M+Na]+ |
| 1111.55092 | 1111.5649 | 0.014 | PIP 46:7 | C55H94O16P2 | [M+K]+ |

| | | | | | |
|---|---|---|---|---|---|
| 525.257937 | 525.2588 | 0.0008 | PA 22:3 | C25H43O8P | [M+Na]+ |
| 771.427084 | 771.4324 | 0.0053 | SQDG 29:2 | C38H68O12S | [M+Na]+ |
| 802.388234 | 802.4056 | 0.0174 | PS 35:7 | C41H66NO10P | [M+K]+ |
| 887.548418 | 887.541 | 0.0074 | PI O-36:2 | C45H85O12P | [M+K]+ |
| 1383.58286 | 1383.6227 | 0.0399 | PIP3 54:7 | C63H112O22P4 | [M+K]+ |
| 594.279397 | 594.2802 | 0.0008 | PS O-22:5 | C28H46NO9P | [M+Na]+ |
| 1266.56964 | 1266.5428 | 0.0268 | CoA 32:1 | C53H96N7O17P3S | [M+K]+ |
| 994.59526 | 994.5934 | 0.0019 | PS 49:9 | C55H90NO10P | [M+K]+ |
| 844.529958 | 844.5253 | 0.0046 | PE 41:6 | C46H80NO8P | [M+K]+ |
| 765.378767 | 765.374 | 0.0048 | LPI O-29:7 | C38H63O11P | [M+K]+ |
| 905.490628 | 905.4941 | 0.0034 | PI O-38:7 | C47H79O12P | [M+K]+ |
| 1255.56102 | 1255.5601 | 0.0009 | PIP3 44:1 | C53H104O22P4 | [M+K]+ |
| 1391.57778 | 1391.5914 | 0.0137 | PIP3 55:10 | C64H108O22P4 | [M+K]+ |

*Supporting Table 6: List of annotations for signals of the E2 cell line identified as statistically different (ANOVA multiple-sample test, p-value ≤ 0.05) between the naïve and LPS-stimulated state (biological triplicates, each consisting of 22,500 individual mass spectra of multiple individual cells) in positive-ion mode from LIPID MAPS database research with the lowest mass difference for each input mass.*

| Input mass | Matched mass | Δ *m/z* | Name | Formula | Ion |
|---|---|---|---|---|---|
| 758.570202 | 758.5694 | 0.0008 | PE 37:2 | C42H80NO8P | [M+H]+ |
| 975.606757 | 975.6086 | 0.0018 | PG 50:11 | C56H89O10P | [M+Na]+ |
| 976.610407 | 976.6038 | 0.0066 | PS 49:10 | C55H88NO10P | [M+Na]+ |
| 575.503595 | 575.5034 | 0.0002 | MG 34:4 | C37H66O4 | [M+H]+ |
| 627.534763 | 627.5347 | 0.0001 | DG O-38:6 | C41H70O4 | [M+H]+ |
| 576.507276 | 576.5116 | 0.0044 | NAE 33:0 | C35H71NO2 | [M+K]+ |
| 944.602126 | 944.593 | 0.0091 | PE O-50:12 | C55H88NO7P | [M+K]+ |
| 625.518977 | 625.519 | 0.0001 | DG O-38:7 | C41H68O4 | [M+H]+ |
| 628.538283 | 628.5299 | 0.0084 | CAR 33:5 | C40H69NO4 | [M+H]+ |
| 916.569561 | 916.5617 | 0.0079 | PE O-48:12 | C53H84NO7P | [M+K]+ |
| 1111.62253 | 1111.6222 | 0.0003 | PIP 47:6 | C56H98O16P2 | [M+Na]+ |
| 945.605154 | 945.6098 | 0.0046 | SQDG 40:0 | C49H94O12S | [M+K]+ |
| 947.619696 | 947.6138 | 0.0059 | PG 46:5 | C52H93O10P | [M+K]+ |
| 942.585302 | 942.5983 | 0.013 | PE 50:12 | C55H86NO8P | [M+Na]+ |
| 573.487705 | 573.4877 | 0 | MG 34:5 | C37H64O4 | [M+H]+ |
| 660.290268 | 660.2908 | 0.0005 | PS 26:7 | C32H48NO10P | [M+Na]+ |
| 788.556998 | 788.5565 | 0.0005 | PE O-39:5 | C44H80NO7P | [M+Na]+ |
| 917.573182 | 917.5785 | 0.0053 | SQDG 38:0 | C47H90O12S | [M+K]+ |
| 1250.59644 | 1250.5713 | 0.0252 | CoA 34:4 | C55H94N7O17P3S | [M+H]+ |
| 1061.65087 | 1061.6537 | 0.0029 | DGDG 42:5 | C57H98O15 | [M+K]+ |
| 626.522414 | 626.5143 | 0.0081 | CAR 33:6 | C40H67NO4 | [M+H]+ |
| 524.371959 | 524.371 | 0.0009 | CAR 24:5 | C31H51NO4 | [M+Na]+ |
| 740.502022 | 740.4991 | 0.0029 | PE O-34:2 | C39H76NO7P | [M+K]+ |
| 1059.70047 | 1059.7025 | 0.002 | PG 56:11 | C62H101O10P | [M+Na]+ |
| 943.588785 | 943.5941 | 0.0053 | SQDG 40:1 | C49H92O12S | [M+K]+ |
| 1113.63729 | 1113.6379 | 0.0006 | PIP 47:5 | C56H100O16P2 | [M+Na]+ |

| | | | | | |
|---|---|---|---|---|---|
| 589.519099 | 589.519 | 0.0001 | DG O-35:4 | C38H68O4 | [M+H]+ |
| 772.564882 | 772.5617 | 0.0032 | PE O-36:0 | C41H84NO7P | [M+K]+ |
| 685.548731 | 685.5532 | 0.0044 | DG O-39:3 | C42H78O4 | [M+K]+ |
| 618.579361 | 618.5456 | 0.0338 | CAR 32:3 | C39H71NO4 | [M+H]+ |
| 1251.60099 | 1251.5886 | 0.0124 | PIP2 52:11 | C61H99O19P3 | [M+Na]+ |
| 972.632394 | 972.6243 | 0.0081 | PE O-52:12 | C57H92NO7P | [M+K]+ |
| 678.446456 | 678.4469 | 0.0005 | PE O-31:4 | C36H66NO7P | [M+Na]+ |
| 658.274968 | 658.2906 | 0.0156 | PE 28:8 | C33H50NO8P | [M+K]+ |
| 1276.6143 | 1276.5845 | 0.0298 | CoA 34:2 | C55H98N7O17P3S | [M+Na]+ |
| 551.491814 | 551.4823 | 0.0096 | CE 11:2 | C38H62O2 | [M+H]+ |
| 970.617043 | 970.6296 | 0.0126 | PE 52:12 | C57H90NO8P | [M+Na]+ |
| 977.619043 | 977.6196 | 0.0006 | DGDG 39:7 | C54H88O15 | [M+H]+ |
| 657.529981 | 657.5219 | 0.0081 | DG O-37:3 | C40H74O4 | [M+K]+ |
| 1060.70409 | 1060.6977 | 0.0064 | PS 55:10 | C61H100NO10P | [M+Na]+ |
| 1311.72122 | 1311.7214 | 0.0002 | PIP 61:12 | C70H114O16P2 | [M+K]+ |
| 789.560341 | 789.564 | 0.0037 | PG 37:2 | C43H81O10P | [M+H]+ |
| 1248.58123 | 1248.5556 | 0.0256 | CoA 34:5 | C55H92N7O17P3S | [M+H]+ |
| 1277.61839 | 1277.6043 | 0.0141 | PIP3 49:6 | C58H104O22P4 | [M+H]+ |
| 1114.64176 | 1114.6873 | 0.0455 | PS 58:12 | C64H102NO10P | [M+K]+ |
| 823.599189 | 823.6001 | 0.0009 | DG O-50:11 | C53H84O4 | [M+K]+ |
| 675.570249 | 675.5688 | 0.0014 | DG O-38:1 | C41H80O4 | [M+K]+ |
| 1387.61701 | 1387.6175 | 0.0005 | PIP3 56:11 | C65H108O22P4 | [M+Na]+ |
| 609.382905 | 609.3891 | 0.0062 | PA O-29:3 | C32H59O7P | [M+Na]+ |
| 807.613747 | 807.611 | 0.0028 | PG 38:0 | C44H87O10P | [M+H]+ |
| 768.649288 | 768.6348 | 0.0145 | DGCC 35:1 | C45H85NO8 | [M+H]+ |
| 973.635604 | 973.6411 | 0.0055 | SQDG 42:0 | C51H98O12S | [M+K]+ |
| 914.553296 | 914.567 | 0.0137 | PE 48:12 | C53H82NO8P | [M+Na]+ |
| 507.229768 | 507.2272 | 0.0025 | LPA O-22:6 | C25H41O6P | [M+K]+ |
| 1250.59404 | 1250.5713 | 0.0228 | CoA 34:4 | C55H94N7O17P3S | [M+H]+ |
| 772.603976 | 772.6062 | 0.0022 | DGTS 35:2 | C45H83NO7 | [M+Na]+ |
| 971.620301 | 971.6254 | 0.0051 | SQDG 42:1 | C51H96O12S | [M+K]+ |
| 1061.71416 | 1061.7135 | 0.0007 | DGDG 45:7 | C60H100O15 | [M+H]+ |
| 946.615193 | 946.6086 | 0.0065 | PE O-50:11 | C55H90NO7P | [M+K]+ |
| 975.571785 | 975.5723 | 0.0005 | PI O-43:7 | C52H89O12P | [M+K]+ |
| 1060.61425 | 1060.6403 | 0.0261 | PS 54:11 | C60H96NO10P | [M+K]+ |
| 599.529035 | 599.5034 | 0.0257 | DG O-36:6 | C39H66O4 | [M+H]+ |
| 968.598587 | 968.6141 | 0.0155 | PS O-48:8 | C54H92NO9P | [M+K]+ |
| 940.567534 | 940.5828 | 0.0153 | PS O-46:8 | C52H88NO9P | [M+K]+ |
| 1062.65524 | 1062.656 | 0.0007 | PS 54:10 | C60H98NO10P | [M+K]+ |
| 963.71478 | 963.7165 | 0.0017 | SQDG 44:0 | C53H102O12S | [M+H]+ |
| 1413.63338 | 1413.6331 | 0.0002 | PIP3 58:12 | C67H110O22P4 | [M+Na]+ |
| 770.66208 | 770.6504 | 0.0116 | DGCC 35:0 | C45H87NO8 | [M+H]+ |
| 1057.6847 | 1057.6868 | 0.0021 | PG 56:12 | C62H99O10P | [M+Na]+ |
| 922.507102 | 922.4995 | 0.0076 | PS 44:10 | C50H78NO10P | [M+K]+ |
| 650.567254 | 650.5483 | 0.019 | LPE O-31:0 | C36H76NO6P | [M+H]+ |

*Supporting Table 7: List of annotations for signals identified as statistically different (ANOVA multiple-sample test, p-value ≤ 0.01) between the OX1, B8, C12 and E2 cell line (biological triplicates, each consisting of 22,500 individual mass spectra of multiple individual cells) in naïve state in negative-ion mode from LIPID MAPS database research with the lowest mass difference for each input mass.*

| Input mass | Matched mass | Δ $m/z$ | Name | Formula | Ion |
|---|---|---|---|---|---|
| 673.481354 | 673.4814 | 0 | PA 34:1 | C37H71O8P | [M-H]- |
| 699.498311 | 699.4994 | 0.0011 | TG O-43:11 | C46H68O5 | [M-H]- |
| 616.471659 | 616.4712 | 0.0005 | LPE O-29:2 | C34H68NO6P | [M-H]- |
| 642.486975 | 642.4868 | 0.0002 | LPE O-31:3 | C36H70NO6P | [M-H]- |
| 644.502865 | 644.5025 | 0.0004 | LPE O-31:2 | C36H72NO6P | [M-H]- |
| 687.498463 | 687.4994 | 0.0009 | TG O-42:10 | C45H68O5 | [M-H]- |
| 340.202657 | 340.1588 | 0.0438 | NAT 15:4 | C17H27NO4S | [M-H]- |
| 500.277063 | 500.2783 | 0.0012 | PE O-20:4 | C25H44NO7P | [M-H]- |
| 435.250883 | 435.2517 | 0.0008 | LPA 18:1 | C21H41O7P | [M-H]- |
| 773.536046 | 773.5338 | 0.0022 | PG 36:2 | C42H79O10P | [M-H]- |
| 630.487201 | 630.4868 | 0.0004 | LPE O-30:2 | C35H70NO6P | [M-H]- |
| 643.490491 | 643.4943 | 0.0038 | TG 37:4 | C40H68O6 | [M-H]- |
| 917.541329 | 917.5454 | 0.0041 | SQDG 42:8 | C51H82O12S | [M-H]- |
| 711.337808 | 711.342 | 0.0042 | SQDG 27:6 | C36H56O12S | [M-H]- |
| 645.506159 | 645.51 | 0.0038 | TG 37:3 | C40H70O6 | [M-H]- |
| 821.567757 | 821.5702 | 0.0024 | PG O-41:6 | C47H83O9P | [M-H]- |
| 754.614455 | 754.6202 | 0.0058 | DGCC 34:0 | C44H85NO8 | [M-H]- |
| 764.558294 | 764.56 | 0.0017 | PE O-39:5 | C44H80NO7P | [M-H]- |
| 632.502951 | 632.5025 | 0.0005 | LPE O-30:1 | C35H72NO6P | [M-H]- |
| 887.535291 | 887.508 | 0.0273 | PI O-40:10 | C49H77O12P | [M-H]- |
| 885.519832 | 885.4923 | 0.0275 | PI O-40:11 | C49H75O12P | [M-H]- |
| 901.546746 | 901.5236 | 0.0231 | PI O-41:10 | C50H79O12P | [M-H]- |
| 728.597379 | 728.5964 | 0.001 | LPC O-34:2 | C42H84NO6P | [M-H]- |
| 701.490014 | 701.4787 | 0.0113 | TG 42:10 | C45H66O6 | [M-H]- |
| 699.473596 | 699.463 | 0.0106 | TG 42:11 | C45H64O6 | [M-H]- |
| 861.51819 | 861.544 | 0.0258 | PA 49:12 | C52H79O8P | [M-H]- |
| 886.523567 | 886.5604 | 0.0368 | PS 44:8 | C50H82NO10P | [M-H]- |
| 713.330091 | 713.3308 | 0.0007 | PI 26:6 | C35H55O13P | [M-H]- |
| 715.493169 | 715.4943 | 0.0012 | TG 43:10 | C46H68O6 | [M-H]- |
| 883.50313 | 883.5342 | 0.0311 | PI 38:5 | C47H81O13P | [M-H]- |
| 697.336284 | 697.3358 | 0.0004 | PI O-26:7 | C35H55O12P | [M-H]- |
| 888.539399 | 888.576 | 0.0366 | PS 44:7 | C50H84NO10P | [M-H]- |
| 702.492784 | 702.495 | 0.0023 | DGCC 31:5 | C41H69NO8 | [M-H]- |
| 862.52345 | 862.5392 | 0.0158 | PE 46:12 | C51H78NO8P | [M-H]- |
| 700.476601 | 700.4794 | 0.0028 | DGCC 31:6 | C41H67NO8 | [M-H]- |
| 771.520473 | 771.5182 | 0.0023 | PG 36:3 | C42H77O10P | [M-H]- |
| 745.503407 | 745.5025 | 0.0009 | PG 34:2 | C40H75O10P | [M-H]- |
| 688.502092 | 688.4923 | 0.0098 | PE 32:1 | C37H72NO8P | [M-H]- |
| 599.302427 | 599.2991 | 0.0034 | PG 24:5 | C30H49O10P | [M-H]- |
| 597.303788 | 597.3045 | 0.0008 | LPI 18:1 | C27H51O12P | [M-H]- |
| 859.501889 | 859.4767 | 0.0252 | PI O-38:10 | C47H73O12P | [M-H]- |

| | | | | | |
|---|---|---|---|---|---|
| 884.507512 | 884.5447 | 0.0372 | PS 44:9 | C50H80NO10P | [M-H]- |
| 789.521268 | 789.5287 | 0.0075 | LPI O-33:3 | C42H79O11P | [M-H]- |
| 687.523362 | 687.5334 | 0.0101 | PA O-36:1 | C39H77O7P | [M-H]- |
| 616.451576 | 616.4348 | 0.0168 | PE O-28:2 | C33H64NO7P | [M-H]- |
| 674.462052 | 674.4637 | 0.0017 | DGCC 29:5 | C39H65NO8 | [M-H]- |
| 526.292467 | 526.2939 | 0.0015 | PE O-22:5 | C27H46NO7P | [M-H]- |
| 773.513875 | 773.5127 | 0.0012 | PA 42:7 | C45H75O8P | [M-H]- |
| 671.449551 | 671.4657 | 0.0162 | PA 34:2 | C37H69O8P | [M-H]- |
| 687.476311 | 687.463 | 0.0133 | TG 41:10 | C44H64O6 | [M-H]- |
| 760.489201 | 760.4923 | 0.0031 | PE 38:7 | C43H72NO8P | [M-H]- |
| 725.353341 | 725.3576 | 0.0043 | SQDG 28:6 | C37H58O12S | [M-H]- |
| 673.464905 | 673.4814 | 0.0165 | PA 34:1 | C37H71O8P | [M-H]- |
| 498.261457 | 498.2626 | 0.0012 | PE O-20:5 | C25H42NO7P | [M-H]- |
| 618.465613 | 618.4504 | 0.0152 | PE O-28:1 | C33H66NO7P | [M-H]- |
| 1151.6724 | 1151.657 | 0.0154 | PIP 52:9 | C61H102O16P2 | [M-H]- |
| 860.50799 | 860.5447 | 0.0367 | PS 42:7 | C48H80NO10P | [M-H]- |
| 689.537871 | 689.5491 | 0.0112 | PA O-36:0 | C39H79O7P | [M-H]- |
| 753.357686 | 753.3621 | 0.0044 | PI 29:7 | C38H59O13P | [M-H]- |
| 788.516412 | 788.5236 | 0.0072 | PE 40:7 | C45H76NO8P | [M-H]- |
| 863.53501 | 863.5596 | 0.0246 | PA 49:11 | C52H81O8P | [M-H]- |
| 933.50356 | 933.5499 | 0.0463 | PI 42:8 | C51H83O13P | [M-H]- |
| 833.49619 | 833.5127 | 0.0165 | PA 47:12 | C50H75O8P | [M-H]- |
| 744.527929 | 744.5185 | 0.0094 | PS O-34:2 | C40H76NO9P | [M-H]- |
| 902.550049 | 902.5917 | 0.0416 | PS 45:7 | C51H86NO10P | [M-H]- |
| 755.617792 | 755.6195 | 0.0017 | TG 45:4 | C48H84O6 | [M-H]- |
| 530.323728 | 530.3252 | 0.0015 | PE O-22:3 | C27H50NO7P | [M-H]- |
| 786.503641 | 786.5079 | 0.0043 | PE 40:8 | C45H74NO8P | [M-H]- |
| 915.525711 | 915.5298 | 0.0041 | SQDG 42:9 | C51H80O12S | [M-H]- |
| 1087.63027 | 1087.6257 | 0.0045 | PIP 47:6 | C56H98O16P2 | [M-H]- |
| 742.511948 | 742.5029 | 0.0091 | PS O-34:3 | C40H74NO9P | [M-H]- |
| 742.612827 | 742.5756 | 0.0372 | PE O-37:2 | C42H82NO7P | [M-H]- |
| 605.327457 | 605.3249 | 0.0026 | PA 30:7 | C33H51O8P | [M-H]- |
| 711.314892 | 711.3151 | 0.0002 | PI 26:7 | C35H53O13P | [M-H]- |
| 723.350272 | 723.3515 | 0.0012 | PI O-28:8 | C37H57O12P | [M-H]- |
| 889.545391 | 889.5236 | 0.0217 | PI O-40:9 | C49H79O12P | [M-H]- |
| 645.432726 | 645.4161 | 0.0167 | TG 38:10 | C41H58O6 | [M-H]- |
| 935.518205 | 935.5655 | 0.0473 | PI 42:7 | C51H85O13P | [M-H]- |
| 917.506439 | 917.5162 | 0.0098 | PIP 34:0 | C43H84O16P2 | [M-H]- |
| 864.541225 | 864.5549 | 0.0137 | PE 46:11 | C51H80NO8P | [M-H]- |
| 787.505698 | 787.5131 | 0.0074 | LPI O-33:4 | C42H77O11P | [M-H]- |
| 1085.6146 | 1085.6101 | 0.0045 | PIP 47:7 | C56H96O16P2 | [M-H]- |
| 1153.68063 | 1153.6727 | 0.0079 | PIP 52:8 | C61H104O16P2 | [M-H]- |
| 835.503827 | 835.5283 | 0.0245 | PA 47:11 | C50H77O8P | [M-H]- |
| 688.528624 | 688.5287 | 0 | PE O-33:1 | C38H76NO7P | [M-H]- |
| 772.523869 | 772.5287 | 0.0048 | PE O-40:8 | C45H76NO7P | [M-H]- |

| | | | | | |
|---|---|---|---|---|---|
| 765.561638 | 765.5522 | 0.0094 | MGDG 35:3 | C44H78O10 | [M-H]- |
| 647.448674 | 647.4317 | 0.017 | TG 38:9 | C41H60O6 | [M-H]- |
| 476.277179 | 476.2783 | 0.0011 | LPE 18:2 | C23H44NO7P | [M-H]- |
| 761.497103 | 761.4974 | 0.0003 | LPI O-31:3 | C40H75O11P | [M-H]- |
| 899.531137 | 899.508 | 0.0231 | PI O-41:11 | C50H77O12P | [M-H]- |
| 857.488328 | 857.5186 | 0.0302 | PI 36:4 | C45H79O13P | [M-H]- |
| 725.36757 | 725.3671 | 0.0004 | PI O-28:7 | C37H59O12P | [M-H]- |
| 767.243723 | 767.2814 | 0.0377 | PIP 24:5 | C33H54O16P2 | [M-H]- |
| 797.535351 | 797.5338 | 0.0015 | PG 38:4 | C44H79O10P | [M-H]- |
| 478.27909 | 478.2939 | 0.0148 | LPE 18:1 | C23H46NO7P | [M-H]- |
| 770.508151 | 770.513 | 0.0049 | PE O-40:9 | C45H74NO7P | [M-H]- |
| 931.487755 | 931.5318 | 0.0441 | PIP 35:0 | C44H86O16P2 | [M-H]- |
| 501.280483 | 501.2987 | 0.0182 | PA O-23:3 | C26H47O7P | [M-H]- |
| 797.632327 | 797.643 | 0.0106 | PA O-44:2 | C47H91O7P | [M-H]- |
| 715.468558 | 715.4708 | 0.0023 | PA O-39:8 | C42H69O7P | [M-H]- |
| 919.522069 | 919.5342 | 0.0121 | PI 41:8 | C50H81O13P | [M-H]- |
| 834.493531 | 834.5079 | 0.0144 | PE 44:12 | C49H74NO8P | [M-H]- |
| 457.235127 | 457.2361 | 0.0009 | PA O-20:4 | C23H39O7P | [M-H]- |
| 790.525263 | 790.5263 | 0.0011 | DGCC 38:10 | C48H73NO8 | [M-H]- |
| 934.511014 | 934.5604 | 0.0494 | PS 48:12 | C54H82NO10P | [M-H]- |
| 1059.60218 | 1059.5944 | 0.0077 | PIP 45:6 | C54H94O16P2 | [M-H]- |
| 433.235213 | 433.2361 | 0.0009 | LPA 18:2 | C21H39O7P | [M-H]- |
| 703.495575 | 703.4943 | 0.0013 | TG 42:9 | C45H68O6 | [M-H]- |
| 461.266435 | 461.2674 | 0.0009 | PA O-20:2 | C23H43O7P | [M-H]- |
| 689.477758 | 689.4787 | 0.0009 | TG 41:9 | C44H66O6 | [M-H]- |
| 822.56969 | 822.5655 | 0.0042 | PS O-40:5 | C46H82NO9P | [M-H]- |
| 819.552138 | 819.5546 | 0.0024 | PG O-41:7 | C47H81O9P | [M-H]- |
| 740.500402 | 740.5107 | 0.0103 | DGCC 34:7 | C44H71NO8 | [M-H]- |
| 799.551337 | 799.5495 | 0.0019 | PG 38:3 | C44H81O10P | [M-H]- |
| 901.515646 | 901.5141 | 0.0015 | SQDG 41:9 | C50H78O12S | [M-H]- |
| 690.545536 | 690.5443 | 0.0012 | PE O-33:0 | C38H78NO7P | [M-H]- |
| 678.493534 | 678.495 | 0.0015 | DGCC 29:3 | C39H69NO8 | [M-H]- |
| 579.312226 | 579.3092 | 0.003 | PA 28:6 | C31H49O8P | [M-H]- |
| 741.358124 | 741.3621 | 0.0039 | PI 28:6 | C37H59O13P | [M-H]- |
| 678.506976 | 678.5079 | 0.001 | LPS O-30:0 | C36H74NO8P | [M-H]- |
| 836.510425 | 836.5236 | 0.0132 | PE 44:11 | C49H76NO8P | [M-H]- |
| 743.487648 | 743.4869 | 0.0008 | PG 34:3 | C40H73O10P | [M-H]- |
| 717.490687 | 717.4865 | 0.0042 | PA O-39:7 | C42H71O7P | [M-H]- |
| 874.668132 | 874.6695 | 0.0014 | PE O-47:6 | C52H94NO7P | [M-H]- |
| 728.533199 | 728.5236 | 0.0096 | PE 35:2 | C40H76NO8P | [M-H]- |
| 600.306823 | 600.2943 | 0.0125 | PS 23:4 | C29H48NO10P | [M-H]- |
| 799.647719 | 799.6586 | 0.0109 | PA O-44:1 | C47H93O7P | [M-H]- |
| 633.360988 | 633.3644 | 0.0034 | MGDG 26:6 | C35H54O10 | [M-H]- |
| 758.473497 | 758.4766 | 0.0031 | PE 38:8 | C43H70NO8P | [M-H]- |
| 678.512611 | 678.5079 | 0.0047 | LPS O-30:0 | C36H74NO8P | [M-H]- |

| | | | | | |
|---|---|---|---|---|---|
| 767.577338 | 767.5679 | 0.0095 | MGDG 35:2 | C44H80O10 | [M-H]- |
| 528.308099 | 528.3096 | 0.0015 | PE O-22:4 | C27H48NO7P | [M-H]- |
| 509.287101 | 509.2885 | 0.0014 | LPG 18:1 | C24H47O9P | [M-H]- |
| 694.369136 | 694.3726 | 0.0034 | PS 30:6 | C36H58NO10P | [M-H]- |
| 459.250772 | 459.2517 | 0.001 | PA O-20:3 | C23H41O7P | [M-H]- |
| 593.328298 | 593.3249 | 0.0034 | PA 29:6 | C32H51O8P | [M-H]- |
| 903.526785 | 903.5298 | 0.003 | SQDG 41:8 | C50H80O12S | [M-H]- |
| 784.551678 | 784.5498 | 0.0019 | PS O-37:3 | C43H80NO9P | [M-H]- |
| 650.3778 | 650.3827 | 0.0049 | PE 30:6 | C35H58NO8P | [M-H]- |
| 915.494031 | 915.5005 | 0.0065 | PIP 34:1 | C43H82O16P2 | [M-H]- |
| 814.529826 | 814.5392 | 0.0094 | PE 42:8 | C47H78NO8P | [M-H]- |
| 798.567157 | 798.5655 | 0.0017 | PS O-38:3 | C44H82NO9P | [M-H]- |
| 563.318058 | 563.3143 | 0.0037 | PA O-28:7 | C31H49O7P | [M-H]- |
| 548.26131 | 548.2994 | 0.0381 | PS O-20:2 | C26H48NO9P | [M-H]- |
| 739.346157 | 739.3464 | 0.0003 | PI 28:7 | C37H57O13P | [M-H]- |
| 803.582533 | 803.5808 | 0.0018 | PG 38:1 | C44H85O10P | [M-H]- |
| 816.512335 | 816.5185 | 0.0062 | PS O-40:8 | C46H76NO9P | [M-H]- |
| 591.348098 | 591.3456 | 0.0025 | PA O-30:7 | C33H53O7P | [M-H]- |
| 504.308372 | 504.3096 | 0.0012 | PE O-20:2 | C25H48NO7P | [M-H]- |
| 695.318157 | 695.3107 | 0.0075 | SQDG 26:7 | C35H52O12S | [M-H]- |
| 900.534442 | 900.576 | 0.0416 | PS 45:8 | C51H84NO10P | [M-H]- |
| 744.491369 | 744.4974 | 0.006 | PE O-38:8 | C43H72NO7P | [M-H]- |
| 784.490009 | 784.4923 | 0.0023 | PE 40:9 | C45H72NO8P | [M-H]- |
| 763.378227 | 763.3733 | 0.0049 | SQDG 31:8 | C40H60O12S | [M-H]- |
| 817.570996 | 817.5753 | 0.0043 | PA 45:6 | C48H83O8P | [M-H]- |
| 774.504983 | 774.5079 | 0.003 | PE 39:7 | C44H74NO8P | [M-H]- |
| 761.536116 | 761.5338 | 0.0023 | PG 35:1 | C41H79O10P | [M-H]- |
| 876.683716 | 876.6852 | 0.0015 | PE O-47:5 | C52H96NO7P | [M-H]- |
| 675.462967 | 675.463 | 0.0001 | TG 40:9 | C43H64O6 | [M-H]- |
| 589.334754 | 589.33 | 0.0048 | PA O-30:8 | C33H51O7P | [M-H]- |
| 899.504094 | 899.508 | 0.0039 | PI O-41:11 | C50H77O12P | [M-H]- |
| 798.635315 | 798.6382 | 0.0029 | PE O-41:2 | C46H90NO7P | [M-H]- |
| 920.526963 | 920.5447 | 0.0178 | PS 47:12 | C53H80NO10P | [M-H]- |
| 680.480439 | 680.4661 | 0.0144 | PE O-33:5 | C38H68NO7P | [M-H]- |
| 473.281529 | 473.2674 | 0.0142 | PA O-21:3 | C24H43O7P | [M-H]- |
| 616.374786 | 616.3855 | 0.0107 | DGCC 25:6 | C35H55NO8 | [M-H]- |
| 481.298161 | 481.2936 | 0.0046 | LPG O-17:1 | C23H47O8P | [M-H]- |
| 751.340969 | 751.3464 | 0.0054 | PI 29:8 | C38H57O13P | [M-H]- |
| 802.530082 | 802.5263 | 0.0037 | DGCC 39:11 | C49H73NO8 | [M-H]- |
| 487.281874 | 487.283 | 0.0011 | PA O-22:3 | C25H45O7P | [M-H]- |
| 787.515697 | 787.5131 | 0.0026 | LPI O-33:4 | C42H77O11P | [M-H]- |
| 1087.71523 | 1087.722 | 0.0068 | PI 53:8 | C62H105O13P | [M-H]- |
| 678.521341 | 678.5314 | 0.0101 | DGTS 30:2 | C40H73NO7 | [M-H]- |
| 856.509553 | 856.5134 | 0.0039 | PS 42:9 | C48H76NO10P | [M-H]- |
| 791.374745 | 791.3753 | 0.0006 | PIP 25:0 | C34H66O16P2 | [M-H]- |

| | | | | | |
|---|---|---|---|---|---|
| 1175.66238 | 1175.657 | 0.0053 | PIP 54:11 | C63H102O16P2 | [M-H]- |
| 749.367549 | 749.3576 | 0.0099 | SQDG 30:8 | C39H58O12S | [M-H]- |
| 814.497427 | 814.5029 | 0.0054 | PS O-40:9 | C46H74NO9P | [M-H]- |
| 648.363632 | 648.3671 | 0.0035 | PE 30:7 | C35H56NO8P | [M-H]- |
| 646.432595 | 646.4324 | 0.0001 | DGCC 27:5 | C37H61NO8 | [M-H]- |
| 789.531725 | 789.5287 | 0.003 | LPI O-33:3 | C42H79O11P | [M-H]- |
| 847.504105 | 847.5283 | 0.0242 | PA 48:12 | C51H77O8P | [M-H]- |
| 564.25749 | 564.2943 | 0.0368 | PS 20:1 | C26H48NO10P | [M-H]- |
| 598.307195 | 598.3151 | 0.0079 | PS O-24:5 | C30H50NO9P | [M-H]- |
| 835.37424 | 835.344 | 0.0302 | PIP 29:6 | C38H62O16P2 | [M-H]- |
| 765.355607 | 765.3621 | 0.0065 | PI 30:8 | C39H59O13P | [M-H]- |
| 804.541213 | 804.542 | 0.0008 | DGCC 39:10 | C49H75NO8 | [M-H]- |
| 689.454372 | 689.4552 | 0.0008 | PA O-37:7 | C40H67O7P | [M-H]- |
| 683.318669 | 683.3202 | 0.0015 | PI O-25:7 | C34H53O12P | [M-H]- |
| 483.250551 | 483.2517 | 0.0012 | PA O-22:5 | C25H41O7P | [M-H]- |
| 527.29592 | 527.2779 | 0.018 | PA 24:4 | C27H45O8P | [M-H]- |
| 1073.70018 | 1073.7064 | 0.0062 | PI 52:8 | C61H103O13P | [M-H]- |
| 551.281429 | 551.2779 | 0.0035 | PA 26:6 | C29H45O8P | [M-H]- |
| 696.470589 | 696.461 | 0.0096 | PE 33:4 | C38H68NO8P | [M-H]- |
| 761.390271 | 761.4247 | 0.0344 | PI 29:3 | C38H67O13P | [M-H]- |
| 535.285835 | 535.283 | 0.0028 | PA O-26:7 | C29H45O7P | [M-H]- |
| 865.547685 | 865.5389 | 0.0088 | PG O-45:12 | C51H79O9P | [M-H]- |
| 875.498419 | 875.4985 | 0.0001 | SQDG 39:8 | C48H76O12S | [M-H]- |
| 742.360585 | 742.409 | 0.0484 | PS O-35:10 | C41H62NO9P | [M-H]- |
| 499.264829 | 499.283 | 0.0182 | PA O-23:4 | C26H45O7P | [M-H]- |
| 532.266557 | 532.2681 | 0.0015 | LPS 19:3 | C25H44NO9P | [M-H]- |
| 606.327805 | 606.3201 | 0.0077 | PE 27:7 | C32H50NO8P | [M-H]- |
| 741.489107 | 741.4865 | 0.0026 | PA O-41:9 | C44H71O7P | [M-H]- |
| 1089.6385 | 1089.6414 | 0.0029 | PIP 47:5 | C56H100O16P2 | [M-H]- |
| 726.372779 | 726.414 | 0.0413 | PE 36:10 | C41H62NO8P | [M-H]- |
| 781.388694 | 781.3934 | 0.0047 | PI 31:7 | C40H63O13P | [M-H]- |
| 737.366421 | 737.3671 | 0.0007 | PI O-29:8 | C38H59O12P | [M-H]- |
| 776.546556 | 776.5471 | 0.0005 | DGTS 38:9 | C48H75NO7 | [M-H]- |
| 678.315007 | 678.3413 | 0.0263 | PS 29:7 | C35H54NO10P | [M-H]- |
| 849.517554 | 849.4923 | 0.0252 | PI O-37:8 | C46H75O12P | [M-H]- |
| 807.552528 | 807.5546 | 0.002 | PG O-40:6 | C46H81O9P | [M-H]- |
| 800.653916 | 800.6539 | 0 | PE O-41:1 | C46H92NO7P | [M-H]- |
| 763.514272 | 763.5131 | 0.0012 | LPI O-31:2 | C40H77O11P | [M-H]- |
| 779.373133 | 779.3777 | 0.0046 | PI 31:8 | C40H61O13P | [M-H]- |
| 773.520462 | 773.5209 | 0.0005 | MGDG 36:6 | C45H74O10 | [M-H]- |
| 452.277352 | 452.2783 | 0.0009 | LPE 16:0 | C21H44NO7P | [M-H]- |
| 710.587043 | 710.594 | 0.007 | DGTS 32:0 | C42H81NO7 | [M-H]- |
| 648.449355 | 648.4481 | 0.0013 | DGCC 27:4 | C37H63NO8 | [M-H]- |
| 799.570185 | 799.5647 | 0.0055 | PA O-45:8 | C48H81O7P | [M-H]- |
| 784.47776 | 784.4923 | 0.0145 | PE 40:9 | C45H72NO8P | [M-H]- |

| | | | | | |
|---|---|---|---|---|---|
| 449.266413 | 449.2674 | 0.001 | LPA 19:1 | C22H43O7P | [M-H]- |
| 750.58276 | 750.5889 | 0.0062 | DGCC 34:2 | C44H81NO8 | [M-H]- |
| 787.551553 | 787.5495 | 0.0021 | PG 37:2 | C43H81O10P | [M-H]- |
| 596.263738 | 596.263 | 0.0007 | PS 23:6 | C29H44NO10P | [M-H]- |
| 1179.68863 | 1179.6883 | 0.0003 | PIP 54:9 | C63H106O16P2 | [M-H]- |
| 650.344828 | 650.3464 | 0.0015 | PS O-28:7 | C34H54NO9P | [M-H]- |
| 1088.72022 | 1088.7325 | 0.0123 | PS 59:12 | C65H104NO10P | [M-H]- |
| 577.334959 | 577.33 | 0.005 | PA O-29:7 | C32H51O7P | [M-H]- |
| 476.262592 | 476.2783 | 0.0157 | LPE 18:2 | C23H44NO7P | [M-H]- |
| 727.34718 | 727.3464 | 0.0008 | PI 27:6 | C36H57O13P | [M-H]- |
| 353.214983 | 353.2099 | 0.0051 | LPA O-13:0 | C16H35O6P | [M-H]- |
| 617.359792 | 617.3613 | 0.0015 | PA O-32:8 | C35H55O7P | [M-H]- |
| 715.333597 | 715.3464 | 0.0128 | PI 26:5 | C35H57O13P | [M-H]- |
| 546.245641 | 546.2838 | 0.0381 | PS O-20:3 | C26H46NO9P | [M-H]- |
| 660.497379 | 660.4974 | 0 | PE O-31:1 | C36H72NO7P | [M-H]- |
| 531.327065 | 531.3092 | 0.0178 | PA 24:2 | C27H49O8P | [M-H]- |
| 600.293412 | 600.2943 | 0.0009 | PS 23:4 | C29H48NO10P | [M-H]- |
| 789.566679 | 789.5651 | 0.0016 | PG 37:1 | C43H83O10P | [M-H]- |
| 690.415619 | 690.414 | 0.0016 | PE 33:7 | C38H62NO8P | [M-H]- |
| 755.52511 | 755.5256 | 0.0005 | TG 46:11 | C49H72O6 | [M-H]- |
| 1101.646 | 1101.6414 | 0.0046 | PIP 48:6 | C57H100O16P2 | [M-H]- |
| 678.305917 | 678.3413 | 0.0354 | PS 29:7 | C35H54NO10P | [M-H]- |
| 762.531836 | 762.5314 | 0.0004 | DGTS 37:9 | C47H73NO7 | [M-H]- |
| 890.553307 | 890.5705 | 0.0172 | PE 48:12 | C53H82NO8P | [M-H]- |
| 1017.64545 | 1017.6438 | 0.0017 | PI 48:8 | C57H95O13P | [M-H]- |
| 733.480385 | 733.4814 | 0.001 | PA 39:6 | C42H71O8P | [M-H]- |
| 662.382235 | 662.3827 | 0.0005 | PE 31:7 | C36H58NO8P | [M-H]- |
| 807.478447 | 807.497 | 0.0186 | PA 45:11 | C48H73O8P | [M-H]- |
| 689.504711 | 689.5127 | 0.008 | PA 35:0 | C38H75O8P | [M-H]- |
| 786.56749 | 786.5655 | 0.002 | PS O-37:2 | C43H82NO9P | [M-H]- |
| 893.506591 | 893.5186 | 0.012 | PI 39:7 | C48H79O13P | [M-H]- |
| 751.386442 | 751.3828 | 0.0036 | PI O-30:8 | C39H61O12P | [M-H]- |
| 565.299151 | 565.3018 | 0.0027 | MGDG 21:5 | C30H46O10 | [M-H]- |
| 826.536638 | 826.5392 | 0.0026 | PE 43:9 | C48H78NO8P | [M-H]- |
| 1018.65663 | 1018.6543 | 0.0024 | PS 54:12 | C60H94NO10P | [M-H]- |
| 524.277041 | 524.2783 | 0.0012 | PE O-22:6 | C27H44NO7P | [M-H]- |
| 875.671807 | 875.6747 | 0.0029 | PG 43:0 | C49H97O10P | [M-H]- |
| 748.567449 | 748.5733 | 0.0058 | DGCC 34:3 | C44H79NO8 | [M-H]- |
| 1089.73098 | 1089.7377 | 0.0067 | PI 53:7 | C62H107O13P | [M-H]- |
| 678.294209 | 678.3413 | 0.0471 | PS 29:7 | C35H54NO10P | [M-H]- |
| 772.488436 | 772.4923 | 0.0039 | PE 39:8 | C44H72NO8P | [M-H]- |
| 771.292281 | 771.3127 | 0.0205 | PIP 24:3 | C33H58O16P2 | [M-H]- |
| 1173.64983 | 1173.6414 | 0.0084 | PIP 54:12 | C63H100O16P2 | [M-H]- |
| 619.304714 | 619.2889 | 0.0158 | PI O-20:4 | C29H49O12P | [M-H]- |
| 704.503378 | 704.5025 | 0.0009 | LPC O-33:7 | C41H72NO6P | [M-H]- |

| | | | | | |
|---|---|---|---|---|---|
| 831.479887 | 831.5029 | 0.023 | PI 34:3 | C43H77O13P | [M-H]- |
| 775.505878 | 775.5131 | 0.0072 | LPI O-32:3 | C41H77O11P | [M-H]- |
| 577.297732 | 577.3018 | 0.0041 | MGDG 22:6 | C31H46O10 | [M-H]- |
| 676.361387 | 676.362 | 0.0006 | PS O-30:8 | C36H56NO9P | [M-H]- |
| 784.522331 | 784.5287 | 0.0063 | PE O-41:9 | C46H76NO7P | [M-H]- |
| 842.522802 | 842.5342 | 0.0114 | PS O-42:9 | C48H78NO9P | [M-H]- |
| 666.341306 | 666.3413 | 0 | PS 28:6 | C34H54NO10P | [M-H]- |
| 678.291522 | 678.3413 | 0.0497 | PS 29:7 | C35H54NO10P | [M-H]- |
| 685.298856 | 685.2995 | 0.0006 | PI 24:6 | C33H51O13P | [M-H]- |
| 815.537 | 815.5233 | 0.0137 | PG O-41:9 | C47H77O9P | [M-H]- |
| 837.514323 | 837.5076 | 0.0067 | PG O-43:12 | C49H75O9P | [M-H]- |
| 800.514436 | 800.5236 | 0.0092 | PE 41:8 | C46H76NO8P | [M-H]- |
| 729.508613 | 729.5076 | 0.001 | PG O-34:3 | C40H75O9P | [M-H]- |
| 799.233922 | 799.2478 | 0.0139 | PIP2 20:1 | C29H55O19P3 | [M-H]- |
| 680.46119 | 680.4661 | 0.0049 | PE O-33:5 | C38H68NO7P | [M-H]- |
| 761.500449 | 761.4974 | 0.003 | LPI O-31:3 | C40H75O11P | [M-H]- |
| 633.433008 | 633.429 | 0.0041 | LPA O-34:7 | C37H63O6P | [M-H]- |
| 561.301923 | 561.2987 | 0.0033 | PA O-28:8 | C31H47O7P | [M-H]- |
| 675.474349 | 675.463 | 0.0113 | TG 40:9 | C43H64O6 | [M-H]- |
| 711.461682 | 711.4607 | 0.001 | PG O-33:5 | C39H69O9P | [M-H]- |
| 542.287151 | 542.2888 | 0.0017 | PE 22:4 | C27H46NO8P | [M-H]- |
| 788.574603 | 788.5811 | 0.0065 | PS O-37:1 | C43H84NO9P | [M-H]- |
| 1074.70418 | 1074.7169 | 0.0127 | PS 58:12 | C64H102NO10P | [M-H]- |
| 891.492407 | 891.5029 | 0.0105 | PI 39:8 | C48H77O13P | [M-H]- |
| 1003.63032 | 1003.6281 | 0.0022 | PI 47:8 | C56H93O13P | [M-H]- |
| 522.282259 | 522.2838 | 0.0015 | LPS 18:1 | C24H46NO9P | [M-H]- |
| 455.219622 | 455.2204 | 0.0008 | PA O-20:5 | C23H37O7P | [M-H]- |
| 574.278063 | 574.2787 | 0.0006 | PS 21:3 | C27H46NO10P | [M-H]- |
| 805.525201 | 805.5236 | 0.0016 | PI O-33:2 | C42H79O12P | [M-H]- |
| 375.315811 | 375.3269 | 0.011 | FA 25:3 | C25H44O2 | [M-H]- |
| 660.363338 | 660.3671 | 0.0037 | PE 31:8 | C36H56NO8P | [M-H]- |
| 755.37271 | 755.3777 | 0.005 | PI 29:6 | C38H61O13P | [M-H]- |
| 812.481066 | 812.4872 | 0.0061 | PS O-40:10 | C46H72NO9P | [M-H]- |
| 516.271819 | 516.2732 | 0.0014 | PE 20:3 | C25H44NO8P | [M-H]- |
| 709.334222 | 709.3263 | 0.0079 | SQDG 27:7 | C36H54O12S | [M-H]- |
| 562.241794 | 562.2787 | 0.0369 | PS 20:2 | C26H46NO10P | [M-H]- |
| 575.318166 | 575.3143 | 0.0038 | PA O-29:8 | C32H49O7P | [M-H]- |
| 544.30282 | 544.3045 | 0.0017 | PE 22:3 | C27H48NO8P | [M-H]- |
| 546.318419 | 546.3201 | 0.0017 | PE 22:2 | C27H50NO8P | [M-H]- |
| 753.401622 | 753.3984 | 0.0032 | PI O-30:7 | C39H63O12P | [M-H]- |
| 756.629893 | 756.5913 | 0.0386 | PE O-38:2 | C43H84NO7P | [M-H]- |
| 788.481097 | 788.4872 | 0.0061 | PS O-38:8 | C44H72NO9P | [M-H]- |
| 845.569854 | 845.5702 | 0.0003 | PG O-43:8 | C49H83O9P | [M-H]- |
| 563.2826 | 563.2779 | 0.0047 | PA 27:7 | C30H45O8P | [M-H]- |
| 791.594019 | 791.596 | 0.002 | PA O-44:5 | C47H85O7P | [M-H]- |

| | | | | | |
|---|---|---|---|---|---|
| 787.494047 | 787.492 | 0.0021 | PG O-39:9 | C45H73O9P | [M-H]- |
| 728.520308 | 728.5236 | 0.0033 | PE 35:2 | C40H76NO8P | [M-H]- |
| 549.301874 | 549.2987 | 0.0032 | PA O-27:7 | C30H47O7P | [M-H]- |
| 785.536085 | 785.5338 | 0.0023 | PG 37:3 | C43H79O10P | [M-H]- |
| 478.242459 | 478.2575 | 0.0151 | LPS O-16:2 | C22H42NO8P | [M-H]- |
| 375.321567 | 375.3269 | 0.0053 | FA 25:3 | C25H44O2 | [M-H]- |
| 504.295774 | 504.3096 | 0.0138 | PE O-20:2 | C25H48NO7P | [M-H]- |
| 760.552283 | 760.5498 | 0.0025 | PS O-35:1 | C41H80NO9P | [M-H]- |
| 375.286437 | 375.2905 | 0.004 | MG O-21:5 | C24H40O3 | [M-H]- |
| 658.453059 | 658.4453 | 0.0077 | PE 30:2 | C35H66NO8P | [M-H]- |
| 375.329501 | 375.3269 | 0.0026 | FA 25:3 | C25H44O2 | [M-H]- |
| 669.306616 | 669.295 | 0.0116 | SQDG 24:6 | C33H50O12S | [M-H]- |
| 913.510199 | 913.5141 | 0.0039 | SQDG 42:10 | C51H78O12S | [M-H]- |
| 1004.6414 | 1004.6386 | 0.0028 | PS 53:12 | C59H92NO10P | [M-H]- |
| 1031.66067 | 1031.6594 | 0.0013 | PI 49:8 | C58H97O13P | [M-H]- |
| 718.492726 | 718.5029 | 0.0101 | PS O-32:1 | C38H74NO9P | [M-H]- |
| 676.397485 | 676.3984 | 0.0009 | PE 32:7 | C37H60NO8P | [M-H]- |
| 711.353035 | 711.3515 | 0.0015 | PI O-27:7 | C36H57O12P | [M-H]- |
| 801.247823 | 801.2634 | 0.0156 | PIP2 20:0 | C29H57O19P3 | [M-H]- |
| 375.276935 | 375.2905 | 0.0135 | MG O-21:5 | C24H40O3 | [M-H]- |
| 843.505783 | 843.5029 | 0.0029 | PI 35:4 | C44H77O13P | [M-H]- |
| 1075.71529 | 1075.722 | 0.0067 | PI 52:7 | C61H105O13P | [M-H]- |
| 893.583968 | 893.5702 | 0.0138 | PG O-47:12 | C53H83O9P | [M-H]- |
| 465.288521 | 465.2987 | 0.0102 | PA O-20:0 | C23H47O7P | [M-H]- |
| 785.555182 | 785.5491 | 0.0061 | PA O-44:8 | C47H79O7P | [M-H]- |
| 549.264924 | 549.2623 | 0.0026 | PA 26:7 | C29H43O8P | [M-H]- |
| 861.404439 | 861.3597 | 0.0447 | PIP 31:7 | C40H64O16P2 | [M-H]- |
| 588.351628 | 588.3542 | 0.0026 | DGCC 23:6 | C33H51NO8 | [M-H]- |
| 375.27011 | 375.2541 | 0.016 | MG 20:5 | C23H36O4 | [M-H]- |
| 1085.65697 | 1085.6101 | 0.0469 | PIP 47:7 | C56H96O16P2 | [M-H]- |
| 375.338725 | 375.3269 | 0.0119 | FA 25:3 | C25H44O2 | [M-H]- |
| 375.353711 | 375.3269 | 0.0269 | FA 25:3 | C25H44O2 | [M-H]- |
| 840.51202 | 840.5185 | 0.0065 | PS O-42:10 | C48H76NO9P | [M-H]- |
| 439.224662 | 439.2103 | 0.0144 | LPG 13:1 | C19H37O9P | [M-H]- |
| 375.257881 | 375.2541 | 0.0038 | MG 20:5 | C23H36O4 | [M-H]- |
| 1061.60968 | 1061.6101 | 0.0004 | PIP 45:5 | C54H96O16P2 | [M-H]- |
| 634.35451 | 634.3514 | 0.0031 | PE 29:7 | C34H54NO8P | [M-H]- |
| 817.521999 | 817.5236 | 0.0016 | PI O-34:3 | C43H79O12P | [M-H]- |
| 375.376166 | 375.3269 | 0.0493 | FA 25:3 | C25H44O2 | [M-H]- |
| 458.238495 | 458.2313 | 0.0072 | LPE 17:4 | C22H38NO7P | [M-H]- |
| 739.385294 | 739.3828 | 0.0025 | PI O-29:7 | C38H61O12P | [M-H]- |
| 624.294399 | 624.2943 | 0.0001 | PS 25:6 | C31H48NO10P | [M-H]- |
| 785.498643 | 785.4974 | 0.0012 | LPI O-33:5 | C42H75O11P | [M-H]- |
| 375.345664 | 375.3269 | 0.0188 | FA 25:3 | C25H44O2 | [M-H]- |
| 929.472516 | 929.5162 | 0.0437 | PIP 35:1 | C44H84O16P2 | [M-H]- |

| | | | | | |
|---|---|---|---|---|---|
| 568.215408 | 568.2317 | 0.0163 | PS 21:6 | C27H40NO10P | [M-H]- |
| 375.361987 | 375.3269 | 0.0351 | FA 25:3 | C25H44O2 | [M-H]- |
| 761.514621 | 761.5127 | 0.0019 | PA 41:6 | C44H75O8P | [M-H]- |
| 877.687222 | 877.6774 | 0.0098 | MGDG 43:3 | C52H94O10 | [M-H]- |
| 341.222192 | 341.2486 | 0.0264 | FA 23:6 | C23H34O2 | [M-H]- |
| 520.302949 | 520.3045 | 0.0015 | PE 20:1 | C25H48NO8P | [M-H]- |
| 632.373371 | 632.3722 | 0.0012 | PE O-30:8 | C35H56NO7P | [M-H]- |
| 375.370204 | 375.3269 | 0.0433 | FA 25:3 | C25H44O2 | [M-H]- |
| 375.239332 | 375.2541 | 0.0148 | MG 20:5 | C23H36O4 | [M-H]- |
| 732.466019 | 732.461 | 0.005 | PE 36:7 | C41H68NO8P | [M-H]- |
| 375.250144 | 375.2541 | 0.0039 | MG 20:5 | C23H36O4 | [M-H]- |
| 669.341008 | 669.3409 | 0.0001 | LPI O-25:7 | C34H55O11P | [M-H]- |
| 770.391629 | 770.4039 | 0.0122 | PS 36:10 | C42H62NO10P | [M-H]- |
| 823.571602 | 823.5706 | 0.001 | PI O-34:0 | C43H85O12P | [M-H]- |
| 742.475941 | 742.4817 | 0.0058 | PE O-38:9 | C43H70NO7P | [M-H]- |
| 819.520734 | 819.5182 | 0.0026 | PG 40:7 | C46H77O10P | [M-H]- |
| 474.261485 | 474.2626 | 0.0011 | LPE 18:3 | C23H42NO7P | [M-H]- |
| 303.232574 | 303.233 | 0.0004 | FA 20:4 | C20H32O2 | [M-H]- |
| 803.538004 | 803.5444 | 0.0064 | LPI O-34:3 | C43H81O11P | [M-H]- |
| 824.519807 | 824.5236 | 0.0038 | PE 43:10 | C48H76NO8P | [M-H]- |
| 755.462942 | 755.4657 | 0.0028 | PA 41:9 | C44H69O8P | [M-H]- |
| 620.33775 | 620.3358 | 0.002 | PE 28:7 | C33H52NO8P | [M-H]- |
| 560.319023 | 560.3358 | 0.0168 | PE 23:2 | C28H52NO8P | [M-H]- |
| 662.34605 | 662.3464 | 0.0003 | PS O-29:8 | C35H54NO9P | [M-H]- |
| 475.245483 | 475.2466 | 0.0012 | PA 20:2 | C23H41O8P | [M-H]- |
| 809.378848 | 809.4247 | 0.0458 | PI 33:7 | C42H67O13P | [M-H]- |
| 801.567082 | 801.5651 | 0.002 | PG 38:2 | C44H83O10P | [M-H]- |
| 828.477794 | 828.4821 | 0.0043 | PS 40:9 | C46H72NO10P | [M-H]- |
| 672.364994 | 672.3671 | 0.0021 | PE 32:9 | C37H56NO8P | [M-H]- |
| 482.366668 | 482.4004 | 0.0337 | NAE 30:6 | C32H53NO2 | [M-H]- |
| 1061.64157 | 1061.6101 | 0.0315 | PIP 45:5 | C54H96O16P2 | [M-H]- |
| 533.270202 | 533.2521 | 0.0181 | BMP 19:3 | C25H43O10P | [M-H]- |
| 692.352196 | 692.3569 | 0.0047 | PS 30:7 | C36H56NO10P | [M-H]- |
| 712.468486 | 712.4794 | 0.0109 | DGCC 32:7 | C42H67NO8 | [M-H]- |
| 706.369956 | 706.3726 | 0.0026 | PS 31:7 | C37H58NO10P | [M-H]- |
| 450.250053 | 450.2626 | 0.0126 | LPE 16:1 | C21H42NO7P | [M-H]- |
| 804.584079 | 804.5784 | 0.0057 | DGTS 40:9 | C50H79NO7 | [M-H]- |
| 674.38085 | 674.3827 | 0.0019 | PE 32:8 | C37H58NO8P | [M-H]- |
| 1153.68705 | 1153.6727 | 0.0144 | PIP 52:8 | C61H104O16P2 | [M-H]- |
| 789.510404 | 789.5076 | 0.0028 | PG O-39:8 | C45H75O9P | [M-H]- |
| 516.308075 | 516.3096 | 0.0015 | PE O-21:3 | C26H48NO7P | [M-H]- |
| 905.473386 | 905.5186 | 0.0452 | PI 40:8 | C49H79O13P | [M-H]- |
| 572.262361 | 572.263 | 0.0007 | PS 21:4 | C27H44NO10P | [M-H]- |
| 739.470246 | 739.4708 | 0.0006 | PA O-41:10 | C44H69O7P | [M-H]- |
| 1070.74891 | 1070.7795 | 0.0306 | PS 57:7 | C63H110NO10P | [M-H]- |

| | | | | | |
|---|---|---|---|---|---|
| 603.348013 | 603.3456 | 0.0024 | PA O-31:8 | C34H53O7P | [M-H]- |
| 791.527312 | 791.5256 | 0.0017 | TG 49:14 | C52H72O6 | [M-H]- |
| 850.527955 | 850.5392 | 0.0113 | PE 45:11 | C50H78NO8P | [M-H]- |
| 754.477568 | 754.4817 | 0.0042 | PE O-39:10 | C44H70NO7P | [M-H]- |
| 763.534369 | 763.5366 | 0.0022 | MGDG 35:4 | C44H76O10 | [M-H]- |
| 821.494569 | 821.5127 | 0.0181 | PA 46:11 | C49H75O8P | [M-H]- |
| 421.235292 | 421.2361 | 0.0008 | LPA 17:1 | C20H39O7P | [M-H]- |
| 530.250777 | 530.2525 | 0.0017 | LPS 19:4 | C25H42NO9P | [M-H]- |
| 741.471465 | 741.4712 | 0.0002 | PG 34:4 | C40H71O10P | [M-H]- |
| 622.317957 | 622.3151 | 0.0029 | PS O-26:7 | C32H50NO9P | [M-H]- |
| 825.566698 | 825.5651 | 0.0016 | PG 40:4 | C46H83O10P | [M-H]- |
| 733.539954 | 733.5389 | 0.0011 | PG O-34:1 | C40H79O9P | [M-H]- |
| 777.397301 | 777.3984 | 0.0011 | PI O-32:9 | C41H63O12P | [M-H]- |
| 854.496937 | 854.4978 | 0.0008 | PS 42:10 | C48H74NO10P | [M-H]- |
| 800.583316 | 800.5811 | 0.0022 | PS O-38:2 | C44H84NO9P | [M-H]- |
| 636.330221 | 636.3307 | 0.0005 | PS O-27:7 | C33H52NO9P | [M-H]- |
| 743.375101 | 743.3777 | 0.0026 | PI 28:5 | C37H61O13P | [M-H]- |
| 1090.73495 | 1090.7482 | 0.0132 | PS 59:11 | C65H106NO10P | [M-H]- |
| 1119.65778 | 1119.6883 | 0.0306 | PIP 49:4 | C58H106O16P2 | [M-H]- |
| 685.335711 | 685.3358 | 0.0001 | PI O-25:6 | C34H55O12P | [M-H]- |
| 770.472733 | 770.4766 | 0.0039 | PE 39:9 | C44H70NO8P | [M-H]- |
| 819.616412 | 819.6121 | 0.0043 | PG 39:0 | C45H89O10P | [M-H]- |
| 395.245591 | 395.2568 | 0.0112 | LPA O-16:0 | C19H41O6P | [M-H]- |
| 660.350449 | 660.3671 | 0.0166 | PE 31:8 | C36H56NO8P | [M-H]- |
| 646.348121 | 646.3514 | 0.0033 | PE 30:8 | C35H54NO8P | [M-H]- |
| 652.325516 | 652.3256 | 0.0001 | PS 27:6 | C33H52NO10P | [M-H]- |
| 810.467251 | 810.4716 | 0.0043 | PS O-40:11 | C46H70NO9P | [M-H]- |
| 848.507892 | 848.5236 | 0.0157 | PE 45:12 | C50H76NO8P | [M-H]- |
| 1084.76456 | 1084.7951 | 0.0306 | PS 58:7 | C64H112NO10P | [M-H]- |
| 632.339192 | 632.3358 | 0.0034 | PE 29:8 | C34H52NO8P | [M-H]- |
| 790.590617 | 790.5968 | 0.0061 | PS O-37:0 | C43H86NO9P | [M-H]- |
| 1167.62542 | 1167.5921 | 0.0333 | PIP2 47:6 | C56H99O19P3 | [M-H]- |
| 510.290408 | 510.2838 | 0.0067 | LPS 17:0 | C23H46NO9P | [M-H]- |
| 763.496394 | 763.492 | 0.0044 | PG O-37:7 | C43H73O9P | [M-H]- |
| 937.562593 | 937.5812 | 0.0186 | PI 42:6 | C51H87O13P | [M-H]- |
| 481.256327 | 481.2572 | 0.0009 | LPG 16:1 | C22H43O9P | [M-H]- |
| 690.376704 | 690.3777 | 0.0009 | PS O-31:8 | C37H58NO9P | [M-H]- |
| 1019.66046 | 1019.6594 | 0.0011 | PI 48:7 | C57H97O13P | [M-H]- |
| 518.28734 | 518.2888 | 0.0015 | PE 20:2 | C25H46NO8P | [M-H]- |
| 460.254002 | 460.247 | 0.007 | LPE 17:3 | C22H40NO7P | [M-H]- |
| 1033.5862 | 1033.5788 | 0.0074 | PIP 43:5 | C52H92O16P2 | [M-H]- |
| 901.434306 | 901.391 | 0.0433 | PIP 34:8 | C43H68O16P2 | [M-H]- |
| 809.491939 | 809.5127 | 0.0207 | PA 45:10 | C48H75O8P | [M-H]- |
| 1005.6452 | 1005.6438 | 0.0014 | PI 47:7 | C56H95O13P | [M-H]- |
| 680.356389 | 680.3569 | 0.0005 | PS 29:6 | C35H56NO10P | [M-H]- |

| | | | | | |
|---|---|---|---|---|---|
| 790.571023 | 790.5756 | 0.0046 | PE O-41:6 | C46H82NO7P | [M-H]- |
| 849.601025 | 849.6015 | 0.0005 | PG O-43:6 | C49H87O9P | [M-H]- |
| 892.506475 | 892.5134 | 0.0069 | PS 45:12 | C51H76NO10P | [M-H]- |
| 927.562149 | 927.5393 | 0.0229 | PI O-43:11 | C52H81O12P | [M-H]- |
| 761.36223 | 761.3284 | 0.0338 | PIP 23:1 | C32H60O16P2 | [M-H]- |
| 719.513978 | 719.5233 | 0.0093 | PG O-33:1 | C39H77O9P | [M-H]- |
| 507.256613 | 507.2729 | 0.0162 | LPG 18:2 | C24H45O9P | [M-H]- |
| 734.478088 | 734.4766 | 0.0015 | PE 36:6 | C41H70NO8P | [M-H]- |
| 1117.64172 | 1117.6727 | 0.031 | PIP 49:5 | C58H104O16P2 | [M-H]- |
| 945.469153 | 945.4536 | 0.0156 | PIP 37:7 | C46H76O16P2 | [M-H]- |
| 776.582916 | 776.5811 | 0.0018 | PS O-36:0 | C42H84NO9P | [M-H]- |
| 1127.66031 | 1127.657 | 0.0033 | PIP 50:7 | C59H102O16P2 | [M-H]- |
| 1101.61001 | 1101.6414 | 0.0314 | PIP 48:6 | C57H100O16P2 | [M-H]- |
| 894.516593 | 894.5291 | 0.0125 | PS 45:11 | C51H78NO10P | [M-H]- |
| 883.525635 | 883.5342 | 0.0086 | PI 38:5 | C47H81O13P | [M-H]- |
| 876.505642 | 876.5549 | 0.0492 | PE 47:12 | C52H80NO8P | [M-H]- |
| 889.481516 | 889.4849 | 0.0034 | PIP 32:0 | C41H80O16P2 | [M-H]- |
| 764.481702 | 764.4872 | 0.0055 | PS O-36:6 | C42H72NO9P | [M-H]- |
| 532.339251 | 532.3409 | 0.0016 | PE O-22:2 | C27H52NO7P | [M-H]- |
| 861.483398 | 861.4828 | 0.0006 | SQDG 38:8 | C47H74O12S | [M-H]- |
| 880.500073 | 880.5134 | 0.0133 | PS 44:11 | C50H76NO10P | [M-H]- |
| 973.532092 | 973.5788 | 0.0467 | PIP 38:0 | C47H92O16P2 | [M-H]- |
| 729.239242 | 729.2658 | 0.0266 | PIP 21:3 | C30H52O16P2 | [M-H]- |
| 1151.63797 | 1151.6547 | 0.0167 | PIP2 45:0 | C54H107O19P3 | [M-H]- |
| 1145.671 | 1145.704 | 0.033 | PIP 51:5 | C60H108O16P2 | [M-H]- |
| 1003.4102 | 1003.4356 | 0.0254 | PIP2 35:4 | C44H79O19P3 | [M-H]- |
| 898.520872 | 898.5604 | 0.0395 | PS 45:9 | C51H82NO10P | [M-H]- |

*Supporting Table 8: List of annotations for signals identified as statistically different (ANOVA multiple-sample test, p-value ≤ 0.01) between the OX1, B8, C12 and E2 cell line (biological triplicates, each consisting of 22,500 individual mass spectra of multiple individual cells) in LPS-stimulated state in negative-ion mode from LIPID MAPS database research with the lowest mass difference for each input mass.*

| Input mass | Matched mass | Δ $m/z$ | Name | Formula | Ion |
|---|---|---|---|---|---|
| 807.277222 | 807.3127 | 0.0355 | PIP 27:6 | C36H58O16P2 | [M-H]- |
| 821.567757 | 821.5702 | 0.0024 | PG O-41:6 | C47H83O9P | [M-H]- |
| 809.274345 | 809.2321 | 0.0422 | PIP2 21:3 | C30H53O19P3 | [M-H]- |
| 847.584831 | 847.5859 | 0.001 | PG O-43:7 | C49H85O9P | [M-H]- |
| 822.56969 | 822.5655 | 0.0042 | PS O-40:5 | C46H82NO9P | [M-H]- |
| 819.552138 | 819.5546 | 0.0024 | PG O-41:7 | C47H81O9P | [M-H]- |
| 678.493534 | 678.495 | 0.0015 | DGCC 29:3 | C39H69NO8 | [M-H]- |
| 747.566407 | 747.5569 | 0.0095 | TG 45:8 | C48H76O6 | [M-H]- |
| 678.476949 | 678.4716 | 0.0054 | PS O-29:0 | C35H70NO9P | [M-H]- |
| 804.577797 | 804.5784 | 0.0006 | DGTS 40:9 | C50H79NO7 | [M-H]- |
| 1087.71523 | 1087.722 | 0.0068 | PI 53:8 | C62H105O13P | [M-H]- |
| 805.574527 | 805.5753 | 0.0008 | PA 44:5 | C47H83O8P | [M-H]- |

| | | | | | |
|---|---|---|---|---|---|
| 848.587898 | 848.5811 | 0.0068 | PS O-42:6 | C48H84NO9P | [M-H]- |
| 1145.72845 | 1145.704 | 0.0245 | PIP 51:5 | C60H108O16P2 | [M-H]- |
| 864.450836 | 864.4821 | 0.0313 | PS 43:12 | C49H72NO10P | [M-H]- |
| 830.592946 | 830.594 | 0.0011 | DGTS 42:10 | C52H81NO7 | [M-H]- |
| 1088.72022 | 1088.7325 | 0.0123 | PS 59:12 | C65H104NO10P | [M-H]- |
| 749.583888 | 749.5726 | 0.0113 | TG 45:7 | C48H78O6 | [M-H]- |
| 1175.70531 | 1175.7509 | 0.0456 | PIP 53:4 | C62H114O16P2 | [M-H]- |
| 748.567449 | 748.5733 | 0.0058 | DGCC 34:3 | C44H79NO8 | [M-H]- |
| 1089.73098 | 1089.7377 | 0.0067 | PI 53:7 | C62H107O13P | [M-H]- |
| 628.448894 | 628.4348 | 0.0141 | PE O-29:3 | C34H64NO7P | [M-H]- |
| 845.569854 | 845.5702 | 0.0003 | PG O-43:8 | C49H83O9P | [M-H]- |
| 791.594019 | 791.596 | 0.002 | PA O-44:5 | C47H85O7P | [M-H]- |
| 728.380316 | 728.4297 | 0.0494 | PE 36:9 | C41H64NO8P | [M-H]- |
| 780.569237 | 780.5784 | 0.0091 | DGTS 38:7 | C48H79NO7 | [M-H]- |
| 1143.71426 | 1143.6883 | 0.0259 | PIP 51:6 | C60H106O16P2 | [M-H]- |
| 465.288521 | 465.2987 | 0.0102 | PA O-20:0 | C23H47O7P | [M-H]- |
| 1073.6574 | 1073.6101 | 0.0473 | PIP 46:6 | C55H96O16P2 | [M-H]- |
| 779.559007 | 779.5596 | 0.0006 | PA 42:4 | C45H81O8P | [M-H]- |
| 823.571602 | 823.5706 | 0.001 | PI O-34:0 | C43H85O12P | [M-H]- |
| 831.590637 | 831.5909 | 0.0003 | PA 46:6 | C49H85O8P | [M-H]- |
| 482.366668 | 482.4004 | 0.0337 | NAE 30:6 | C32H53NO2 | [M-H]- |
| 804.584079 | 804.5784 | 0.0057 | DGTS 40:9 | C50H79NO7 | [M-H]- |
| 821.494569 | 821.5127 | 0.0181 | PA 46:11 | C49H75O8P | [M-H]- |
| 557.321303 | 557.3249 | 0.0036 | PA 26:3 | C29H51O8P | [M-H]- |
| 309.278808 | 309.2799 | 0.0011 | FA 20:1 | C20H38O2 | [M-H]- |
| 865.454922 | 865.4873 | 0.0323 | PI 37:7 | C46H75O13P | [M-H]- |
| 857.678202 | 857.6665 | 0.0117 | TG 53:9 | C56H90O6 | [M-H]- |
| 690.376704 | 690.3777 | 0.0009 | PS O-31:8 | C37H58NO9P | [M-H]- |
| 558.264701 | 558.2474 | 0.0173 | PS 20:4 | C26H42NO10P | [M-H]- |
| 782.473837 | 782.4766 | 0.0028 | PE 40:10 | C45H70NO8P | [M-H]- |
| 1171.74422 | 1171.7196 | 0.0246 | PIP 53:6 | C62H110O16P2 | [M-H]- |
| 849.601025 | 849.6015 | 0.0005 | PG O-43:6 | C49H87O9P | [M-H]- |
| 683.285407 | 683.3107 | 0.0253 | SQDG 25:6 | C34H52O12S | [M-H]- |
| 1003.4102 | 1003.4356 | 0.0254 | PIP2 35:4 | C44H79O19P3 | [M-H]- |

*Supporting Table 9: List of annotations for signals of the OX1 cell line identified as statistically different (ANOVA multiple-sample test, p-value ≤ 0.01) between the naïve and LPS-stimulated state (biological triplicates, each consisting of 22,500 individual mass spectra of multiple individual cells) in positive-ion mode from LIPID MAPS database research with the lowest mass difference for each input mass.*

| Input mass | Matched mass | Δ *m/z* | Name | Formula | Ion |
|---|---|---|---|---|---|
| 478.29285 | 478.2939 | 0.0011 | LPE 18:1 | C23H46NO7P | [M-H]- |
| 500.277063 | 500.2783 | 0.0012 | PE O-20:4 | C25H44NO7P | [M-H]- |
| 1085.65202 | 1085.6101 | 0.0419 | PIP 47:7 | C56H96O16P2 | [M-H]- |
| 711.337808 | 711.342 | 0.0042 | SQDG 27:6 | C36H56O12S | [M-H]- |
| 502.292683 | 502.2939 | 0.0012 | PE O-20:3 | C25H46NO7P | [M-H]- |

| | | | | | |
|---|---|---|---|---|---|
| 526.292467 | 526.2939 | 0.0015 | PE O-22:5 | C27H46NO7P | [M-H]- |
| 498.261457 | 498.2626 | 0.0012 | PE O-20:5 | C25H42NO7P | [M-H]- |
| 530.323728 | 530.3252 | 0.0015 | PE O-22:3 | C27H50NO7P | [M-H]- |
| 847.584831 | 847.5859 | 0.001 | PG O-43:7 | C49H85O9P | [M-H]- |
| 476.277179 | 476.2783 | 0.0011 | LPE 18:2 | C23H44NO7P | [M-H]- |
| 501.280483 | 501.2987 | 0.0182 | PA O-23:3 | C26H47O7P | [M-H]- |
| 1059.60218 | 1059.5944 | 0.0077 | PIP 45:6 | C54H94O16P2 | [M-H]- |
| 450.261738 | 450.2626 | 0.0009 | LPE 16:1 | C21H42NO7P | [M-H]- |
| 822.56969 | 822.5655 | 0.0042 | PS O-40:5 | C46H82NO9P | [M-H]- |
| 819.552138 | 819.5546 | 0.0024 | PG O-41:7 | C47H81O9P | [M-H]- |
| 479.296277 | 479.3143 | 0.018 | PA O-21:0 | C24H49O7P | [M-H]- |
| 528.308099 | 528.3096 | 0.0015 | PE O-22:4 | C27H48NO7P | [M-H]- |
| 459.250772 | 459.2517 | 0.001 | PA O-20:3 | C23H41O7P | [M-H]- |
| 504.308372 | 504.3096 | 0.0012 | PE O-20:2 | C25H48NO7P | [M-H]- |
| 798.53892 | 798.5443 | 0.0054 | PE O-42:9 | C47H78NO7P | [M-H]- |
| 451.266043 | 451.283 | 0.017 | LPA 19:0 | C22H45O7P | [M-H]- |
| 1087.71523 | 1087.722 | 0.0068 | PI 53:8 | C62H105O13P | [M-H]- |
| 848.587898 | 848.5811 | 0.0068 | PS O-42:6 | C48H84NO9P | [M-H]- |
| 483.250551 | 483.2517 | 0.0012 | PA O-22:5 | C25H41O7P | [M-H]- |
| 527.29592 | 527.2779 | 0.018 | PA 24:4 | C27H45O8P | [M-H]- |
| 499.264829 | 499.283 | 0.0182 | PA O-23:4 | C26H45O7P | [M-H]- |
| 449.266413 | 449.2674 | 0.001 | LPA 19:1 | C22H43O7P | [M-H]- |
| 1088.72022 | 1088.7325 | 0.0123 | PS 59:12 | C65H104NO10P | [M-H]- |
| 528.295274 | 528.3096 | 0.0143 | PE O-22:4 | C27H48NO7P | [M-H]- |
| 660.497379 | 660.4974 | 0 | PE O-31:1 | C36H72NO7P | [M-H]- |
| 524.277041 | 524.2783 | 0.0012 | PE O-22:6 | C27H44NO7P | [M-H]- |
| 421.212774 | 421.1997 | 0.0131 | LPG O-13:3 | C19H35O8P | [M-H]- |
| 680.46119 | 680.4661 | 0.0049 | PE O-33:5 | C38H68NO7P | [M-H]- |
| 542.287151 | 542.2888 | 0.0017 | PE 22:4 | C27H46NO8P | [M-H]- |
| 516.271819 | 516.2732 | 0.0014 | PE 20:3 | C25H44NO8P | [M-H]- |
| 544.30282 | 544.3045 | 0.0017 | PE 22:3 | C27H48NO8P | [M-H]- |
| 1179.65084 | 1179.686 | 0.0351 | PIP2 47:0 | C56H111O19P3 | [M-H]- |
| 845.569854 | 845.5702 | 0.0003 | PG O-43:8 | C49H83O9P | [M-H]- |
| 1131.65655 | 1131.6883 | 0.0318 | PIP 50:5 | C59H106O16P2 | [M-H]- |
| 1113.67609 | 1113.6414 | 0.0347 | PIP 49:7 | C58H100O16P2 | [M-H]- |
| 477.280806 | 477.2987 | 0.0179 | PA O-21:1 | C24H47O7P | [M-H]- |
| 520.302949 | 520.3045 | 0.0015 | PE 20:1 | C25H48NO8P | [M-H]- |
| 485.266194 | 485.2674 | 0.0012 | PA O-22:4 | C25H43O7P | [M-H]- |
| 407.206636 | 407.2204 | 0.0138 | LPA 16:1 | C19H37O7P | [M-H]- |
| 823.571602 | 823.5706 | 0.001 | PI O-34:0 | C43H85O12P | [M-H]- |
| 831.590637 | 831.5909 | 0.0003 | PA 46:6 | C49H85O8P | [M-H]- |
| 533.270202 | 533.2521 | 0.0181 | BMP 19:3 | C25H43O10P | [M-H]- |
| 1171.67361 | 1171.7196 | 0.046 | PIP 53:6 | C62H110O16P2 | [M-H]- |
| 777.397301 | 777.3984 | 0.0011 | PI O-32:9 | C41H63O12P | [M-H]- |
| 1119.65778 | 1119.6883 | 0.0306 | PIP 49:4 | C58H106O16P2 | [M-H]- |

| | | | | | |
|---|---|---|---|---|---|
| 652.325516 | 652.3256 | 0.0001 | PS 27:6 | C33H52NO10P | [M-H]- |
| 309.278808 | 309.2799 | 0.0011 | FA 20:1 | C20H38O2 | [M-H]- |
| 1161.68916 | 1161.7353 | 0.0461 | PIP 52:4 | C61H112O16P2 | [M-H]- |
| 1019.60524 | 1019.5631 | 0.0421 | PIP 42:5 | C51H90O16P2 | [M-H]- |
| 510.290408 | 510.2838 | 0.0067 | LPS 17:0 | C23H46NO9P | [M-H]- |
| 481.256327 | 481.2572 | 0.0009 | LPG 16:1 | C22H43O9P | [M-H]- |
| 518.28734 | 518.2888 | 0.0015 | PE 20:2 | C25H46NO8P | [M-H]- |
| 1043.60452 | 1043.5631 | 0.0414 | PIP 44:7 | C53H90O16P2 | [M-H]- |
| 1109.73378 | 1109.7332 | 0.0006 | SQDG 56:10 | C65H106O12S | [M-H]- |
| 790.571023 | 790.5756 | 0.0046 | PE O-41:6 | C46H82NO7P | [M-H]- |
| 601.296036 | 601.3147 | 0.0187 | PG 24:4 | C30H51O10P | [M-H]- |
| 1117.64172 | 1117.6727 | 0.031 | PIP 49:5 | C58H104O16P2 | [M-H]- |
| 1157.67328 | 1157.704 | 0.0307 | PIP 52:6 | C61H108O16P2 | [M-H]- |
| 1127.66031 | 1127.657 | 0.0033 | PIP 50:7 | C59H102O16P2 | [M-H]- |
| 820.523443 | 820.5287 | 0.0052 | PE O-44:12 | C49H76NO7P | [M-H]- |
| 1133.72057 | 1133.7146 | 0.006 | DGDG 51:12 | C66H102O15 | [M-H]- |

*Supporting Table 10: List of annotations for signals of the B8 cell line identified as statistically different (ANOVA multiple-sample test, p-value ≤ 0.01) between the naïve and LPS-stimulated state (biological triplicates, each consisting of 22,500 individual mass spectra of multiple individual cells) in positive-ion mode from LIPID MAPS database research with the lowest mass difference for each input mass.*

| Input mass | Matched mass | Δ *m/z* | Name | Formula | Ion |
|---|---|---|---|---|---|
| 807.277222 | 807.3127 | 0.0355 | PIP 27:6 | C36H58O16P2 | [M-H]- |
| 579.230197 | 579.2576 | 0.0274 | LPI 17:3 | C26H45O12P | [M-H]- |
| 753.385492 | 753.3889 | 0.0034 | SQDG 30:6 | C39H62O12S | [M-H]- |
| 711.337808 | 711.342 | 0.0042 | SQDG 27:6 | C36H56O12S | [M-H]- |
| 821.567757 | 821.5702 | 0.0024 | PG O-41:6 | C47H83O9P | [M-H]- |
| 701.490014 | 701.4787 | 0.0113 | TG 42:10 | C45H66O6 | [M-H]- |
| 861.51819 | 861.544 | 0.0258 | PA 49:12 | C52H79O8P | [M-H]- |
| 713.330091 | 713.3308 | 0.0007 | PI 26:6 | C35H55O13P | [M-H]- |
| 697.336284 | 697.3358 | 0.0004 | PI O-26:7 | C35H55O12P | [M-H]- |
| 862.52345 | 862.5392 | 0.0158 | PE 46:12 | C51H78NO8P | [M-H]- |
| 799.53054 | 799.5283 | 0.0022 | PA 44:8 | C47H77O8P | [M-H]- |
| 789.521268 | 789.5287 | 0.0075 | LPI O-33:3 | C42H79O11P | [M-H]- |
| 687.523362 | 687.5334 | 0.0101 | PA O-36:1 | C39H77O7P | [M-H]- |
| 760.489201 | 760.4923 | 0.0031 | PE 38:7 | C43H72NO8P | [M-H]- |
| 767.401176 | 767.4046 | 0.0034 | SQDG 31:6 | C40H64O12S | [M-H]- |
| 1151.6724 | 1151.657 | 0.0154 | PIP 52:9 | C61H102O16P2 | [M-H]- |
| 689.537871 | 689.5491 | 0.0112 | PA O-36:0 | C39H79O7P | [M-H]- |
| 753.357686 | 753.3621 | 0.0044 | PI 29:7 | C38H59O13P | [M-H]- |
| 788.516412 | 788.5236 | 0.0072 | PE 40:7 | C45H76NO8P | [M-H]- |
| 863.53501 | 863.5596 | 0.0246 | PA 49:11 | C52H81O8P | [M-H]- |
| 809.274345 | 809.2321 | 0.0422 | PIP2 21:3 | C30H53O19P3 | [M-H]- |
| 786.503641 | 786.5079 | 0.0043 | PE 40:8 | C45H74NO8P | [M-H]- |
| 1087.63027 | 1087.6257 | 0.0045 | PIP 47:6 | C56H98O16P2 | [M-H]- |

| | | | | | |
|---|---|---|---|---|---|
| 723.350272 | 723.3515 | 0.0012 | PI O-28:8 | C37H57O12P | [M-H]- |
| 917.506439 | 917.5162 | 0.0098 | PIP 34:0 | C43H84O16P2 | [M-H]- |
| 787.505698 | 787.5131 | 0.0074 | LPI O-33:4 | C42H77O11P | [M-H]- |
| 856.539902 | 856.5498 | 0.0099 | PS O-43:9 | C49H80NO9P | [M-H]- |
| 847.584831 | 847.5859 | 0.001 | PG O-43:7 | C49H85O9P | [M-H]- |
| 835.503827 | 835.5283 | 0.0245 | PA 47:11 | C50H77O8P | [M-H]- |
| 688.528624 | 688.5287 | 0 | PE O-33:1 | C38H76NO7P | [M-H]- |
| 648.388403 | 648.3882 | 0.0002 | PS 26:1 | C32H60NO10P | [M-H]- |
| 725.36757 | 725.3671 | 0.0004 | PI O-28:7 | C37H59O12P | [M-H]- |
| 801.54689 | 801.544 | 0.0029 | PA 44:7 | C47H79O8P | [M-H]- |
| 631.362047 | 631.3617 | 0.0004 | PG 26:3 | C32H57O10P | [M-H]- |
| 797.632327 | 797.643 | 0.0106 | PA O-44:2 | C47H91O7P | [M-H]- |
| 919.522069 | 919.5342 | 0.0121 | PI 41:8 | C50H81O13P | [M-H]- |
| 619.362214 | 619.3617 | 0.0005 | PG 25:2 | C31H57O10P | [M-H]- |
| 815.557313 | 815.5596 | 0.0023 | PA 45:7 | C48H81O8P | [M-H]- |
| 822.56969 | 822.5655 | 0.0042 | PS O-40:5 | C46H82NO9P | [M-H]- |
| 901.515646 | 901.5141 | 0.0015 | SQDG 41:9 | C50H78O12S | [M-H]- |
| 690.545536 | 690.5443 | 0.0012 | PE O-33:0 | C38H78NO7P | [M-H]- |
| 912.59865 | 912.6124 | 0.0138 | PS O-47:9 | C53H88NO9P | [M-H]- |
| 781.416745 | 781.4202 | 0.0035 | SQDG 32:6 | C41H66O12S | [M-H]- |
| 741.358124 | 741.3621 | 0.0039 | PI 28:6 | C37H59O13P | [M-H]- |
| 664.383141 | 664.3984 | 0.0152 | PE 31:6 | C36H60NO8P | [M-H]- |
| 856.599328 | 856.6073 | 0.008 | PS 41:2 | C47H88NO10P | [M-H]- |
| 799.647719 | 799.6586 | 0.0109 | PA O-44:1 | C47H93O7P | [M-H]- |
| 747.566407 | 747.5569 | 0.0095 | TG 45:8 | C48H76O6 | [M-H]- |
| 857.542954 | 857.5454 | 0.0025 | SQDG 37:3 | C46H82O12S | [M-H]- |
| 903.526785 | 903.5298 | 0.003 | SQDG 41:8 | C50H80O12S | [M-H]- |
| 909.640069 | 909.6379 | 0.0022 | PA 52:9 | C55H91O8P | [M-H]- |
| 649.372428 | 649.3722 | 0.0002 | LPI O-23:3 | C32H59O11P | [M-H]- |
| 882.555202 | 882.5655 | 0.0103 | PS O-45:10 | C51H82NO9P | [M-H]- |
| 915.494031 | 915.5005 | 0.0065 | PIP 34:1 | C43H82O16P2 | [M-H]- |
| 563.318058 | 563.3143 | 0.0037 | PA O-28:7 | C31H49O7P | [M-H]- |
| 739.346157 | 739.3464 | 0.0003 | PI 28:7 | C37H57O13P | [M-H]- |
| 911.656069 | 911.6535 | 0.0025 | PA 52:8 | C55H93O8P | [M-H]- |
| 763.378227 | 763.3733 | 0.0049 | SQDG 31:8 | C40H60O12S | [M-H]- |
| 376.330091 | 376.3221 | 0.008 | NAE 22:3 | C24H43NO2 | [M-H]- |
| 1053.5564 | 1053.5475 | 0.0089 | PIP 45:9 | C54H88O16P2 | [M-H]- |
| 804.577797 | 804.5784 | 0.0006 | DGTS 40:9 | C50H79NO7 | [M-H]- |
| 589.334754 | 589.33 | 0.0048 | PA O-30:8 | C33H51O7P | [M-H]- |
| 899.504094 | 899.508 | 0.0039 | PI O-41:11 | C50H77O12P | [M-H]- |
| 920.526963 | 920.5447 | 0.0178 | PS 47:12 | C53H80NO10P | [M-H]- |
| 473.281529 | 473.2674 | 0.0142 | PA O-21:3 | C24H43O7P | [M-H]- |
| 1167.66453 | 1167.6883 | 0.0238 | PIP 53:8 | C62H106O16P2 | [M-H]- |
| 802.530082 | 802.5263 | 0.0037 | DGCC 39:11 | C49H73NO8 | [M-H]- |
| 805.574527 | 805.5753 | 0.0008 | PA 44:5 | C47H83O8P | [M-H]- |

| | | | | | |
|---|---|---|---|---|---|
| 848.587898 | 848.5811 | 0.0068 | PS O-42:6 | C48H84NO9P | [M-H]- |
| 814.497427 | 814.5029 | 0.0054 | PS O-40:9 | C46H74NO9P | [M-H]- |
| 648.363632 | 648.3671 | 0.0035 | PE 30:7 | C35H56NO8P | [M-H]- |
| 809.292836 | 809.3284 | 0.0356 | PIP 27:5 | C36H60O16P2 | [M-H]- |
| 1153.72482 | 1153.7666 | 0.0418 | PIP 51:1 | C60H116O16P2 | [M-H]- |
| 883.558011 | 883.5611 | 0.0031 | SQDG 39:4 | C48H84O12S | [M-H]- |
| 683.318669 | 683.3202 | 0.0015 | PI O-25:7 | C34H53O12P | [M-H]- |
| 535.285835 | 535.283 | 0.0028 | PA O-26:7 | C29H45O7P | [M-H]- |
| 1145.72845 | 1145.704 | 0.0245 | PIP 51:5 | C60H108O16P2 | [M-H]- |
| 1089.6385 | 1089.6414 | 0.0029 | PIP 47:5 | C56H100O16P2 | [M-H]- |
| 641.386386 | 641.3824 | 0.004 | PG O-28:5 | C34H59O9P | [M-H]- |
| 841.284369 | 841.2947 | 0.0104 | PIP2 23:1 | C32H61O19P3 | [M-H]- |
| 726.372779 | 726.414 | 0.0413 | PE 36:10 | C41H62NO8P | [M-H]- |
| 781.388694 | 781.3934 | 0.0047 | PI 31:7 | C40H63O13P | [M-H]- |
| 737.366421 | 737.3671 | 0.0007 | PI O-29:8 | C38H59O12P | [M-H]- |
| 779.373133 | 779.3777 | 0.0046 | PI 31:8 | C40H61O13P | [M-H]- |
| 577.334959 | 577.33 | 0.005 | PA O-29:7 | C32H51O7P | [M-H]- |
| 910.644574 | 910.6543 | 0.0097 | PS 45:3 | C51H94NO10P | [M-H]- |
| 617.359792 | 617.3613 | 0.0015 | PA O-32:8 | C35H55O7P | [M-H]- |
| 1003.36126 | 1003.3393 | 0.0219 | PIP3 29:2 | C38H72O22P4 | [M-H]- |
| 680.37938 | 680.3933 | 0.0139 | PS O-30:6 | C36H60NO9P | [M-H]- |
| 869.278409 | 869.326 | 0.0476 | PIP2 25:1 | C34H65O19P3 | [M-H]- |
| 749.583888 | 749.5726 | 0.0113 | TG 45:7 | C48H78O6 | [M-H]- |
| 1175.70531 | 1175.7509 | 0.0456 | PIP 53:4 | C62H114O16P2 | [M-H]- |
| 807.478447 | 807.497 | 0.0186 | PA 45:11 | C48H73O8P | [M-H]- |
| 751.386442 | 751.3828 | 0.0036 | PI O-30:8 | C39H61O12P | [M-H]- |
| 748.567449 | 748.5733 | 0.0058 | DGCC 34:3 | C44H79NO8 | [M-H]- |
| 925.664675 | 925.6692 | 0.0045 | PA 53:8 | C56H95O8P | [M-H]- |
| 667.401804 | 667.3981 | 0.0038 | PG O-30:6 | C36H61O9P | [M-H]- |
| 856.549001 | 856.5498 | 0.0008 | PS O-43:9 | C49H80NO9P | [M-H]- |
| 809.286209 | 809.3284 | 0.0422 | PIP 27:5 | C36H60O16P2 | [M-H]- |
| 775.505878 | 775.5131 | 0.0072 | LPI O-32:3 | C41H77O11P | [M-H]- |
| 912.660993 | 912.6699 | 0.0089 | PS 45:2 | C51H96NO10P | [M-H]- |
| 784.522331 | 784.5287 | 0.0063 | PE O-41:9 | C46H76NO7P | [M-H]- |
| 685.298856 | 685.2995 | 0.0006 | PI 24:6 | C33H51O13P | [M-H]- |
| 800.514436 | 800.5236 | 0.0092 | PE 41:8 | C46H76NO8P | [M-H]- |
| 891.492407 | 891.5029 | 0.0105 | PI 39:8 | C48H77O13P | [M-H]- |
| 660.363338 | 660.3671 | 0.0037 | PE 31:8 | C36H56NO8P | [M-H]- |
| 812.481066 | 812.4872 | 0.0061 | PS O-40:10 | C46H72NO9P | [M-H]- |
| 709.334222 | 709.3263 | 0.0079 | SQDG 27:7 | C36H54O12S | [M-H]- |
| 845.569854 | 845.5702 | 0.0003 | PG O-43:8 | C49H83O9P | [M-H]- |
| 791.594019 | 791.596 | 0.002 | PA O-44:5 | C47H85O7P | [M-H]- |
| 787.494047 | 787.492 | 0.0021 | PG O-39:9 | C45H73O9P | [M-H]- |
| 549.301874 | 549.2987 | 0.0032 | PA O-27:7 | C30H47O7P | [M-H]- |
| 1143.71426 | 1143.6883 | 0.0259 | PIP 51:6 | C60H106O16P2 | [M-H]- |

| | | | | | |
|---|---|---|---|---|---|
| 676.397485 | 676.3984 | 0.0009 | PE 32:7 | C37H60NO8P | [M-H]- |
| 711.353035 | 711.3515 | 0.0015 | PI O-27:7 | C36H57O12P | [M-H]- |
| 465.288521 | 465.2987 | 0.0102 | PA O-20:0 | C23H47O7P | [M-H]- |
| 1169.67818 | 1169.704 | 0.0258 | PIP 53:7 | C62H108O16P2 | [M-H]- |
| 861.404439 | 861.3597 | 0.0447 | PIP 31:7 | C40H64O16P2 | [M-H]- |
| 840.51202 | 840.5185 | 0.0065 | PS O-42:10 | C48H76NO9P | [M-H]- |
| 755.391091 | 755.386 | 0.0051 | DGDG 23:5 | C38H60O15 | [M-H]- |
| 634.35451 | 634.3514 | 0.0031 | PE 29:7 | C34H54NO8P | [M-H]- |
| 739.385294 | 739.3828 | 0.0025 | PI O-29:7 | C38H61O12P | [M-H]- |
| 1073.6574 | 1073.6101 | 0.0473 | PIP 46:6 | C55H96O16P2 | [M-H]- |
| 535.229747 | 535.2314 | 0.0016 | LPI O-15:4 | C24H41O11P | [M-H]- |
| 632.373371 | 632.3722 | 0.0012 | PE O-30:8 | C35H56NO7P | [M-H]- |
| 732.466019 | 732.461 | 0.005 | PE 36:7 | C41H68NO8P | [M-H]- |
| 1015.34021 | 1015.3393 | 0.0009 | PIP3 30:3 | C39H72O22P4 | [M-H]- |
| 620.393744 | 620.3933 | 0.0004 | PS O-25:1 | C31H60NO9P | [M-H]- |
| 803.538004 | 803.5444 | 0.0064 | LPI O-34:3 | C43H81O11P | [M-H]- |
| 831.590637 | 831.5909 | 0.0003 | PA 46:6 | C49H85O8P | [M-H]- |
| 620.33775 | 620.3358 | 0.002 | PE 28:7 | C33H52NO8P | [M-H]- |
| 828.477794 | 828.4821 | 0.0043 | PS 40:9 | C46H72NO10P | [M-H]- |
| 672.364994 | 672.3671 | 0.0021 | PE 32:9 | C37H56NO8P | [M-H]- |
| 482.366668 | 482.4004 | 0.0337 | NAE 30:6 | C32H53NO2 | [M-H]- |
| 706.369956 | 706.3726 | 0.0026 | PS 31:7 | C37H58NO10P | [M-H]- |
| 804.584079 | 804.5784 | 0.0057 | DGTS 40:9 | C50H79NO7 | [M-H]- |
| 791.527312 | 791.5256 | 0.0017 | TG 49:14 | C52H72O6 | [M-H]- |
| 754.477568 | 754.4817 | 0.0042 | PE O-39:10 | C44H70NO7P | [M-H]- |
| 622.317957 | 622.3151 | 0.0029 | PS O-26:7 | C32H50NO9P | [M-H]- |
| 857.519543 | 857.5186 | 0.001 | PI 36:4 | C45H79O13P | [M-H]- |
| 777.397301 | 777.3984 | 0.0011 | PI O-32:9 | C41H63O12P | [M-H]- |
| 854.496937 | 854.4978 | 0.0008 | PS 42:10 | C48H74NO10P | [M-H]- |
| 685.335711 | 685.3358 | 0.0001 | PI O-25:6 | C34H55O12P | [M-H]- |
| 770.472733 | 770.4766 | 0.0039 | PE 39:9 | C44H70NO8P | [M-H]- |
| 638.331016 | 638.3464 | 0.0153 | PS O-27:6 | C33H54NO9P | [M-H]- |
| 567.257236 | 567.2576 | 0.0004 | LPI 16:2 | C25H45O12P | [M-H]- |
| 652.325516 | 652.3256 | 0.0001 | PS 27:6 | C33H52NO10P | [M-H]- |
| 632.339192 | 632.3358 | 0.0034 | PE 29:8 | C34H52NO8P | [M-H]- |
| 865.454922 | 865.4873 | 0.0323 | PI 37:7 | C46H75O13P | [M-H]- |
| 901.434306 | 901.391 | 0.0433 | PIP 34:8 | C43H68O16P2 | [M-H]- |
| 1171.74422 | 1171.7196 | 0.0246 | PIP 53:6 | C62H110O16P2 | [M-H]- |
| 809.491939 | 809.5127 | 0.0207 | PA 45:10 | C48H75O8P | [M-H]- |
| 478.220065 | 478.2212 | 0.0011 | LPS 15:2 | C21H38NO9P | [M-H]- |
| 849.601025 | 849.6015 | 0.0005 | PG O-43:6 | C49H87O9P | [M-H]- |
| 866.666904 | 866.6644 | 0.0025 | PE 45:3 | C50H94NO8P | [M-H]- |
| 968.722255 | 968.7325 | 0.0103 | PS 49:2 | C55H104NO10P | [M-H]- |
| 719.513978 | 719.5233 | 0.0093 | PG O-33:1 | C39H77O9P | [M-H]- |
| 795.422504 | 795.4173 | 0.0052 | DGDG 26:6 | C41H64O15 | [M-H]- |

| | | | | | |
|---|---|---|---|---|---|
| 894.516593 | 894.5291 | 0.0125 | PS 45:11 | C51H78NO10P | [M-H]- |
| 883.525635 | 883.5342 | 0.0086 | PI 38:5 | C47H81O13P | [M-H]- |
| 889.481516 | 889.4849 | 0.0034 | PIP 32:0 | C41H80O16P2 | [M-H]- |
| 764.481702 | 764.4872 | 0.0055 | PS O-36:6 | C42H72NO9P | [M-H]- |
| 693.383886 | 693.3889 | 0.005 | SQDG 25:1 | C34H62O12S | [M-H]- |
| 861.483398 | 861.4828 | 0.0006 | SQDG 38:8 | C47H74O12S | [M-H]- |
| 492.210276 | 492.2368 | 0.0265 | LPS 16:2 | C22H40NO9P | [M-H]- |
| 1151.63797 | 1151.6547 | 0.0167 | PIP2 45:0 | C54H107O19P3 | [M-H]- |
| 1003.4102 | 1003.4356 | 0.0254 | PIP2 35:4 | C44H79O19P3 | [M-H]- |

*Supporting Table 11: List of annotations for signals of the C12 cell line identified as statistically different (ANOVA multiple-sample test, p-value ≤ 0.01) between the naïve and LPS-stimulated state (biological triplicates, each consisting of 22,500 individual mass spectra of multiple individual cells) in positive-ion mode from LIPID MAPS database research with the lowest mass difference for each input mass.*

| Input mass | Matched mass | Δ *m/z* | Name | Formula | Ion |
|---|---|---|---|---|---|
| 795.552627 | 795.5546 | 0.0019 | PG O-39:5 | C45H81O9P | [M-H]- |
| 793.537045 | 793.5389 | 0.0019 | PG O-39:6 | C45H79O9P | [M-H]- |
| 500.264285 | 500.2783 | 0.014 | PE O-20:4 | C25H44NO7P | [M-H]- |
| 1153.68063 | 1153.6727 | 0.0079 | PIP 52:8 | C61H104O16P2 | [M-H]- |
| 633.486769 | 633.4865 | 0.0003 | PA O-32:0 | C35H71O7P | [M-H]- |
| 661.464599 | 661.4814 | 0.0168 | PA 33:0 | C36H71O8P | [M-H]- |
| 1181.70342 | 1181.704 | 0.0006 | PIP 54:8 | C63H108O16P2 | [M-H]- |
| 658.445391 | 658.4453 | 0.0001 | PE 30:2 | C35H66NO8P | [M-H]- |
| 591.348098 | 591.3456 | 0.0025 | PA O-30:7 | C33H53O7P | [M-H]- |
| 1129.66293 | 1129.6727 | 0.0098 | PIP 50:6 | C59H104O16P2 | [M-H]- |
| 506.323903 | 506.3252 | 0.0013 | PE O-20:1 | C25H50NO7P | [M-H]- |
| 1123.69175 | 1123.7196 | 0.0279 | PIP 49:2 | C58H110O16P2 | [M-H]- |
| 599.463247 | 599.4681 | 0.0049 | TG O-35:5 | C38H64O5 | [M-H]- |
| 1073.70018 | 1073.7064 | 0.0062 | PI 52:8 | C61H103O13P | [M-H]- |
| 1157.72813 | 1157.704 | 0.0241 | PIP 52:6 | C61H108O16P2 | [M-H]- |
| 781.537482 | 781.5389 | 0.0014 | PG O-38:5 | C44H79O9P | [M-H]- |
| 824.581892 | 824.5811 | 0.0008 | PS O-40:4 | C46H84NO9P | [M-H]- |
| 1155.68856 | 1155.6883 | 0.0002 | PIP 52:7 | C61H106O16P2 | [M-H]- |
| 1179.68863 | 1179.6883 | 0.0003 | PIP 54:9 | C63H106O16P2 | [M-H]- |
| 546.245641 | 546.2838 | 0.0381 | PS O-20:3 | C26H46NO9P | [M-H]- |
| 805.384607 | 805.391 | 0.0064 | PIP 26:0 | C35H68O16P2 | [M-H]- |
| 1074.70418 | 1074.7169 | 0.0127 | PS 58:12 | C64H102NO10P | [M-H]- |
| 602.381903 | 602.3827 | 0.0008 | PE 26:2 | C31H58NO8P | [M-H]- |
| 1153.67459 | 1153.6727 | 0.0019 | PIP 52:8 | C61H104O16P2 | [M-H]- |
| 1179.69618 | 1179.6883 | 0.0078 | PIP 54:9 | C63H106O16P2 | [M-H]- |
| 375.321567 | 375.3269 | 0.0053 | FA 25:3 | C25H44O2 | [M-H]- |
| 375.329501 | 375.3269 | 0.0026 | FA 25:3 | C25H44O2 | [M-H]- |
| 477.280806 | 477.2987 | 0.0179 | PA O-21:1 | C24H47O7P | [M-H]- |
| 798.603762 | 798.6018 | 0.0019 | PE 40:2 | C45H86NO8P | [M-H]- |
| 375.338725 | 375.3269 | 0.0119 | FA 25:3 | C25H44O2 | [M-H]- |

| | | | | | |
|---|---|---|---|---|---|
| 375.353711 | 375.3269 | 0.0269 | FA 25:3 | C25H44O2 | [M-H]- |
| 755.391091 | 755.386 | 0.0051 | DGDG 23:5 | C38H60O15 | [M-H]- |
| 375.376166 | 375.3269 | 0.0493 | FA 25:3 | C25H44O2 | [M-H]- |
| 375.345664 | 375.3269 | 0.0188 | FA 25:3 | C25H44O2 | [M-H]- |
| 817.390784 | 817.391 | 0.0002 | PIP 27:1 | C36H68O16P2 | [M-H]- |
| 375.361987 | 375.3269 | 0.0351 | FA 25:3 | C25H44O2 | [M-H]- |
| 375.370204 | 375.3269 | 0.0433 | FA 25:3 | C25H44O2 | [M-H]- |
| 660.424775 | 660.4246 | 0.0002 | PS O-28:2 | C34H64NO9P | [M-H]- |
| 823.571602 | 823.5706 | 0.001 | PI O-34:0 | C43H85O12P | [M-H]- |
| 783.539522 | 783.5334 | 0.0061 | PA O-44:9 | C47H77O7P | [M-H]- |
| 769.468043 | 769.4661 | 0.0019 | LPI O-32:6 | C41H71O11P | [M-H]- |
| 823.425121 | 823.4403 | 0.0152 | PI 34:7 | C43H69O13P | [M-H]- |
| 1183.7045 | 1183.7196 | 0.0152 | PIP 54:7 | C63H110O16P2 | [M-H]- |
| 793.379588 | 793.3934 | 0.0138 | PI 32:8 | C41H63O13P | [M-H]- |
| 1105.69649 | 1105.7019 | 0.0054 | SQDG 56:12 | C65H102O12S | [M-H]- |
| 863.434637 | 863.4716 | 0.037 | PI 37:8 | C46H73O13P | [M-H]- |
| 545.30646 | 545.2885 | 0.018 | PG O-21:4 | C27H47O9P | [M-H]- |
| 447.299713 | 447.2881 | 0.0116 | LPA O-20:2 | C23H45O6P | [M-H]- |
| 1183.66301 | 1183.6234 | 0.0396 | PIP2 48:5 | C57H103O19P3 | [M-H]- |
| 866.666904 | 866.6644 | 0.0025 | PE 45:3 | C50H94NO8P | [M-H]- |
| 913.454346 | 913.4849 | 0.0306 | PIP 34:2 | C43H80O16P2 | [M-H]- |
| 668.351191 | 668.3569 | 0.0057 | PS 28:5 | C34H56NO10P | [M-H]- |
| 532.339251 | 532.3409 | 0.0016 | PE O-22:2 | C27H52NO7P | [M-H]- |
| 729.384125 | 729.3889 | 0.0048 | SQDG 28:4 | C37H62O12S | [M-H]- |
| 681.382613 | 681.3773 | 0.0053 | PG 30:6 | C36H59O10P | [M-H]- |

*Supporting Table 12: List of annotations for signals of the E2 cell line identified as statistically different (ANOVA multiple-sample test, p-value ≤ 0.05) between the naïve and LPS-stimulated state (biological triplicates, each consisting of 22,500 individual mass spectra of multiple individual cells) in positive-ion mode from LIPID MAPS database research with the lowest mass difference for each input mass.*

| Input mass | Matched mass | Δ $m/z$ | Name | Formula | Ion |
|---|---|---|---|---|---|
| 642.466961 | 642.4504 | 0.0165 | PE O-30:3 | C35H66NO7P | [M-H]- |
| 835.503827 | 835.5283 | 0.0245 | PA 47:11 | C50H77O8P | [M-H]- |
| 857.488328 | 857.5186 | 0.0302 | PI 36:4 | C45H79O13P | [M-H]- |
| 909.523479 | 909.5499 | 0.0264 | PI 40:6 | C49H83O13P | [M-H]- |
| 799.500186 | 799.492 | 0.0082 | PG O-40:10 | C46H73O9P | [M-H]- |
| 457.222906 | 457.2361 | 0.0132 | PA O-20:4 | C23H39O7P | [M-H]- |
| 559.275509 | 559.2678 | 0.0077 | PG 21:4 | C27H45O10P | [M-H]- |
| 726.372779 | 726.414 | 0.0413 | PE 36:10 | C41H62NO8P | [M-H]- |
| 826.536638 | 826.5392 | 0.0026 | PE 43:9 | C48H78NO8P | [M-H]- |
| 912.660993 | 912.6699 | 0.0089 | PS 45:2 | C51H96NO10P | [M-H]- |
| 455.219622 | 455.2204 | 0.0008 | PA O-20:5 | C23H37O7P | [M-H]- |
| 636.370049 | 636.3671 | 0.003 | PE 29:6 | C34H56NO8P | [M-H]- |
| 458.238495 | 458.2313 | 0.0072 | LPE 17:4 | C22H38NO7P | [M-H]- |
| 673.517526 | 673.5178 | 0.0002 | PA O-35:1 | C38H75O7P | [M-H]- |

| | | | | | |
|---|---|---|---|---|---|
| 739.385294 | 739.3828 | 0.0025 | PI O-29:7 | C38H61O12P | [M-H]- |
| 785.498643 | 785.4974 | 0.0012 | LPI O-33:5 | C42H75O11P | [M-H]- |
| 821.494569 | 821.5127 | 0.0181 | PA 46:11 | C49H75O8P | [M-H]- |
| 499.246323 | 499.2466 | 0.0003 | PA 22:4 | C25H41O8P | [M-H]- |

### References


(1) Palmer, A.; Phapale, P.; Chernyavsky, I.; Lavigne, R.; Fay, D.; Tarasov, A.; Kovalev, V.; Fuchser, J.; Nikolenko, S.; Pineau, C.; Becker, M.; Alexandrov, T. *Nature Methods* **2017**, DOI: 10.1038/nmeth.4072.

(2) Aimo, L.; Liechti, R.; Hyka-Nouspikel, N.; Niknejad, A.; Gleizes, A.; Götz, L.; Kuznetsov, D.; David, F. P. A.; van der Goot, F. Gisou; Riezman, H.; Bougueleret, L.; Xenarios, I.; Bridge, A. *Bioinformatics* **2015**, DOI: 10.1093/bioinformatics/btv285.

(3) Tsai, Y.-H.; Bhandari, D. R.; Garrett, T. J.; Carter, C. S.; Spengler, B.; Yost, R. A. *Proteomics* **2016**, DOI: 10.1002/pmic.201500536.

(4) Yang, T.; Gao, D.; Jin, F.; Jiang, Y.; Liu, H. *Rapid communications in mass spectrometry : RCM* **2016**, DOI: 10.1002/rcm.7628.

(5) Goodwin, R. J. A. *Journal of Proteomics* **2012**, DOI: 10.1016/j.jprot.2012.04.012.
